# Single-cell transcriptome profiling of an adult human cell atlas of 15 major organs

**DOI:** 10.1101/2020.03.18.996975

**Authors:** Shuai He, Lin-He Wang, Yang Liu, Yi-Qi Li, Hai-Tian Chen, Jing-Hong Xu, Wan Peng, Guo-Wang Lin, Pan-Pan Wei, Bo Li, Xiaojun Xia, Dan Wang, Jin-Xin Bei, Xiaoshun He, Zhiyong Guo

**Author notes:** Email addresses of authors: Shuai He,; Lin-He Wang,; Yang Liu,; Yi-Qi Li,; Hai-Tian Chen,; Jing-Hong Xu,; Wan Peng,; Guo-Wang Lin,; Pan-Pan Wei,; Bo Li,; Xiaojun Xia,; Dan Wang,; Jin-Xin Bei,; Xiaoshun He,; Zhiyong Guo. These authors contributed equally (Shuai He, Lin-He Wang, Yang Liu). Correspondence should be addressed to (JXB), or (XH), or (ZG).

## Abstract

**Background:** As core units of organ tissues, cells of various types play their harmonious rhythms to maintain the homeostasis of the human body. It is essential to identify the characteristics of cells in human organs and their regulatory networks for understanding the biological mechanisms related to health and disease. However, a systematic and comprehensive single-cell transcriptional profile across multiple organs of a normal human adult is missing.

**Results:** We perform single-cell transcriptomes of 84,363 cells derived from 15 tissue organs of one adult donor and generate an adult human cell atlas. The adult human cell atlas depicts 252 subtypes of cells, including major cell types such as T, B, myeloid, epithelial, and stromal cells, as well as novel *COCH^+^* fibroblasts and FibSmo cells, each of which is distinguished by multiple marker genes and transcriptional profiles. These collectively contribute to the heterogeneity of major human organs. Moreover, T cell and B cell receptor repertoire comparisons and trajectory analyses reveal direct clonal sharing of T and B cells with various developmental states among different tissues. Furthermore, novel cell markers, transcription factors and ligand-receptor pairs are identified with potential functional regulations in maintaining the homeostasis of human cells among tissues.

**Conclusions:** The adult human cell atlas reveals the inter- and intra-organ heterogeneity of cell characteristics and provides a useful resource in uncovering key events during the development of human diseases in the context of the heterogeneity of cells and organs.

## INTRODUCTION

The human body consists of multiple organs, where multiple types of cells are the core units of structure and function. Like instruments from different families in a symphony orchestra, cells and organs play their harmonious rhythms to maintain the homeostasis of the human body. Yet, perturbations in the homeostasis leads to various pathological conditions. Therefore, it is essential to identify characteristics of the cells in human organs and their regulatory networks for understanding the biological mechanisms related to health and disease.

Recent technological innovations in transcriptional profiling using single-cell RNA sequencing (scRNA-seq) have provided a promising strategy to quantify gene expression at the genome-wide level in thousands of individual cells simultaneously[1–3]. This has expanded our knowledge regarding cellular heterogeneity and networks, as well as our understanding in developments in human tissues and organs at the single-cell resolution[4–11]. Previous studies have demonstrated the cell composition for many human and mice tissues, including the brain[12], kidneys[13], lungs[14], and skin[15]. The strategy also empowers the identification of novel cell types. Cells marked by cystic fibrosis transmembrane conductance regulator (*CFTR*) were identified in the lungs of human and mouse, and were able to regulate luminal pH that was implicated in the pathogenesis of cystic fibrosis[16]. Non-genetic cellular heterogeneity has been revealed in hematopoietic progenitor cells and keratinocytes, which play important roles in maintaining hematopoiesis[17] and compartmentalizing crucial molecular activities in human epidermis[15], respectively. For the development of human embryos, transcriptome analyses of about 70,000 single cells from the first-trimester’s placenta with matched maternal blood and decidual cells uncover the cellular organization of decidua and placenta, as well as distinctive immunomodulatory and chemokine profiles of decidual natural killer (NK) cells[18]. In addition, the single-cell transcriptional profiles of embryonic and adult organs in mice have been reported, which reveal the landscape of organogenesis and the cellular heterogeneity in organs[8, 9, 19]. A very recent study on major human cell types using multiple organs from different donors revealed the genetic regulation for fetal-to-adult cell-type transitions and genetic conservation in mammalian cells[20]. However, a systematic and comprehensive single-cell transcriptional profile of multiple organs from a normal human adult has been pending. Previous studies with scRNA-seq on human samples were mostly restricted to a few specific organs with disease conditions and they did not attempt to characterize the heterogeneity and connections among multiple organs in a same individual.

Here, we aimed to investigate the transcriptional heterogeneity and interactions of cells from an adult human’s organs at the single-cell resolution level. Using scRNA-seq, we profiled the transcriptomes of more than 84,000 cells of 15 organs from one individual donor. Comprehensive comparisons within and across tissues for distinct cell types were performed to reveal the inter-cellular complexity of gene profiles, active transcription factors and potential biological functions, as well as potential inter-cell connections. The resulting high-resolution adult human cell atlas (AHCA) provides a global view of various cell populations and connections in the human body, and is also a useful resource to investigate the biology of normal human cells and the development of diseases affecting different organs.

## RESULTS

### Global view of single-cell RNA sequencing of 15 organ samples

Viable single cells were prepared from the tissue samples of 15 different organs of a research-consented adult donor (**Fig. 1A**). mRNA transcripts from each sample were ligated with barcoded indexes at 5’-end and reverse transcribed into cDNA, using GemCode technology (10x Genomics, USA). cDNA libraries including enriched fragments spanning the full-length V(D)J segments of T cell receptors (TCR) or B cell receptors (BCR), and 5’-end fragments for gene expression were separately constructed, which were subsequently subjected for high-throughput sequencing. On average, we obtained more than 400 million sequencing reads for each organ sample, which resulted in a median sequencing saturation (covering the fraction of library complexity) of 88% (61.6%-97%) for each sample (**Additional file 1: Figure S1A** and **Additional file 2: Table S1**). After primary quality control (QC) filters, 91,393 cells were identified (**Additional file 1: Figure S1B** and **Additional file 2: Table S2**). Higher number of UMIs and more transcribed genes were observed in the skin and trachea samples (with median UMIs of 4,022.5 and 4,100.5 and genes of 1,528 and 1,653, respectively) compared with the other organs **(Additional file 1: Figure S1C, D**, and **Additional file 2: Table S2**). We obtained 66,225 sequencing read pairs for each cell and 6,093 cells for each organ on average (**Additional file 2: Table S1, S2**), with more than 2.4 and 3.5 times deeper sequencing of median genes and UMIs than a recent study[20]. The cells in each organ were classified using unsupervised clustering, and cell types were assigned based on canonical marker genes (**Additional file 2: Table S3**). Next, visualization of the cells by t-distributed stochastic neighbor embedding (t-SNE) revealed multiple subpopulations of cells in each organ, with the numbers of clusters ranging from 9 in the blood to 25 in the skin (**Additional file 1: Figure S2, Figure S2 continued**, and **Additional file 2: Table S4-S18**). Clusters due to cell doublets were identified and excluded for each organ, which resulted in a total of 84,363 cells for the downstream analyses (**Additional file 2: Table S2**).

**Figure 1.**
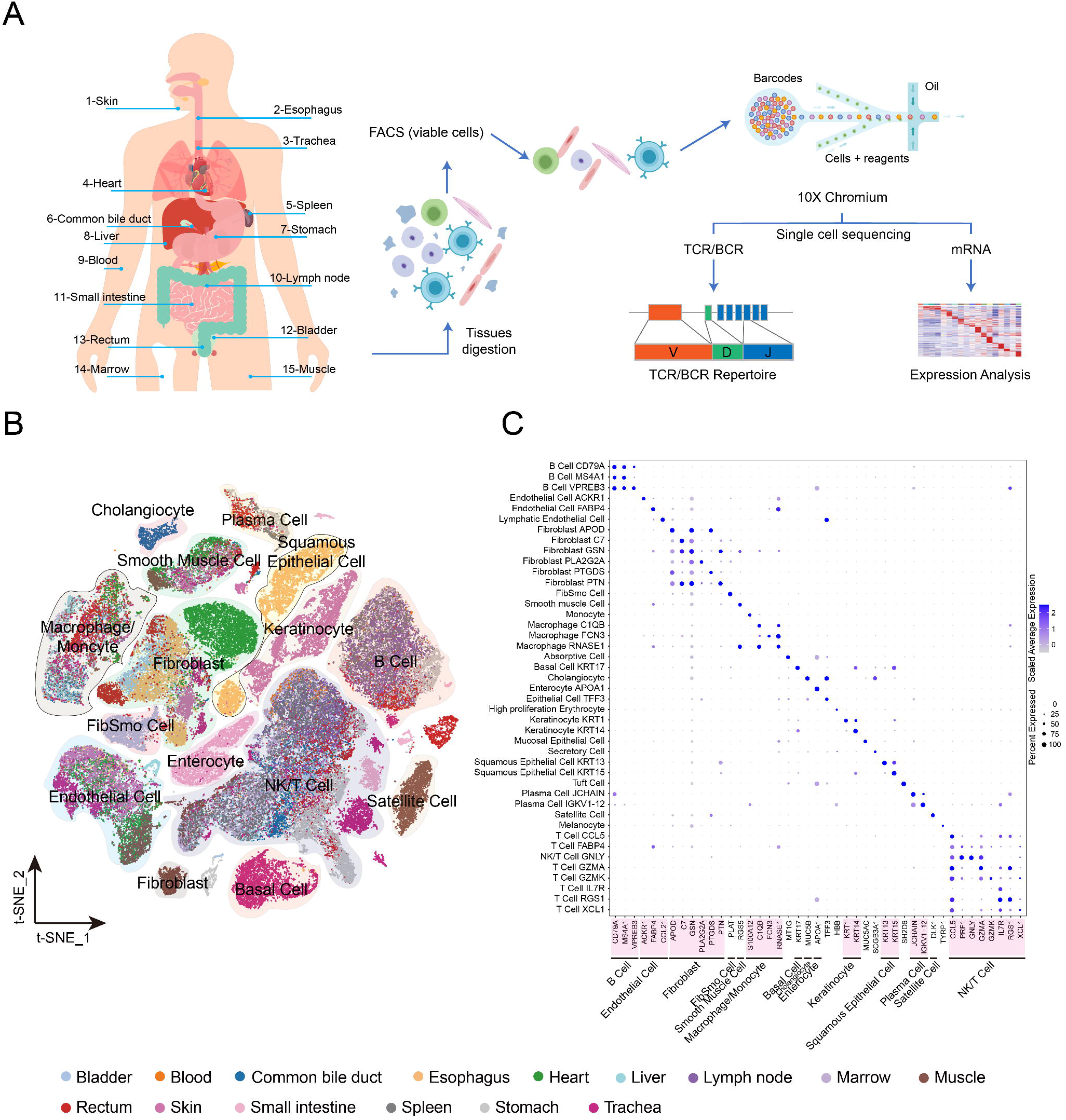
Overview of single-cell RNA sequencing of 15 organ tissues from a male adult donor. **A.** An experiment schematic diagram highlighting the sites of the organs for tissue collection and sample processing. Live cells were collected using flow cytometry sorting (FACS) and subjected for cell barcoding. cDNA libraries for TCR, BCR, and 5’-mRNA expression were constructed independently, followed by high throughput sequencing and downstream analyses. **B.** t-SNE visualization of all cells (84,363) in organs. Each dot represents one cell, with colors coded according to the origin of organ. Labeled cell types are the predominant cell types in each cluster. **C.** Dot plots showing the most highly expressed marker genes (x axis) of major cell types (y axis) in **Figure 1B**. The depth of the color from white to blue and the size of the dot represents the average expression from low to high and the percent of cells expressing the gene.

With transcriptional profiles of such large number of cells, we identified some rare and novel cell populations. A group of Langerhans cells were identified in the skin sample (1% of all skin cells) with specific expression of *CD207* and *CD1A*[21]. An even smaller group of 26 sweat gland epithelial cells (0.31%) were also identified in the skin sample, which had specific expression of *DCD, SCGB2A2, KRT19, MUCL1*, and *PIP* genes (**Additional file 1: Figure S3A** and **Additional file 2: Table S14**). A novel group of fibroblasts (0.43%) with exclusive expression of *COCH* were identified in the skin (**Additional file 1: Figure S2 continued**, **Figure S3A**, and **Additional file 2: Table S14**). Of note, another novel group of cells were assigned as FibSmo with a co-expression of *MMP2* and *ACTA2*, which are marker genes for fibroblasts and smooth muscle cells, and were identified with higher proportions in the rectum (6.66%), bladder (17.59%), and heart (7.75%) than in the other tissue organs (**Additional file 1: Figure S2, Figure S2 continued, Figure S3B**, and **Additional file 2: Table S4, S13**). Moreover, in contrast to the broad distribution of FibSmo cells in multiple organs, *COCH^+^* fibroblasts were identified in limited organs with low abundance, while sweat gland epithelial cells were found specifically in the skin (**Additional file 1: Figure S3B**). Furthermore, the presence of sweat gland epithelial cells, *COCH^+^* fibroblasts, and FibSmo cells were confirmed in multiple tissue samples from independent donors using existing datasets and multiplex immunofluorescence staining assays (**Additional file 1: Figure S4-S10** and **Additional file 3: Supplementary Notes**).

We combined all the 84,363 cells in the cluster analysis and identified 43 clusters in 15 organs (**Fig. 1B, Additional file 1: Figure S11** and **Additional file 2: Table S19**). We observed close clustering of cells from different organs (more than seven organs) for major cell types, including T, B, plasma, endothelial, and smooth muscle cells, as well as fibroblasts, macrophages, and monocytes (**Fig. 1B, C, Additional file 1: Figure S11, Additional file 1: Figure S12**, and **Additional file 2: Table S20**). This is consistent with the understanding that cells derived from the same lineage are widely distributed within the human body, especially circulating immune cells. Moreover, multiple clusters were further identified for several major cell types (T cells, B cells, fibroblasts, myeloid cells, and endothelial cells), reflecting their heterogeneous transcriptional profiles (**Fig. 1C, Additional file 2: Table S19**).

### The heterogeneity of T cells in developmental state and clonalities around the body

We identified a total of 20,034 T cells prevailing in the immune cells of most organ tissues (**Additional file 4: Table S21**), which is consistent with a previous finding[22]. These included 1,472 γδ and 18,292 αβ T cells. The latter were divided into CD4^+^ (7,006) and CD8^+^ (11,286) T cells according to their gene profiles and were further grouped into 11 and 21 major unsupervised clusters, respectively (**Fig. 2A, B**), including naïve/central memory T (T_N/CM_), effector memory T (T_EM_), regulatory T (T_reg_), tissue-resident memory T (T_RM_), effector T (T_h1_ for CD4^+^ and T_EFF_ for CD8^+^ T cell), intraepithelial lymphocytes (IEL) T, and mucosal-associated invariant T (MAIT) cells, based on known markers[23] (**Fig. 2C, D**). Some T_N/CM_ cells were further assigned as T_N_, and T_CM_ clusters based on their gene signatures (**Fig. 2C, D, Additional file 1: Figure S13A, B**). Both CD4^+^ and CD8^+^ T cell clusters showed a distribution pattern in an organ-specific manner (**Additional file 1: Figure S13C, D**) and each of them had differentially expressed genes (**Additional file 1: Figure S13A, B** and **Additional file 4: Table S22, S23**). An overlapping of clusters was also observed between organs (such as blood and marrow), suggesting the sharing of common T cell subtypes (**Additional file 1: Figure S13D**).

**Figure 2.**
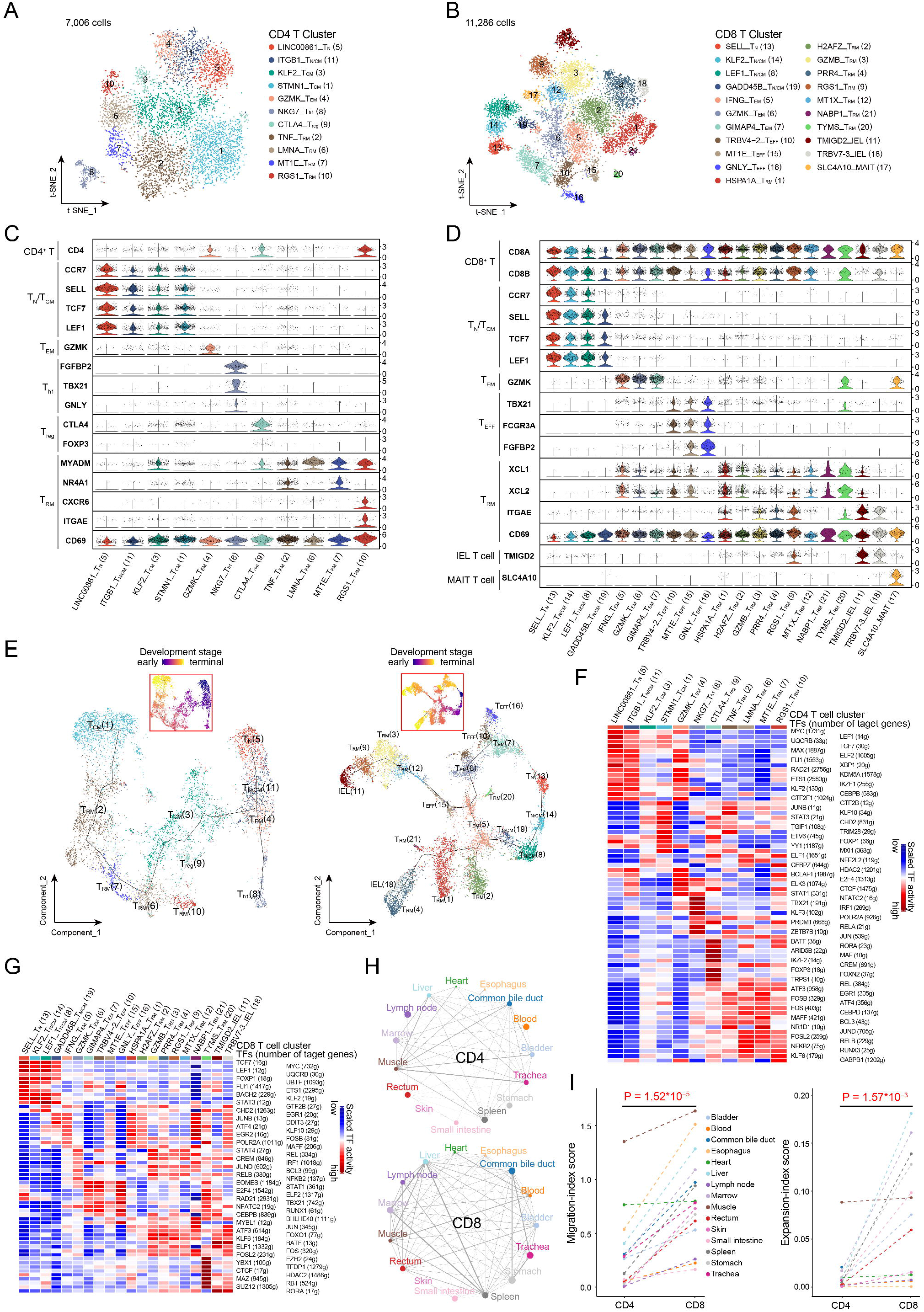
The heterogeneity, development and clonality of T cells in human organs. **A, B.** t-SNE plots of 7,006 CD4^+^ (**A**, 11 clusters) and 11,256 CD8^+^ (**B**, 21 clusters) T cells from 15 organ tissues. Each dot represents one cell. Each color-coded region represents one cell cluster, which is indicated on the right. **C, D.** Violin plots showing the normalized expression of marker genes for each CD4^+^ (**C**) and CD8^+^ (**D**) T cell cluster as indicated at the bottom. For each panel, the y-axis shows the normalized expression level for a marker gene as indicated on the left. Marker genes were also grouped according to functional cell types. **E.** Pseudo-time trajectory analysis of all CD4^+^ (left panel) and CD8^+^ T cells (right panel) with high variable genes. Each dot represents one cell and is colored according to their cluster above: **A** for CD4^+^ and **B** for CD8^+^. The inlet t-SNE plot shows each cell with a pseudo-time score from dark blue to yellow, indicating early and terminal states, respectively. **F,G.** Heat maps of the activation scores of each T cell cluster for expression regulated by transcription factors (TFs). T cell clusters are indicated on top, and the scores were estimated using SCENIC analysis. Only shows the top 15 TFs for CD4^+^ T cells (**F**) and the top 10 for CD8^+^ T cells (**G**), with the highest difference in expression regulation estimates between each cluster and all other cells, under a Wilcoxon rank-sum test. **H.** Sharing intensity of TCR clones in CD4^+^ (top panel) and CD8^+^ (bottom panel) T cells between different organ samples. Each line represents a sharing of TCR between two organs at the ends and the thickness of the line represents a migration-index score between paired organs calculated by STARTRAC. The sizes of the dots are shown according to the logarithm to the base 2 of the size of T cell clones in organs with different colors. **I.** Migration- (left panel) and expansion-index (right panel) scores of CD4^+^ and CD8^+^ T cells of each tissue calculated and compared using STARTRAC with a paired Student’s t-test.

To better understand the developmental state of T cells, we performed trajectory analyses of CD4^+^ and CD8^+^ T cells. We observed that the trajectory trees rooted from T_N_ cells, sprouting into T_CM_, T_h1_, and T_RM_ branches for CD4^+^ T cells and T_N/CM_, T_EFF_, IEL, and T_RM_ branches for CD8^+^ T cells (**Fig. 2E, Additional file 1: Figure S13E, F**). T_RM_ cells and IEL cells with higher pseudo-time scores were found in CD4^+^ and CD8^+^ T cells, respectively, suggesting their terminal developmental state (**Fig. 2E, Additional file 1: Figure S13E, F**). Moreover, several T_RM_ clusters at the end of other branches with mediated scores for both CD4^+^ and CD8^+^ T cells, indicating the middle developmental state of these clusters. The T_RM_ clusters with different developmental states showed an organ-specific pattern (**Additional file 1: Figure S13E, F**), while their origin from marrow or spleen was unclear due to the limited number of cells. These observations reveal the heterogeneity in the developmental states of both CD4^+^ and CD8^+^ cells in human organs.

Transcription factors (TFs) have been demonstrated as important regulators of gene expression and with ability to shape different phenotypes of T cells[24]. We therefore, performed Single-Cell Regulatory Network Inference and Clustering (SCENIC) analysis to assess TFs underlying differential gene expression in T cells. We identified well-defined and cell-subtype-specific TFs for CD4^+^ (**Fig. 2F**) and CD8^+^ T cell (**Fig. 2G**) clusters (**Additional file 4: Table S24, S25**), such as higher activity of *FOXP3*[24] and *BATF*[25] in T_reg_ cells, upregulation of *LEF1, MYC, TCF7*[26], and *KLF2*[27] in T_N_ cells and *TBX21, STAT1*, and *IRF1* in T_EFF_ cells[24, 28, 29] (**Fig. 2F, G** and **Additional file 1: Figure S13G, H**). In addition, many other poorly investigated TFs were also observed in both CD4^+^ and CD8^+^ T_RM_ cell clusters, such as the upregulation of several AP-1 dimerization partners (*FOS, JUN, JUND, FOSL2*, and *ATF3*), *REL*, and *RELB* (**Fig. 2F, G** and **Additional file 1: Figure S13G, H**). Collectively, these results indicate that the combinations of multiple TFs regulate T cell development to maintain the heterogeneous states of CD4^+^ and CD8^+^ T cells.

To better investigate the clonalities and dynamic relationships among T cell subtypes across tissues, we preformed TCR clonal typing accompanied with transcriptome analysis (**Fig. 1A**). After stringent QC filters, we identified 5,183 TCR clonotypes with unique heterodimer α and β chains among 8,394 T cells (45.89% of the whole 18,292 CD4^+^ and CD8^+^ T cells), including 3,248 CD4^+^ T cells and 5,146 CD8^+^ T cells. Among them, 4,645 cells (2,906 CD4^+^ and 1,739 CD8^+^ T cells) had a unique TCR clonotype for each, while the remaining 3,749 cells (342 CD4^+^ and 3,407 CD8^+^ T cells) shared two or more of the 538 TCR clonotypes (**Additional file 4: Table S26, S27**). We observed similar numbers of V and J segments for the TCR α chain in both CD4^+^ and CD8^+^ T cells, both of which shared 60% and 40% of the top 10 frequent V and J segments, respectively (**Additional file 1: Figure S14A, B**). By contrast, the diversity of the V segment was much higher than that of the J segment for β chain in both CD4^+^ and CD8^+^ T cells, which shared 40 % and 80% of the top 10 frequent V and J segments, respectively (**Additional file 1: Figure S14A, B**). Although more CD8^+^ T cells were detected than CD4^+^ T cells among all tissues, no significant difference in clone sizes (the number of unique clonotypes) was observed between the two cell populations (**Additional file 1: Figure S14C**). Singular cells of unique TCR clonotypes were prevalent for CD4^+^ T cells among tissues, except that a higher proportion of multiple cells with identical TCR clonotypes or clonal expansions were observed in the muscle, common bile duct, and marrow (**Additional file 1: Figure S14D** top panel). Clonal expansions were much commonly found for CD8^+^ T cells in all tissues (**Additional file 1: Figure S14D** bottom panel).

To investigate the clonotype distribution of T cells across tissues, we evaluated the ability of sharing TCR for each tissue with others. We observed a more intensive and broader sharing of TCR clonotype for CD8^+^ than CD4^+^ T cells across tissues (**Fig. 2H**). A higher migration capacity, reflected in the migration-index score of tissues, was found in CD8^+^ T cells than in CD4^+^ T cells (**Fig. 2I** left panel). Moreover, higher expansion ability but lower diversity was observed in CD8^+^ T cells in each tissue compared with CD4^+^ T cells (**Fig. 2I** right panel, **Additional file 1: Figure S14E**). Cells with clonal expansion had considerable proportions in T_EM_, T_EFF_, T_RM_, and IEL clusters of CD8^+^ T cells (**Additional file 1: Figure S14F** bottom panel), and T_h1_ and RGS1_T_RM_ of CD4^+^ T cells (**Additional file 1: Figure S14F** top panel). We further evaluated the clonal expansion and transition (clonotype sharing ability of each subpopulation) of each T cell cluster, which revealed significantly stronger expansion and transition abilities of CD8^+^ compared with CD4^+^ T cells (**Additional file 1: Figure S14G**, **H**). This is consistent with the stronger sharing links of clonotypes between subtypes of CD8^+^ than CD4^+^ T cells (**Fig. 2H, Additional file 1: Figure S14I, J**). Moreover, we observed the clonal expansion of certain CD8^+^ T cells (T_EFF_, T_RM_, T_EM_, and IEL T cells) distributed across multiple tissues with different developmental states (**Additional file 1: Figure S15A**), suggesting that most of these cells recognizing the same antigens propagate and migrate more intensively than T cells of other types. In addition, we observed consistent results of TCR diversity using β chains (**Additional file 1: Figure S15B-D**). These observations based on TCR tracing suggest the widespread of diverse T cells across the human body through clonal expansion and transition.

### The heterogeneity of B cells and plasma cells

Cluster analysis revealed 14 distinct cell clusters among 10,100 B and plasma cells from 11 organ samples, including nine B (*CD20*) and six plasma cell (*SDC1*) clusters (**Fig. 3A**). We observed the predominance of B cells over plasma cells in all tested organs except for the esophagus and rectum (**Fig. 3B**). Differential gene expression (DEG) analysis revealed that B cells exhibited distinct gene profiles from plasma cells (**Additional file 5: Table S28** and **Additional file 1: Figure S16A**). Although *CD27* is a canonical marker of memory B cells[30], we observed a low transcription level of *CD27* in memory B cells, but a higher level in plasma cells (**Fig. 3C**). *TCL1A* was significantly expressed in two naïve B cell clusters with distinct gene expression profiles, TCL1A_ly_naive_B from the lymph nodes and TCL1A_naive_B from multiple tissues, compared with the other B and plasma cell clusters (**Fig. 3A, Additional file 1: Figure S16A**). Moreover, *TCL1A* was exclusively expressed in non-CD27-expressing B cell clusters and had a significantly reverse correlation with *CD27* transcription (Person’s R = - 0.84, *P* = 0; **Fig. 3C, Additional file 1: Figure S16B**). Given that *TCL1A* has been reported as an important gene in B cell lymphomas[31], these findings suggest that *TCL1A* might be a novel marker of naïve B cells.

**Figure 3.**
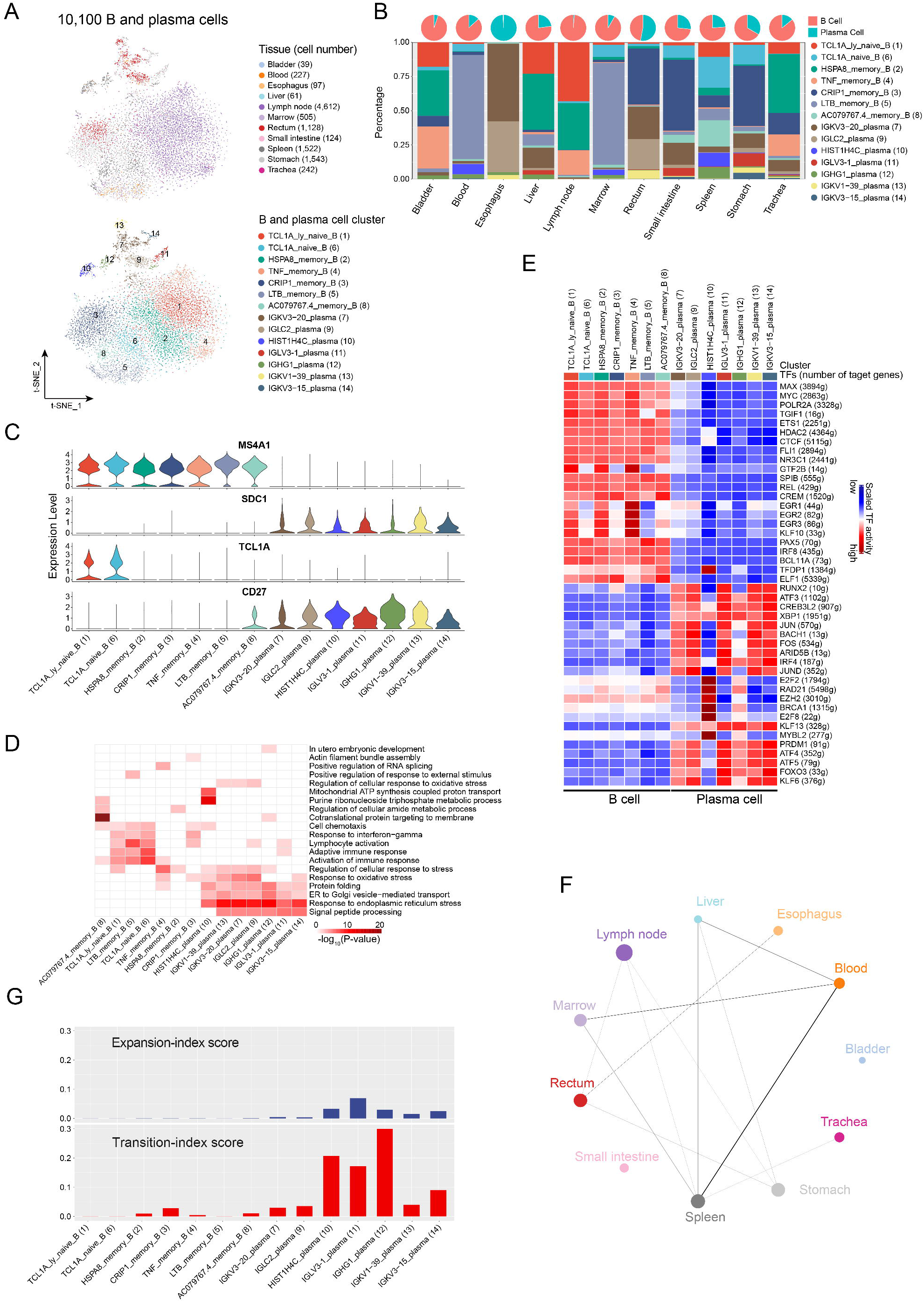
The heterogeneity and clonality of B cells in human organs. **A.** t-SNE plots showing 14 clusters (10,100 cells) of B and plasma cells. Each dot represents a cell, colored according to the origin of tissue (top panel) and cell subtype (bottom panel). **B.** Distribution of B and plasma cells in each organ. Pie charts on top illustrate the proportions of B and plasma cells in each organ. The stacked bars represent the percentage of each cluster in the indicated organ. **C.** Violin plots of the normalized expression of maker genes for B (*MS4A1*), plasma cells (*SDC1*), naïve B cell (*TCL1A*), and memory B cells (*CD27*). For each panel, the y-axis shows the normalized expression level for a marker gene as indicated on the title, and the x-axis indicates cell clusters. **D.** Gene Ontology enrichment analysis results of B and plasma cell clusters. Cell clusters as indicated at the bottom are colored according to their −log™*P-*values in columns. Only the top 20 significant GO terms (*P*-value < 0.05) are shown in rows. **E.** Heat map of the activation scores of each B and plasma cell cluster for expression regulated by transcription factors (TFs). Cell clusters are indicated on top, and the scores were estimated using SCENIC analysis. It shows the top 10 TFs with the highest difference in expression regulation estimates between each cluster and all other cells, tested with a Wilcoxon ranksum test. **F.** Sharing intensity of BCR clones between different organs. Each line represents a sharing of BCR between two organs at the ends and the thickness of the line represents a migration-index score between paired organs calculated using STARTRAC. The size of the dot is shown as the logarithm to the base 2 of the size of B and plasma cell clones in each organ. **G.** Expansion- (top panel) and transition-index (bottom panel) scores of each B and plasma cell cluster calculated using STARTRAC.

B cells are professional antigen-presenting cells (APCs) with high expression of *CIITA* and *MHC* class II genes, which are silenced during the differentiation to plasma cells[32]. We examined the antigen-presenting ability of B and plasma cell clusters, using the antigenpresenting score (APS) based on the expression of signature genes related to antigenpresenting (See **Methods**). Interestingly, plasma cells had a much lower APS for extracellular antigens compared with B cell clusters (**Additional file 1: Figure S16C**), while they had a much higher APS for intercellular antigens than B cells, suggesting the different abilities to present antigens between the two cell types. We also investigated the potential biological function of different cell clusters using Gene Ontology enrichment (GO) analysis. Potential biological functions of B cells were found to be enriched in immune response (for example, ‘*responding to activation of immune*’) and that of plasma cells were associated with protein synthesis (including ‘*signal peptide processing*’, ‘*ER to Golgi vesicle-mediated transport*’, and ‘*protein folding*’; **Fig. 3D**). In addition, gene set enrichment analysis (GSEA) showed that TCL1A_naive_B cells were enriched in ‘*oxidative phosphorylation*’ and ‘*fatty acid metabolism*’ pathways (**Additional file 1: Figure S16D**). These observations suggest distinct biological functions of B and plasma cells, although plasma cells were derived from B cells.

TFs play a critical role in the differentiation of B cells to plasma cells, engaging B cells with effector or memory functions[33]. Single-cell regulatory network inference and clustering (SCENIC) analysis revealed that TFs exhibited similar activities in B and plasma cells but with distinct patterns between the two populations (**Fig. 3E, Additional file 5: Table S29**). TFs with higher activity were found in B cells, including *MYC*[34] and *REL*[35], which have been known to modulate B cell development, as well as other TFs that have not been characterized in B cells, such as EGF receptors (*EGFR1/2/3*) and *TGIF1*. Likewise, TFs enriched in plasma cells included *PRDM1, XBP1, FOS*, and *IRF4*, which play important roles in the development of plasma cells[33]. Moreover, many TFs with unknown roles were found, including *ATF3, ATF4* and *ATF5*. Consistently, we observed higher activity of these TFs in an independent human cell landscape (HCL) dataset published recently[20], including *MYC, IRF8*, and *REL* in B cell clusters and *XBP1, PRDM1*, and *CREB3L2* in plasma cell clusters (**Additional file 1: Figure S16E, F**, and **Additional file 4: Table S30, S31**). Taken together, these results suggest that various TFs might regulate the development of B cells into plasma cells.

To explore the clonalities of B and plasma cell clusters across organ tissues, we performed a single-cell BCR sequencing analysis. After stringent QC filters, 6,741 out of 10,100 cells were assigned to 6,480 clonotypes, among which 6,330 clonotypes were presented by singular cells and 150 by multiple cells (**Additional file 4: Table S32**). We observed various usage of V and J gene segments for both heavy and light chains of immunoglobulin genes, with a preferred usage of some particular variable segments (**Additional file 1: Figure S16G-I**). Unlike T cells, clonal diversity was common for B cells among all the organs, while clonal expansion of B cells was limited and restricted to the spleen, rectum, and stomach (**Additional file 1: Figure S16J**). Moreover, less sharing of BCR clonotypes between B and plasma cells was observed across organs compared with T cells (**Fig. 2H** and **Fig. 3F**). In addition, BCR analysis showed lower expansion and transition abilities of B cells compared with plasma cells and T cells (**Fig. 3G, Additional file 1: Figure S14G**, and **Additional file 1: Figure S16K, L**), which might be due to an insufficient representation of the richness of the diverse B cell repertoire with the limited number of clonal B cells detected.

### The heterogeneity of myeloid cells

We obtained 5,587 myeloid cells from 15 organ tissues, which were grouped into 18 distinct clusters (**Fig. 4A, B**). Based on the differential expressions of marker genes, we further identified seven monocyte clusters, eight macrophage clusters, and three dendritic cell (DC) clusters (**Fig. 4A, B**, and **Additional file 1: Figure S17A**). The hierarchical cluster analysis revealed that all classical monocytes were closely related as a branch node, as were non-classical monocytes and intermediate monocytes (**Fig. 4C**). All macrophages were grouped together and separated from monocytes, except that one subset of macrophages (C3: SDC3_Mac) were closely related to intermediate and non-classical monocytes (**Fig. 4C**). DEG analysis showed that each myeloid cell cluster had a specific gene signature, suggesting intercell heterogeneity among monocytes, macrophages, and DC cells (**Fig. 4D, Additional file 6: Table S33**). All clusters contained cells from multiple tissues, except for FN1_Intermediate_Mon (C4) and CCL20_Intermediate_Mon (C10) in the rectum and Langerhans clusters (C15: Langerhans) in the skin, suggesting similar transcriptional profiles and origins of these myeloid cells (**Fig. 4A, B**). Monocytes showed a predominance in 11 of the 15 tested tissues, except for the esophagus, heart, lymph nodes, and skin, where macrophages and DC cells accounted for more than 50% of all myeloid cells (**Additional file 1: S17A**).

**Figure 4.**
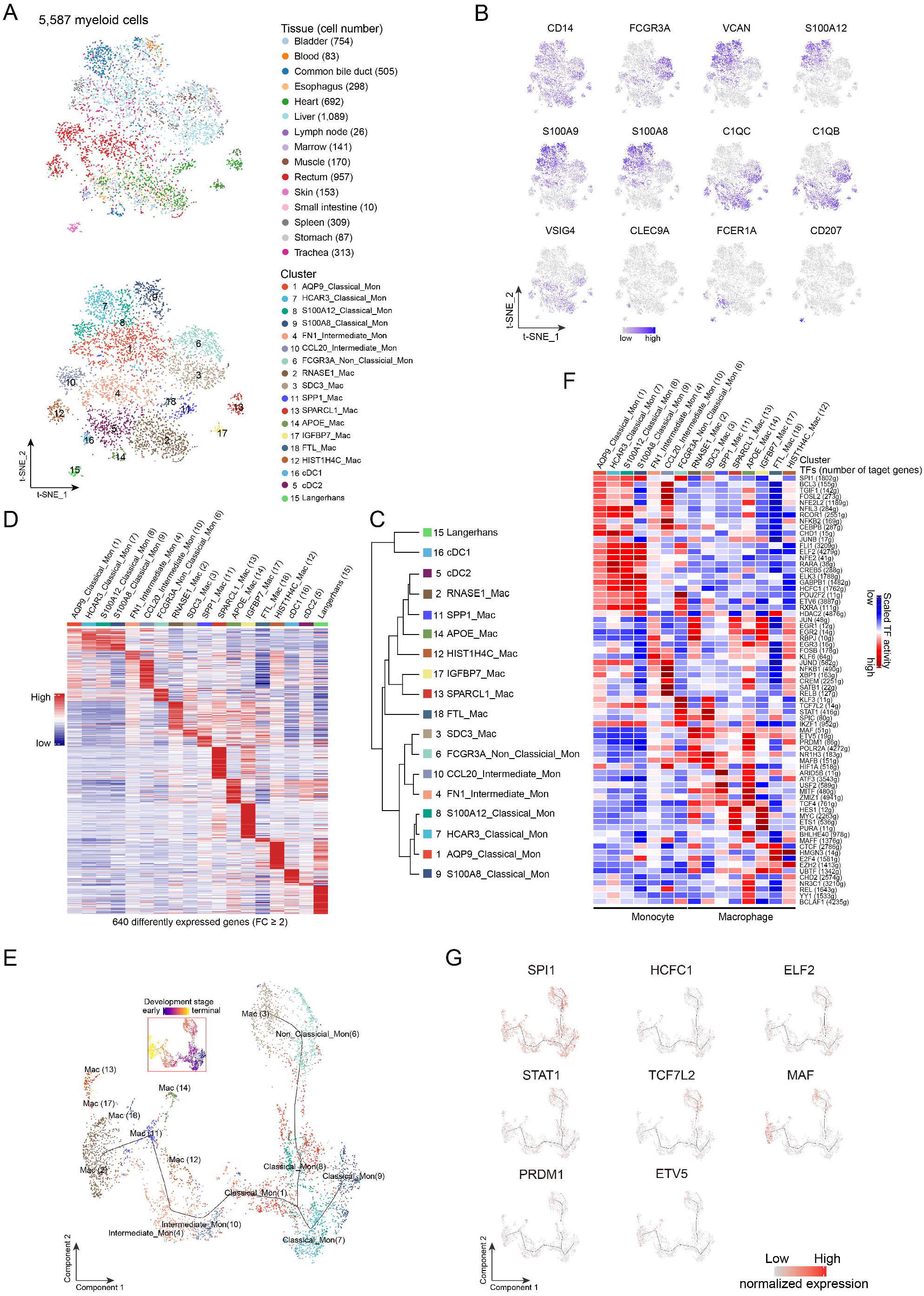
Heterogeneity and developmental stages of myeloid cells. **A.** t-SNE plots of 5,587 myeloid cells. Each dot represents one cell, colored according to their tissue origins (top panel) or cell clusters (bottom panel) as indicated on the right. **B.** t-SNE plots of the normalized expression of marker genes for monocytes (*S100A8/9/12 and VCAN*) and macrophages (pan-marker: *C1QC, C1QB*, and *VSIG4*), cDC1 (*CLEC9A*), cDC2 (*FCER1A*), Langerhans (*CD207*) as well as subpopulation-specific genes (*CD14* and *FCGR3A*). Each dot represents one cell, with a color from grey to blue representing the expression level from low to high. **C.** Dendrogram of 18 clusters based on their normalized mean expression values (correlation distance metric, complete linkage). Only genes with ln(fold-change) above 0.25, *p.adjust* < 0.05 and pct.1 ≥ 0.2 in each cluster were included in the calculations. **D.** Heat map showing the expression profiles of each myeloid cell cluster as indicated on top. The expression of 640 genes in each cell cluster with FC ≥ 2 and *p.adjust* < 0.05 are shown as lines, colored from blue to red according to the expression from low to high. **E.** Pseudo-time trajectory analysis of all myeloid cells with high variable genes. Each dot represents one cell and is colored according to their clustering in **A**. The inlet t-SNE plot shows each cell with a pseudo-time score from dark blue to yellow, indicating early and terminal states, respectively. **F.** Heat map of the activation scores of each monocyte and macrophage subtype for gene expression regulated by transcription factors (TFs). Cell clusters are indicated on top, and the scores were estimated using SCENIC analysis. Only the top 10 TFs are shown with the highest difference in expression regulation estimates between each cluster and all other cells, tested with a Wilcoxon rank-sum test. **G.** Plots showing the normalized expression of representative TFs in **F** along the pseudo-time trajectory maps corresponding to **E**. Each dot in one plot shows the expression of the indicated gene in the plot, colored from grey to red, indicating low and high expression, respectively. *SPIP1, HCFC1, ELF2* for classical monocytes; enhanced expression of *STAT1* and *TCF7L2* for non-classical monocytes and SDC3_Mac (3); *MAF, PRDM1* and *EVT5* for most not-classical monocytes.

Considering two potential origins of macrophages in multiple tissues from circulating monocytes and embryonic progenitor cells[36–38], we examined the connection between macrophages and monocytes. First, we observed the coexistence of macrophages and monocytes in the same organs (**Fig. 4A, B**, and **Additional file 1: Figure S17A**). Second, trajectory analysis revealed that the classical monocytes had initial state at the root and sprouted into branches with more developed states of monocytes and then macrophages either directly or via two intermediate monocytes (**Fig. 4E, Additional file 1: Figure S17B, C**). Notably, high expression of proliferation marker genes including *MKI67* and *PCNA* were exclusively detected in the HISTIH4C_Mac macrophages (C12) in the bladder, esophagus, heart, and rectum (each with more than five of the cells; **Additional file 1: Figure S17D**). Gene set variation analysis (GSVA) also revealed that most of the macrophage clusters had a higher enrichment score of *MYC and E2F* target pathways (**Additional file 1: Figure S17E**). We further performed a trajectory analysis of intestinal monocytes and macrophages derived from our and published datasets[39]. We observed a terminal state for both the embryonic and adult macrophages according to pseudotime scores (**Additional file 1: Figure S17F**). Moreover, we observed a clear differentiation trajectory of the macrophages in the adult rectum from the intermediate monocytes (CCL20_Intermediate_Mon) to the tissuemacrophages (HIST1H4C_Mac). Interestingly, the embryonic macrophages (Embryo_Mac) were found alongside the differentiation trajectory from the intermediate monocyte (FN1_Intermediate_Mon) to the tissue-macrophages (HIST1H4C_Mac), suggesting that the embryonic macrophages may contribute to the tissue-macrophages through local expansion. Consistently, a high expression of the proliferation marker gene *PCNA* was detected in the terminal macrophages (**Additional file 1: Figure S17G**). Taken together, these observations suggest that circulating monocytes might give rise to macrophages in organ tissues and that the local microenvironment may contribute to the expansion and proliferation of different embryo-derived macrophage populations, especially tissue-resident macrophages, in the bladder, esophagus, heart, and rectum.

TFs such as *SPI1* are involved in the development of monocytes to macrophages[40, 41]. However, how TFs regulate their development in the normal human body is still unclear. TF activity analysis revealed a similar activation of certain TFs among the four classical monocytes, which is consistent with their close similarity in gene signatures (**Fig. 4D, F** and **Additional file 6: Table S34**). A high activation of *SPI1* was determined in classical monocytes, non-classical monocytes and SDC3_Mac cells, suggesting its involvement in the development of these cells. We also identified several TFs with high activity in the four classical monocyte clusters, including *HCFC1, ELF2, ETV6, ELK3*, and *NFE2* (**Fig. 4F, G**). Their roles in maintaining the classical state of monocytes have yet to be investigated. Non-classical monocytes and macrophages shared several activated TFs, such as *TCF7L2, STAT1, KLF3, NR1H3*, and *SPIC*, which is consistent with their close pattern in the hierarchical cluster analysis (**Fig. 4C, F and G**). We observed that multiple poorly characterized TFs were highly activated in the intermediate, non-classical monocytes and macrophages, other than classical monocytes, including *POLR2A, MAF, MAFB, PRDM1, ETV5*, and *ATF3* (**Fig. 4C, F and G**). We also observed cluster-specific TFs for myeloid cell clusters (**Fig. 4F, Additional file 6: Table S34**). For instance, the activation of *BHLHE40* and *NR3C1* was higher in APOE_Mac cells (C14). These results suggest that these unique combinations of TFs help shape the different states of myeloid cells in the normal human body.

It has been demonstrated that myeloid cells could act as professional APCs, with the strongest antigen-presenting ability for DCs[42]. We observed various antigen-presenting abilities for extracellular antigens as reflected by APS for different myeloid clusters (**Additional file 1: Figure S17H**). Langerhans cells had a stronger antigen-presenting ability than other macrophages and monocytes (*P* < 2.2 x 10^-16^; **Additional file 1: Figure S17H**). On average, the classical monocytes had the lowest APS among myeloid cell clusters (*P* < 2.2 x 10^-16^). Interestingly, different myeloid cell clusters had a similar APS for presenting intracellular antigen (**Additional file 1: Figure S17H**). These observations support the varied roles of myeloid cells in antigen presentation.

### The similarity and heterogeneity of epithelial cells from intra- and inter-tissues

We obtained a total of 17,436 epithelial cells from nine tested organ tissues (**Fig. 5A**). DEG analysis revealed a clear and distinct pattern of gene expression among the tissue samples (**Additional file 1: Figure S18A** and **Additional file 7: Table S35**). There were 190 genes with tissue-specific expression (FC ≥ 5, pct.1 ≥ 0.2; **Additional file 7: Table S35** and **Additional file 1: Figure S18A**), indicating the heterogeneity of epithelial cells among organ tissues, which was further confirmed in the HCL dataset (**Additional file 1: Figure S18B**). GO analysis also revealed the various biological functions of epithelial cells from different tissues, among which, ‘*epithelial cell differentiation*’ and ‘*regulation of myeloid leukocyte activation*’ were common pathways enriched in the majority of epithelial cells. This suggests that the cells share common functions in the development of epithelial cells and the regulation of immune response (**Additional file 1: Figure S18C**).

**Figure 5.**
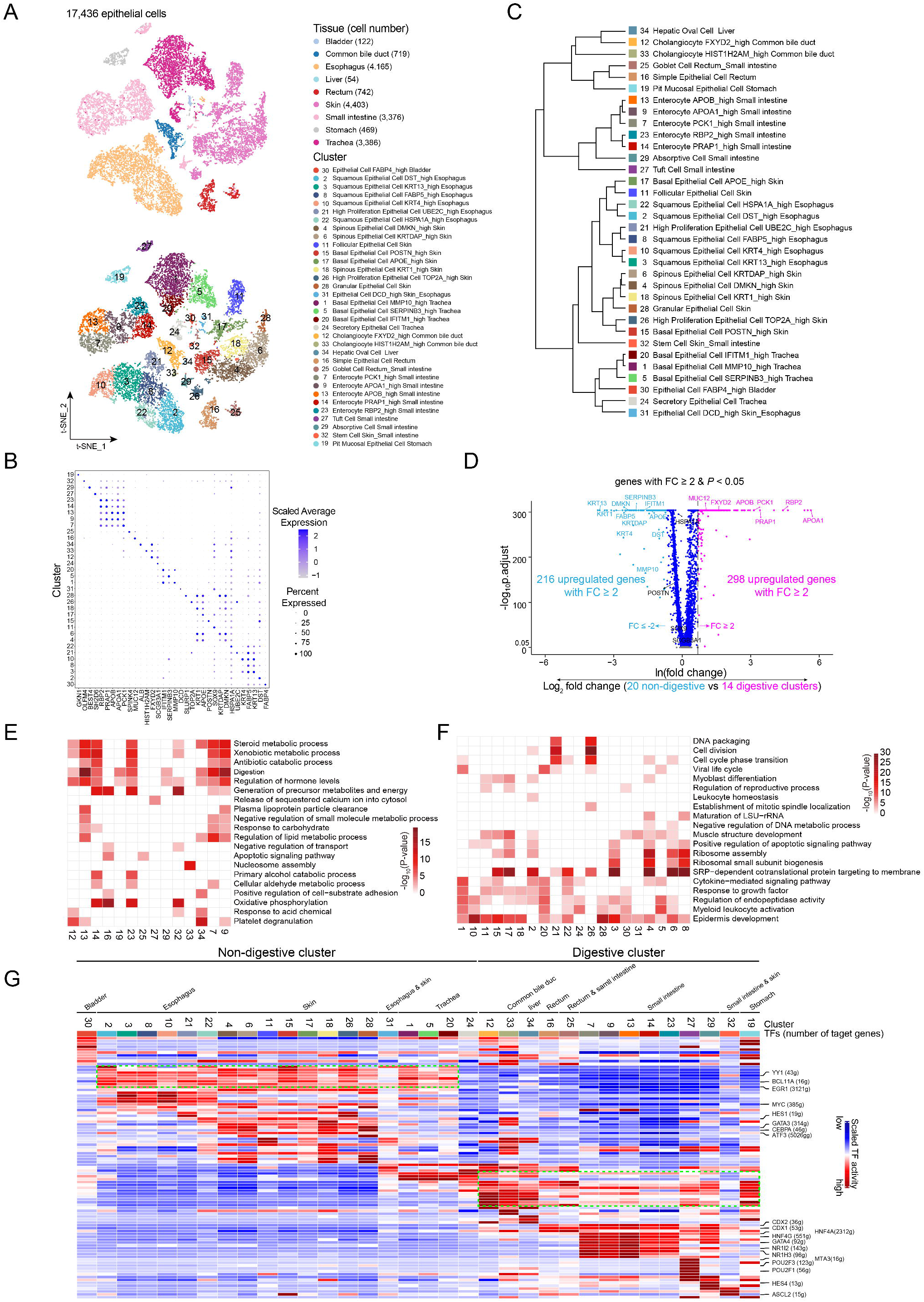
The heterogeneity of epithelial cells inter- and intra-organ tissues. **A.** t-SNE plots of 17,436 epithelial cells. Each dot represents one cell, colored according to their origins of tissues (top panel) or cell clusters (bottom panel). **B.** Dot plot visualizing the normalized expression of marker genes for each epithelial cluster. Cell cluster at y-axis was coded in numbers on the left, corresponding to that in the **Figure 5A**. Maker genes are shown at the x-axis. The size of the dot represents the percentage of cells with a cell type, and the color represents the average expression level. **C.** Dendrogram of 34 clusters based on their normalized mean expression values (correlation distance metric, complete linkage). Only genes with fold change above 1.5, *p.adjust* < 0.05 and pct.1 ≥ 0.2 in each cluster were included in the analysis. **D.** Volcano plot shows the DEGs between the 14 digestive and 20 non-digestive related clusters. Labeled genes are markers for each cluster in **B**. **E.F.** Gene Ontology enrichment analysis results of each epithelial cell cluster in the digestive organs (**E**) and non-digestive organs (**F**). Cell clusters in columns are coded as numbers at the bottom, correspond to that in **Figure 5A**, and are colored according to their −log™*P* values, with white to red for low to high enrichment of a GO term in a row indicated on the right. Only the top 20 significant GO terms (*P*-value < 0.05) are shown. **G.** Heat map of the activation scores of epithelial cell subtypes for gene expression regulated by transcription factors (TFs). Cell clusters are indicated on top, and the scores were estimated using SCENIC analysis. Only the top 10 TFs are shown with the highest difference in expression regulation estimates between each cluster and all other cells, tested with a Wilcoxon rank-sum test. The clusters numbers are in reference to those in **Figure 5A**. Cell clusters are grouped according to the origin of organ and their digestive or non-digestive function as indicated on top.

The epithelial cells were further grouped into 34 clusters (**Fig. 5A**). Grouping of close clusters was seen in all organ tissues, except for tiny clusters of C25 (rectum and small intestine), C31 (esophagus and skin), and C32 (skin and small intestine), suggesting the heterogeneity of epithelial cells at both the intra- and inter-organ contexts (**Additional file 7: Table S36**). The 34 clusters showed different expression profiles with 350 signature genes (FC ≥ 5, pct.1 ≥ 0.2 and pct.2 ≤ 0.2), most of which were expressed exclusively in one cluster (**Fig. 5B, Additional file 1: Figure S18D, F** left panel, and **Additional file 7: Table S37**). Similar results were observed in the previous HCL dataset (**Additional file 1: Figure S18B, E**, and **F** right panel, and **Additional file 7: Table S38-S40**). Hierarchical cluster analysis revealed closer grouping of cell clusters within tissues than across tissues in both our and the HCL datasets (**Fig. 5C, Additional file 1: Figure S18E** and **Additional file 7: Table S39**). Cells from digestive organs including the small intestine, stomach, rectum, and common bile duct were clustered closely, as were the cells from non-digestive tissues including the skin, trachea, and bladder (**Fig. 5C, Additional file 1: Figure S19A**). Although the esophagus is a digestive organ, the epithelial cells were clustered much closer to the skin cells than the digestive organs’ cells. Similarly, the stomach was grouped more closely to non-digestive organs in the HCL dataset, although it is classified as one of the digestive organs together with the ascending colon, colon, sigmoid colon, transverse colon, small intestine, liver, gallbladder, pancreas, and rectum. This might be explained by the different anatomical positions of the specimens (**Additional file 1: Figure S19B**). A volcano plot showed significantly DEG between the two groups, including 514 upregulated genes with a fold change greater than two (298 in the digestion-related clusters and 216 genes in the non-digestion-related clusters; **Fig. 5D**).

To explore the potential functions of cells within each cluster, we performed GO analysis for two groups of cells from the 14 digestion-related clusters and the 20 non-digestion-related clusters, separately. For cells from the digestive sysT_EM_, biological functions related to metabolic process, energy synthesis, and digestion pathways were commonly observed (**Fig. 5E**). For the non-digestion-related cells, the ‘*regulation of endopeptidase activity*’ and ‘*epidermis development*’ were the strongly enriched pathways (**Fig. 5F**). Interestingly, GSVA analysis in both our AHCA and the HCL datasets revealed significantly altered pathways between the digestive and non-digestive clusters, including multiple metabolic pathways with elevated activity in the digestive clusters, such as ‘*Fatty acid metabolism*’, ‘*Citric acid cycle*’, ‘*Protein modification*’, and ‘*Pyrimidine metabolism*’, as well as enrichment of ‘*Epithelial mesenchymal transition*’ and ‘*TNFA signal via NFKB*’ in the non-digestive epithelial cells, suggesting the enhanced metabolic activity of digestive epithelial cells (**Additional file 1: Figure S19C, D**). As we observed the enrichment of immune-related pathways in most of the cell clusters (**Fig. 5E, F**), we examined their antigen-presenting abilities, which revealed a weaker antigen-presenting ability for epithelial cells from the skin and esophagus to present both intra- and extracellular antigens in our AHCA dataset but a higher antigen-presenting ability for epithelial cells from the lungs in the HCL dataset (**Additional file 1: Figure S20A**).

Next, we investigated the contribution of TFs in regulating the heterogeneous transcriptional profiles of epithelial cells in and between organs. SCENIC analysis revealed that the regulation of TFs was similar among cell clusters within a same tissue but was very different among cell clusters between tissues (**Fig. 5G, Additional file 7: Table S41**). Interestingly, the digestion-associated epithelial clusters exhibited similar activation of TFs, as did the cell clusters belonging to the non-digestive tissues including the trachea, skin, and esophagus (**Fig. 5G, Additional file 7: Table S41**). We also observed some cluster-specific TFs in the skin (*CEBPA, HES1, GATA3*, and *ATF3*), small intestine epithelial cells (*HNF4G, NR1H3, NR1I2, HNF4A, CDX1*, and *CDX2*), tuff cells (*MTA3*, *POU2F1*, and *POU2F3*), absorptive cells (*HES4*), and stem cells (*ASCL2*; **Fig. 5G**). Moreover, we identified 204 TFs in our AHCA dataset, which overlapped with about 60% (201) of all cluster-specific TFs in the HCL dataset. A similar activation pattern of TFs was observed between our AHCA and the HCL datasets for digestive-associated tissues (expect for the pancreas) and non-digestive tissues (except for the lungs; **Additional file 1: Figure S20B** and **Additional file 7: Table S42**). Together, these observations suggest that TFs may contribute to the heterogeneity of epithelial cells across tissues and epithelial cells from tissues with similar functions may share a similar activation pattern of TFs.

### The similarity and heterogeneity of stromal cells

Endothelial cells (ECs) line up in a monolayer and form the interior surface of blood and lymphatic vessels as well as heart chambers. We identified a total of 6,932 ECs, including 6,681 blood endothelial cells (BECs, marked with *VWF*), and 251 lymphatic endothelial cells (LECs, marked with *LYVE1*). BECs and LECs could be further grouped into 11 and two clusters, respectively, with unique gene signatures (**Additional file 1: Figure S21 A-C** and **Additional file 8: Table S43**). Although various cell clusters were identified in a tissue, hierarchical cluster analysis revealed that the cell clusters from the same tissue were grouped closer than those between two different tissues (**Additional file 1: Figure S21D**), such as FABP4_BEC and TNFRSF4_BEC, and APOC1_BEC from the muscle. Interestingly, a group of LECs (FCN3_LEC) were strictly identified in the liver and expressed only two of the four genes for LECs (*PECAM1* and *LYVE1;* but not *PDPN* and *PROX1*[43]; **Additional file 1: Figure S21B**, **E**) and other liver-specific markers (*CD4, CD14, FCN2/3, OIT3*, and *CLEC4G*). The other group of LECs (CCL21_LEC) were from tissues except for the liver, with exclusively high expression of *CCL21*, the protein which binds the chemokine receptor 7 (*CCR7*) and promotes adhesion and migration of various immune cells[44]. This suggests that these LECs have a higher potential to attract immune cells than the LECs in the liver. GO analysis results revealed that BECs and LECs from most of the clusters had common endothelial functions including ‘*blood vessel development*’ and ‘*response to wounding*’, as well as functions regulating immune response (**Additional file 1: Figure S21F**). Moreover, the cells from each cluster were shown to have specific biological functions, including ‘*endocrine processing*’ for liver BECs (TIMP1_BEC), ‘*cellular extravasation*’ and ‘*adaptive immune response*’ for heart BECs (ACKR1_BEC), and ‘*macrophage migration*’ for non-liver LECs (CCL21_LEC). We further examined the APS for each cell cluster, which revealed a higher ability in presenting extracellular antigens for BECs from the skin (CTSC_BEC, *P* < 2.2×10^-16^) and the LECs in the liver (FCN3_LEC; *P* = 7.232×10^-15^) than the other tissues (**Additional file 1: Figure S21G**).

We also identified a total of 17,690 fibroblasts and smooth muscle cells from nine tissues. These cells were further grouped into 14 fibroblast clusters (11,697 cells, *MMP2*), four smooth muscle cell clusters (3,165 cells, *ACTA2*), and another five novel clusters assigned as FibSmo (2,828 cells; marked with *MMP2* and *ACTA2;* **Additional file 1: Figure S22A, B**). We observed organ-specific distribution for fibroblasts of different clusters, but a mixture of multiple organs for smooth muscle cell clusters (**Additional file 1: Figure S22A**), which is consistent with previous findings in a mouse model[9]. DEG analysis revealed that cells from each cluster had a unique signature (**Additional file 1: Figure S22C** and **Additional file 8: Table S44**). Hierarchical cluster analysis showed that cells from clusters were grouped in an organ-specific manner, namely close distance for those within a same tissue (**Additional file 1: Figure S22D**). The presence of the novel FibSmo cells was further confirmed by double immunostaining of fibroblast maker MMP2 and smooth muscle cell marker ACTA2 in multiple tissues (**Additional file 1: Figure S10**). Fibroblasts, smooth muscle cells, and FibSmo cells had distinct gene signatures (**Additional file 1: Figure S23A** and **Additional file 8: Table S45**). GO analysis revealed that the three cell types shared classical functions, including ‘*response to wounding*’ and ‘*tissue remodeling*’ (**Additional file 1: Figure S23B**). However, each of them has unique functions. Fibroblasts have specific enrichment of genes that are involved in the ‘*extracellular structure organization*’, which is consistent with their strong expression of extracellular matrix protein genes (*DCN* and *FBLN2*) and genes related to matrix assembly (*MFAP5* and *SFRP2*) and matrix remodeling (*MMP2;* **Additional file 1: Figure S23B, C**). Smooth muscle cells have specific enrichment of genes related to muscle system processing, including *MYH11, MYLK, CAV1*, and *MEF2C* (**Additional file 1: Figure S23B, C**). By contrast, FibSmo cells exhibited a high and specific expression of *PLAT* (**Additional file 1: Figure S7**) and *ID1*, which were related to clotting and angiogenesis[45, 46], as well as a higher expression of *COL3A1* and *COL1A1* (**Additional file 1: Figure S23B, C**), which have been involved in wound healing[47], compared with other stromal cells. Moreover, the enrichment of BMP signal genes *BMP4* and *BMP5* were also observed in the FibSmo cells (**Additional file 1: Figure S23B, C** and **Figure S7**). Furthermore, GO analysis revealed that FibSmo cells had enhanced biological functions in ‘*response to wounding*’ and ‘*growth factor*’, which is consistent with the functions of their highly expressed signature genes detailed above (**Additional file 1: Figure S23B, C**).

### Complex and broad intercellular communication networks within and between tissues

Since we observed heterogeneity for cells in each organ tissue, we explored their potential intercellular communication network. CellphoneDB interaction analysis was conducted to explore cell-cell crosstalk of different cell types in various organs based on the repository of ligands, receptors and their interactions, which mediates cell-cell communication critical to coordinating diverse biological processes[48]. We observed a total of 20,630 significant interactions based on 475 ligand-receptor pairs among cell types and tissues, which varied from 131 in the lymph nodes to 3,229 in the skin (**Additional file 1: Figure S24A**). Among them, *MIF_CD74, HBEGF_CD44, MIF_TNFRSF14, CD55_ADGRE5*, and *APP_CD74* were the top five frequent interacting pairs detected across different cell types (**Additional file 1: Figure S24B** and **Additional file 9: Table S46**), suggesting their important roles in mediating crosstalk between different cell types. Next, we focused on the interactions between pairs of the major cell types (**Additional file 1: Figure S24C** and **Additional file 9: Table S47**). Myeloid cells were the most active cell type interacting with the other types of cells (6,949 inter-cell interactions), especially with epithelial cells (24.3% of the total inter-cell interactions) in the skin and trachea (51.8%). Interestingly, common interactions were observed between myeloid cells and epithelial cells, with the most frequent ligand-receptor pair *HBEGF_CD44* (**Fig. 6A, Additional file 1: Figure S25**), of which dysregulations were involved in tumor and metastasis initiation[49]. CD8^+^ T cells were another cell type with intensive interactions with other types of cells (total inter-cell 4,290 interactions), especially with myeloid cells (29.9% of the total inter-cell interactions). The interactions were found mainly in the liver, trachea, and common bile duct (44.6%). The frequent interaction pairs between CD8^+^ T and myeloid cells were *RPS19_C5AR1, CD55_ADGRE5, MIF_CD74, HBEGF_CD44, CD99_PILRA*, and *ANXA1_FPR1*, most of which play important roles in immune regulation[50–52] (**Fig. 6B, Additional file 1: Figure S26**). CD8^+^ T cells also had broad interactions with non-immune cells (**Additional file 9: Table S47**). For instance, the most frequent interacting chemokine and receptor pair *CXCL12_CXCR4[53]* was observed between stromal (fibroblasts, FibSmo, and smooth muscle cells) and CD8^+^ T cells, suggesting the chemoattraction potential of those stromal cells for T cells migration into tissues (**Fig. 6C, Additional file 1: Figure S27**). We observed broad interactions with different densities among various organ tissues (**Fig. 6D, Additional file 10: Table S48**). We noted that cells in the trachea had a high density of interaction pairs with multiple tissues including the rectum, liver, and spleen, suggesting the potential regulatory communications between these organs. Interestingly, myeloid, CD8^+^ T, and epithelial cells were the core nodes of cell-cell interactions, which had the greatest number of interacting pairs and enhanced pairwise communications (**Fig. 6E, Additional file 10: Table S48**).

**Figure 6.**
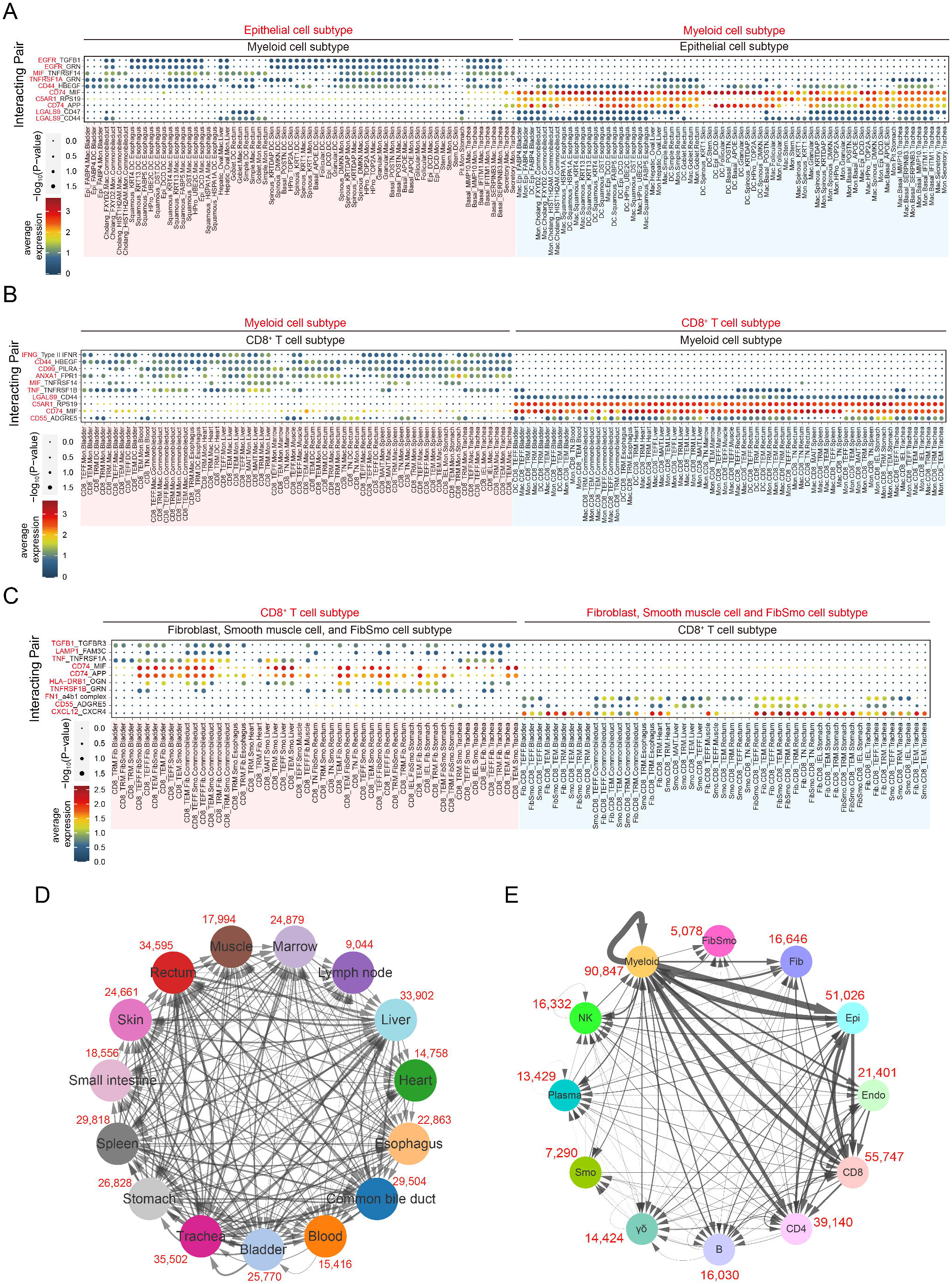
Intercellular communication networks among tissues. **A-C.** The top 10 significant ligand-receptor interactions between cells among different organs for epithelial and myeloid cell subtypes (**A**), myeloid and CD8^+^ T cell subtypes (**B**), and CD8^+^ T and stromal cell subtypes (**C**; Fibroblast, Smooth muscle cell, and FibSmo cell). An interaction is indicated as color-filled circle at the cross of interacting cell types in a tissue (x-axis) and a ligand-receptor pair (y-axis), with circle size representing the significance of - log_10_(*P*-values) in a permutation test and colors representing the means of the average expression level of the interacting pair. The naming system is as follows, taking an example of “EGFR_TGFB1” in “cholang_FXYD2.Mac.Commonbileduct”, the ligand-receptor pair is *EGFR* (red) and *TGFB1* (black), and the circle is colored based on the expression levels of *EGFR* in cholang_FXYD2 cluster and *TFGB1* in Mac cluster in the tissue Commonbileduct. Mon: Monocyte, Mac: Macrophage, DC: Dendritic cell, TRM: Tissue-resident memory T cell, TEFF: Effector T cell, TGD: yδ T cell, MAIT: Mucosal associated invariant T cell, TEM: Effector memory T cell, TIEL: Intraepithelial T lymphocyte, TN: naïve T cell. Fib: Fibroblast, Smo: Smooth muscle cell, FibSmo: Novel cell type named FibSmo cell. Commonbileduct: Common bile duct, Lymphnode: Lymph node, Smallintestine: Small intestine. For epithelial cells, the full names of each cluster refer to **Table S47**. **D.** Connection graph showing the intensity of interactions between one organ to another in colored circles. Interactions were evaluated between major cell types including CD4^+^ T cell, CD8^+^ T cell, yδ T cell, B cell, Plasma cell, myeloid cell, NK cell, epithelial cell, fibroblast, smooth muscle cell, FibSmo cell, and endothelial cell. Numbers in red show the total counts of ligandreceptor pairs between the indicated organ and all others, which include only the unique significant interacting pairs between them (average expression > 0 and *P*-value < 0.05). **E.** Connection graph showing the intensity of interactions within a major cell type or between two major cell types in colored circles. Numbers in red show the total counts of ligand-receptor pairs within or between cell types, which only included the unique significant interacting pairs between them (average expression > 0 and *P*-value < 0.05). CD4, CD4^+^ T cell; CD8, CD8^+^ T cell; yδ: yδ T cell; B, B cell; Plasma, plasma cell; Myeloid, myeloid cell; NK, NK cell; Epi, epithelial cell; Fib, fibroblast; Smo, smooth muscle cell; FibSmo, FibSmo cell; and Endo, endothelial cell.

## DISCUSSION

Here, to the best of our knowledge, we for the first time generated an adult human cell atlas (AHCA), by profiling the single cell transcriptome for 84,363 cells from 15 organs of one adult donor. The AHCA included 252 cell subtypes, each of which was distinguished by multiple marker genes and transcriptional profiles and collectively contributed to the heterogeneity of major human organs. The AHCA empowered us to explore the developmental trajectories of major cell types and identify regulators and interacting networks in one donor that might play important roles in maintaining the homeostasis of the human body. We have made the AHCA publicly available (http://research.gzsums.net:8888), as a resource to uncover key events during the development of human disease in the context of heterogeneity of cells and organs.

It has been demonstrated that T_N_ cells are generated in the thymus and populate lymphoid tissues where they differentiate to T_EFF_ cells upon antigen stimulus, and subsequently develop into long-lived memory T cells[54]. However, how T cells develop into different states throughout human organs and the links across T cells of different states as well as the underlying regulatory networks, are largely unknown, especially in the context of one individual body. In our study, trajectory analysis revealed a clear development route from T_N_ cells into T_EFF_ cells and then CD4^+^ and CD8^+^ T_RM_ cells in non-lymphoid organs with terminal developmental states, which is consistent with previous findings[54]. Indeed, T_RM_ cells share many properties with recently activated effector T cells, supporting the fact that they may constitute a terminally differentiated population[55–57]. However, for CD4^+^ T cells, we noted a clear T_CM_ cell cluster (STMN1_T_CM_) at the end of the trajectory, likely differentiated from T_RM_ cells (TNF_T_RM_). Together with STMN1_T_CM_ cells collected from lymph nodes, this observation provides a clue to solving the puzzle as to whether T_RM_ cells can further differentiate or migrate back to the lymphoid compartment[58]. In support of such, a very recent mice model study demonstrated that T_RM_ cells in the skin could differentiate into T_CM_ and T_EM_ cells upon local reactivation, which then rejoined the circulation. Moreover, trajectory analysis with TNF_T_RM_ and STMN_T_CM_ clusters revealed that TNF_T_RM_ cells from two branches in an early state gradually progressed towards STMN1_T_CM_ cells in a terminal state (**Additional file 1: Figure S28A**). T_RM_ cells at the beginning of the trajectory expressed high levels of T_RM_ markers, such as *RUNX3, NR4A1*, chemokines (*CCL5*)[59], and other T_RM_ associated genes, including *ID2*[60]. By contrast, T_CM_ cells at the end of the trajectory were marked with expression of well-known T_CM_ molecules, such as *SELL* and *CCR7* (**Additional file 1: Figure S28B**). Taken together, these observations suggest that T_RM_ cells have developmental plasticity rather than representing a terminal stage of differentiation[61]. We also noted that T_RM_ cells exhibit development states at organ-specific patterns and consistently these cells were regulated by different types of TFs, suggesting that tissue microenvironments might regulate gene expression by affecting TF’s activity, which in turn, shapes specific T cell phenotypes. TCR analysis tracking cell of a same lineage revealed widespread links among subpopulations of T_RM_ cells, however, these are considered as non-recirculating[62]. Moreover, intensive sharing of TCRs were observed among T_RM_, T_EM_, and T_EFF_ cells (**Additional file 1: Figure S14I, J**). Together with their development states, these results suggest that T_EM_ cells and T_EFF_ cells might enter the tissues and develop into T_RM_ cells and IEL T cells. In addition, we observed the branching out of CTLA4_T_reg_ cells next to KLF2_T_CM_ cells along the development trajectory from T_N_ cells to T_RM_ cells (**Fig. 2E**). Given that previous *in vitro* studies have demonstrated a potential development from central memory T cells to T_reg_ cells upon stimulation[63, 64], this observation suggested that differentiation of T_reg_ cells from central memory T cells is possible. However, whether it holds true *in vivo* remains to be established with precise lineage tracing methods.

Infiltrating macrophages come from classical monocytes in pathological settings, such as cancers[65], while various origins of adult macrophages among tissues in a steady state have been reported[66]. As such, it has been debated how macrophages are renewed in the maintenance of hemostasis in tissues, whether through local proliferation or recruitment of monocytes from peripheral blood[67, 68]. In our study, multiple observations suggest that macrophages in organs are derived from either circulating monocytes or *in situ* expansion and proliferation of macrophage populations coping with the local microenvironments. Consistently, entry of monocytes to steady-state non-lymphoid organs and self-maintenance of tissue macrophages have been reported in mice models[36, 69]. Although our observations suggest a potential developmental relationship between circulating monocytes and tissue-resident macrophages, we acknowledge that further investigations are needed to address whether self-expansion alone, or slightly together with circulating monocytes contributes to the development of tissue-resident macrophages, considering that most tissue-resident macrophages have been demonstrated to be originated from embryonic progenitor cells[36].

We observed a more isolated clustering of epithelial and stromal cells compared with immune cells, which have circulating capability. Because epithelial and stromal cells are fundamental components that form protective barriers and supporting matrix for many organs, disruptions in their homeostasis have been implicated in various diseases[70–73]. Accumulating studies have demonstrated the remarkable functional heterogeneity of epithelial cells among tissues[74–76]. Consistently, our GO enrichment analyses revealed very diverse functions of epithelial cells among different tissues, as well as stromal cells. Taken together, a higher degree of heterogeneity might reflect a higher degree of terminal differentiation states and distinct specific functions of these cells among different organs. Nevertheless, we note that epithelial cells derived from digestive organs had similar biological functions and activation of TFs, which were consistently observed in both our AHCA and the HCL datasets[20]. Given that most of the digestion-related organs develop from the endoderm, this might explain their similarity in genetic profiles and functions[77]. In addition, we suspected that the epithelial cells with the same digestive functions might also share similar responses to pathogens or stimuli[78].

The AHCA not only brings more detailed understanding of cell development and heterogeneity, but also reveals novel cell types and genes as well as regulatory factors that might be important for cell development. We identified subsets of novel cells, including *COCH^+^* fibroblasts and FibSmo cells with a broad distribution among organ tissues. TFs are known as the ‘master regulators’ for gene expression[79, 80]. We identified numerous novel TFs in regulating the development of different cell states of major cell types, such as *CEBPD, EGR1* in CD4^+^ T_RM_, *ELF1* in CD8^+^ T_RM_, and *MAFF* in both CD4^+^ and CD8^+^ T_RM_, *POLR2A* in B cells, *KLF13* in plasma cells, as well as *EGR1, MYC, YY1*, and *BCL11A* in the non-digestive tissues. These findings not only extend our understanding in how the TFs regulate gene expression and shape different phenotypes, but also provide potential gene combinations in reprogramming applications. The AHCA also provides useful data to explore the cellular networks at a single-cell resolution. We discovered a large number of interactions between immune cells and other cells in all tissues, reflecting essential and broad communications between immune cells and other cell types in the human body. Epithelial cells had the most frequent inter-cell interactions compared with other cell types, suggesting that they could interact with each other intensively to regulate their biological functions (**Additional file 1: Figure S29**). Moreover, the AHCA is based on a large-scale of single-cell transcriptomes from multiple organs, which might provide common biological understanding at a higher resolution. First, the heterogeneous nature of human cells in organs is consistent with the findings in mice[8] and a recent study with human tissues[20]. Second, we identified well-known markers for different cell types and well-characterized TFs responsible for cell development, such as *TCF7, SELL, MYC*, and *KLF2* for T_N_ cells, *TBX21, STAT1*, and *IRF1* for T_EFF_ cells, and etc. Third, similar transcription profiles of well-differentiated epithelial cells were observed between the AHCA and the recent published HCL datasets[20] (**Fig. 5A, Additional file 1: Figure S18B**), as were important TFs regulating plasma cell development (**Fig. 3E, Additional file 1: Figure S16F**) and digestion-related cells (**Fig. 5G, Additional file 1: Figure S20B**). Lastly, our discovery of novel and/rare cell types were validated in existing datasets and replicated in independent human samples (**Additional file 1: Figure S4-S10**).

Cellular dissociation is a prerequisite of technical manipulation in single-cell studies. One previous study reported dissociation procedures induced stress and caused transcriptional disturbances of varying degrees, leading to a misinterpretation of results[81]. Enzymes and duration of dissociation procedures might be two important factors[81, 82]. Considering the properties of different enzymes and the heterogeneous cell types in organs, we optimized the protocols to achieve better dissociation and higher cell viability for each organ (**Additional file 11: Table S49**). We observed that most organs had a higher density of low total dissociation scores[81], meaning that dissociation-related genes were not significantly induced in the majority of cells (**Additional file 1: Figure S30**). Moreover, high expression of *FOS*, a dissociation-related gene according to previous studies[81], was observed widely in multiple human tissues before dissociation (**Additional file 1: Figure S31**). Furthermore, each major cell type derived from multiple organs treated with different dissociation procedures shared similar transcriptional profiles as reflected by the cluster analyses. These suggest that the dissociation procedures had minimal effects on the transcriptomes in our study. However, we could not rule out the possibility that the dissociation procedures might have impacts on some cell types, which awaits further investigations.

We acknowledge that the current AHCA has several limitations. First, although we obtained a sufficient number of sequencing reads for each sample, the number of genes detected in each cell was limited. This might underestimate the roles of some lowly expressed genes, such as long non-coding RNAs. Second, we obtained around 5,000 cells on average for each organ, which might limit our ability to identify rare cell types and thus underestimate the heterogeneity of inter-cell interactions in organs. Third, we included only 15 organ tissues from a single donor in our study. Further studies on gene expression profiling at both the transcriptional and protein levels, as well as functional characterization with more organs from a larger number of donors, would provide a much broader and more detailed global view of the human cell atlas and cell biology.

## CONCLUSIONS

We generated an AHCA, by profiling the single cell transcriptome for more than 84,000 cells of 15 organs from one research-consented donor. The AHCA uncovered the heterogeneity of cells in major human organs, containing more than 250 subtypes of cells. Comprehensive analyses of the AHCA enabled us to delineate the developmental trajectories of major cell types and to identify novel cell types, regulators, and key molecular events that might play important roles in maintaining the homeostasis of the human body and or those otherwise developing into human diseases.

## MATERIALS AND METHODS

### Organ tissue collection

An adult male donor who died of a traumatic brain injury was recruited at the First Affiliated Hospital of Sun Yat-sen University (SYSU-1H). For single-cell RNA sequencing, besides the organs for transplantation purposes, we collected tissues from 15 organs in sequence, including blood, bone marrow, liver, common bile duct, lymph node (hilar and mesenteric), spleen, heart (apical), urinary bladder, trachea, esophagus, stomach, small intestine, rectum, skin, and muscle (thigh). All hollow viscera tissues were dissected according to the whole layer structure, and all parenchymal viscera tissues were obtained from the organ lower pole. All the tissue collection procedures were accomplished within 20 minutes to maximize cell viability. To avoid cross-contamination, we used different sets of sterilized surgical instruments. The blood and bone marrow samples were loaded into 10ml anticoagulation tubes containing EDTA (BD Biosciences, Cat. no. BD-366643) and other tissues were placed in physiological saline (4 °C) to wash away the blood and secretions, and then immediately in a D10 resuspension buffer, containing a culture medium (DMEM medium; Gibco™, Cat. no. 11965092) with 10% fetal bovine serum (FBS; Gibco™, Cat. no. 10099141). All tissue samples were kept on ice and delivered to the laboratory within 40 min for further processing. For immunohistochemistry assays, paraffin-embedded normal samples were collected from additional patient donors at the SYSU-1H.

### Tissue dissociation and cell purification

All tissues were dissociated within 1.5 hours and viable cells were collected at the end using fluorescence-activated cell sorting (FACS; BD FACS Aria™ III). For solid tissues exclusive of the liver, each fresh tissue was cut into 1 mm pieces and incubated with a proper digestive solution including enzyme cocktail (**Additional file 11: Table S49**), followed by neutralization with the D10 buffer and then passed through a 40 μm cell strainer (BD, Cat. no. 352340). The cell suspension was centrifuged at 300x g for 5 min at 4 °C, and the pellet was resuspended with a 0.8% NH4Cl (Sigma-Aldrich, Cat. no. 254134-5G) red blood cells lysis buffer (RBCL) on ice for 10 min, followed by an additional wash with the D10 buffer. The liver tissue was cut into 3-4 mm pieces and incubated with 1 mM EGTA (Sigma-Aldrich, Cat. no. E0396-10G) in 1 x DBPS (Gibco™, Cat. no. 14190250) for 10 min at 37 °C with rotation at 50 rpm. After washing with 1 x DPBS to remove EGTA, each tissue was then incubated in a pre-warmed digestion buffer (**Additional file 11: Table S49**) with rotation at 100 rpm at 37 °C for 30 min. The liver cell suspension was carefully passed through a 70 mm nylon cell strainer (BD, Cat. no. 352350), which was further centrifuged at 50x g for 3 min at 4°C to pellet hepatocytes. The supernatant was centrifuged at 300x g for 5 min at 4°C to pellet non-parenchymal cells. The pellet was resuspended and treated with RBCL. The blood and bone marrow samples were pelleted by centrifugation at 300x g for 5 min at 4 °C and resuspended with RBCL on ice for 10 min, followed by an additional wash with the D10 buffer. All cells from each tissue were resuspended with the D10 buffer to a concentration of 50-500 million cells per milliliter and stained with Calcein AM (Component A: AM) and Ethidium homodimer-1 (Component B: EH) in LIVE/DEAD Viability/Cytotoxicity Kit (Invitrogen, Cat. no. L3224) for 20 min on ice. Only the AM^+^EH^-^ cells were collected by FACS for each tissue.

### cDNA library preparation

The concentration of single cell suspension was determined using a Cellometer Auto 2000 instrument (Cellometer) and adjusted to 1,000 cells/μl. Approximately 14,000 cells were loaded into a CHROMIUM instrument (10x Genomics, CA, USA) according to the standard protocol of the Chromium single cell V(D)J kit in order to capture 5,000 ~ 10,000 cells per channel. In brief, mRNA transcripts from each sample were ligated with barcoded indexes at 5’-end and reverse transcribed into cDNA, using GemCode technology (10x Genomics, USA). cDNA libraries including the enriched fragments spanning the full-length V(D)J segments of T cell receptors (TCR) or B cell receptor (BCR), and 5’-end fragments for gene expression were separately constructed, which were subsequently subjected for high-throughput sequencing.

### Single-cell RNA sequencing data processing

5’-end cDNA, TCR and BCR libraries were mixed and subjected for sequencing on Illumina HiSeq XTen instruments with pared-end 150 bp. Raw data (BCL files) from HiSeq platform was converted to fastq files using Illumina-implemented software bcl2fastq (version v2.19.0.316). cDNA reads were aligned to the human reference genome (hg38) and digital gene expression matrix was built using STAR algorithm in CellRanger (“count” option; version 3.0.1; 10x Genomics)[83]. TCR and BCR reads were aligned to human reference VDJ dataset (http://cf.10Xgenomics.com/supp/cell-vdj/refdata-cellranger-vdj-GRCh38-alts-ensembl-2.0.0.tar.gz) using CellRanger (“vdj” option; version 3.1.0; 10x Genomics). Parameters were set as default except for “force-cells” as 13,000. Raw digital gene expression matrix in the “filtered_feature_bc_matrix” file folder generated by CellRanger was used for further analysis.

### Cell clustering, doublet identification, and differential gene expression analysis

Quality control filtering, variable gene selection, dimensionality reduction and clustering for cells were performed using the Seurat package[6] (version 3.1.5; https://satijalab.org/seurat). “DoubletFinder” (version 2.0.3; https://github.com/chris-mcginnis-ucsf/DoubletFinder) was used to identify doublets in each organ. All the analytic packages were performed in R software (version 3.6.3; https://www.r-project.org), with default settings unless otherwise stated. For each tissue, output cells were forced to 13,000 under ‘cellranger count’ module, and we removed cells with low quality (UMI < 1,000, gene number < 500, and mitochondrial genome fragments > 0.25) as well as genes with rare frequencies (0.1% of all cells). For the remaining cells, gene expression counts data for each sample was normalized with “NormalizedData” function, followed by scaling to regress UMIs and mitochondrial content using “ScaleData” function (negative binomial model). Principal Component Analysis (PCA) and t-SNE implemented in the “RunPCA” and “RunTSNE” functions, respectively, were used to identify the deviations among cells. Genes with high variations were identified using “FindVariableGenes” and included for PCA (“mean.cutoff” ≥ 0.1, “dispersion.cutoff” ≥ 0.5). We used a different value of perplexity and the number of principal components (PCs) determined by elbow plots for each tissue and cell type (**Additional file 11: Table S50**). Cell clusters were identified using the “FindClusters” function and shown using t-SNE. Subsequently, “DoubletFinder” was used to identify doublets using the same PCs in PCA analysis above, assuming the 5% doublet formation rate to the loaded cells for each sample in a droplet channel. The optimal pK values were determined for each organ based on the Mean-variance normalized bimodality coefficient (BCmvn; **Additional file 11: Table S51**). After doublet removal, we rerun the above analyses. Next, differential expression markers or genes were determined using the Wilcoxon test implemented in the “FindAllMarkers” function, which was considered significant with an average natural logarithm (fold-change) of at least 0.25 and a Bonferroni-adjusted *P*-value lower than 0.05. Subsequently, the candidate markers were reviewed and were used to annotate cell clusters. We further manually removed cell clusters that had multiple well-defined marker genes and overlapped gene profiles of multiple different cell types (**Additional file 2: Table S4-S18**). For analyses of the merged data from all tissues, we used 30 PCs and a resolution parameter set to 1 for cell clustering.

### Identification and removal of highly transcribed genes with contamination potentials

Because we observed that some cell-specific genes were broadly expressed among all cell types in a tissue, for example, *APOC3* in enterocytes cells in the small intestine with an average of more than 200 UMIs in a cell, we suspected that if a fraction of a certain type of cells were broken during the sample processing, cell-specific genes with high transcriptions would be released and thus contaminate all cell droplets. Especially, as such the genes would screw the differential expression analysis of cell type among all tissues. Therefore, we identified these genes and removed them for comparison analyses within major cell types. We assumed that non-epithelial cells from a same linage have similar gene profiles at a certain degree and that a few genes would have modest expression in only one organ. Here is a given example for T cells, in which we grouped together the cells previously labeled with NK/T, T, and immune cells (**Additional file 2: Table S4-S18**) in each tissue. Next, we determined the top 2% of genes with high transcription in each tissue sample based on the total number of UMIs (**Additional file 11: Table S52**) and marked these as potentially contaminating genes. We further randomly sampled 300 cells for each cell type in a tissue and as such generated artificial data for all tissues, with which differential expression gene analysis was performed using the “FindAllMarker” function. Any gene with a normal logarithm of FC above 0.25 and with an expression in less than 5% cells in all other tissue samples was considered a contaminating gene. We performed the above sampling and calculation for four independent times, with seed numbers 1 to 4 in the “FindMarker” function, and only the genes commonly observed in the four calculations were determined as contaminating genes. After removing the above contaminating genes, we performed cluster analysis with the first 20 PCs and a resolution of 1.5. After further removal of cell clusters that had multiple well-defined marker genes of different cell types, we repeated cluster analysis using a lower resolution setting and removing the genes encoding immunoglobulins from the gene expression matrices (**Additional file 11: Table S50**) to identify CD4^+^, CD8^+^, and NK cell (**Additional file 11: Table S53**) clusters. For B cells, plasma cells, endothelial cells, macrophages, monocytes and fibroblasts, we applied a similar strategy to remove contaminating genes (**Additional file 11: Table S54-S58**) and cell clusters with multiple cell-specific markers.

### Trajectory analysis

We performed trajectory analysis using Monocle3 alpha[4] for all tissue-derived T cells, macrophages/monocytes, according to the general pipeline (http://cole-trapnell-lab.github.io/monocle-release/monocle3/). For T cells, after identification of T cells clusters for CD4^+^ and CD8^+^ T, raw gene expression counts of cells were imported to the software. Only genes matching the thresholds (both of mean expression and dispersion ratio greater than 0.15 for CD4^+^, CD8^+^ T cell and myeloid cells) were used for cell ordering and training the pseudo-time trajectory. For investigating the dynamic gene expression between T_RM_ and T_CM_, we extracted those two CD4^+^ T clusters and performed the trajectory analysis using Monocle2 with 1,000 high variable genes. For trajectory analysis of intestinal macrophages/monocytes, we extracted the cell clusters with more than 50 cells in our dataset and combined 50 embryonic macrophages from a previous study[39]. The analysis was done by correcting the batch effect using function “align_cds” of Monocle3 with 25 PCs included.

### TCR and BCR analysis

We assessed the enrichment of TCR and BCR in various organs using R package STARTRAC (version 0.1.0)[23], which included only the cells with the certain clonotypes assigned by Cellranger (version 3.1.0 with updated algorithms to improve the identification of TCR/BCR clonotypes) and with paired chains (α and β for T cells, heavy and light chains for B cells). In brief, cells sharing identical TCR or BCR clones between tissues were measured using migration-index score, and the degree of cell linking between different clusters of T cells or B cells was determined by the transition-index score. TCR or BCR diversity (Shannon-index score) was calculated using “1 - expansion-index score”. For the detailed pipeline, please refer to the website (https://github.com/Japrin/STARTRAC/blob/master/vignettes/startrac.html).

### Presenting-antigen score

To evaluate the antigen-presenting ability of extra- and intracellular of each cell, antigenpresenting score (APS) were calculated using the “AddModuleScore” function implemented in the Seurat package, with gene sets “MHC_CLASS_II_ANTIGEN_PRESENTATION”, and “REACTOME_CLASS_I_MHC_MEDIATED_ANTIGEN_PROCESSING_PRESENTATION” pathways, respectively, in the REACTOME database (http://www.reactome.org).

### Gene Ontology (GO) pathway enrichment analysis

All GO enrichment analysis was performed using the online tools metascape[84] with the “multiple gene list mode” (http://metascape.org/gp/index.html). For epithelial cells, we selected genes with FC ≥ 2 and *p.adjust* < 0.05 for each tissue, while FC ≥ 1.5, *p.adjust* < 0.05, and ptc.1 > 0.2 for each cluster. For non-epithelial cells, we selected genes with lnFC ≥ 0.25, *p.adjust* < 0.05 and ptc.1 > 0.2 in each cluster. Only the genes that ranked in the top 150 according to the FC were used for comparisons in each tissue and subpopulation. The background was given as all the genes expressed in corresponding cell types. In addition, only the gene sets in the “GO Biological Processes” were considered.

### Cellular interaction analysis

To investigate the cellular interaction, we identified the inferred paired molecules using CellphoneDB software (version 2.0)[18] with default parameters. First, to facilitate the pairwise analyses by reducing cell types, we grouped clusters within major cell types, including T, B, NK, myeloid, endothelial cells, fibroblasts, smooth muscle cells and FibSmo cells within each tissue. Moreover, we manually assigned ITGB1_T_N/CM_, LEF1_T_N/CM_, and GADD45B_T_N/CM_ as T_CM_ cell cluster, and KLF2_T_N/CM_ as T_N_ cell cluster based on their expression profiles and the developmental states along the trajectory trees, although naïve and memory T cell clusters could not be accurately determined in our study. The grouped cell types were as follows, CD4^+^ T cell (T_N_, T_CM_, T_reg_, T_EM_, T_h1_, and T_RM_), CD8^+^ T cell (T_N_, T_CM_, T_EM_, T_EFF_, T_RM_, IEL, and MAIT), yδ T cell, B cell (naive and memory B cell), NK cell, plasma cell, myeloid cell (monocyte: Mon, macrophage: Mac, and dendritic cell: DC), endothelial cell (BEC and LEC), fibroblast (Fib), smooth muscle cell (Smo), and FibSmo cell (FibSmo). We did not merge subpopulations of epithelial cells because of their high degree of heterogeneity in each tissue. Considering the test efficiency and computational burden, we focused on cell types with more than 30 cells and only randomly selected 250 cells of each cell type for analysis in each tissue. The significant ligand-receptor pairs were filtered with a *P* value of less than 0.05 and an average expression of interacting pairs larger than 0. All the analyses above were performed as tissue independent. For the analysis of interaction across organs, we only calculated between any one of the immune cells in a tissue and its interacting cells from a different organ. Visualization of interaction network was done using Cytoscape (version 3.7.0).

### Single-cell regulatory network inference and clustering (SCENIC) analysis

We conducted SCENIC analysis on cells passing the quality controls for each major cell types, using R package SCENIC (version 1.1.3) as previously described[85]. Regions for TFs searching were restricted to 10k distance centered the transcriptional start site (TSS) or 500bp upstream of the TSSs. Transcription factor binding motifs (TFBS) over-represented on a gene list and networks inferring were done using R package RcisTarget (version 1.6.0) and GENIE3 (version 1.8.0), respectively, with the 20-thousand motifs database. We randomly selected no more than 250 cells for each cell cluster. The input matrix was the normalized expression matrix from Seurat. The cluster-specific TFs of one cluster were defined as the top 10 or 15 highly enriched TFs according to a decrease in fold change compared with all the other cell clusters using a Wilcoxon rank-sum test.

### Validation analysis in existing datasets

For B and epithelial cells in the HCL dataset[20], we applied similar procedures for cell clustering and differential gene expression analyses as described above. We only extracted adult B and epithelial cell clusters identified by the HCL dataset, considering high sequencing coverage and with less potential cross-cell contamination compared with the other cell types. Genes encoding immunoglobulin were removed in the epithelial cells from raw counts data before further analysis. We set the “mean.cutoff” and “dispersion.cutoff” as 0.05 and 0.2 in “FindVariableFeatures” step for both B and epithelial cells analysis.

For validation of *COCH^+^* fibroblasts[86–88], sweat gland cells[86, 87], and FibSmo cells[20, 89], cells from each individual dataset were merged and batch effect was removed using the “FindIntegrationAnchors” and “IntegrateData” functions in the Seurat package. The downstream analyses followed the same pipeline of our AHCA dataset.

### Calculation of dissociation scores

For each organ, principal component analysis was performed on a subset of 140 human homologous dissociation-related genes as described previously[81]. The first principal component was used as the ‘dissociation score’ as it corresponds to the variance within these genes.

### Analysis of differential pathway

Gene set variation analysis (GSVA) or gene set enrichment analysis (GSEA) was performed to identify significantly enriched genes in each transcriptional dataset, using R package GSVA (version 1.34.0) or GSEA software (version 4.0.3) (https://www.gsea-msigdb.org/gsea/index.jsp) on the 50 hallmark pathways with default parameters, respectively. For epithelial cells in AHCA and HCL datasets, GSVA analysis was performed on the 50 hallmark pathways and additional curated metabolic pathway dataset[90].

### Immunofluorescence staining assay

For immunofluorescence staining assay, tissue samples were collected within 20 min, washed with 1 x DPBS, fixed in 4% paraformaldehyde (pH 7.0), and embedded in paraffin according to routine methods. These paraffin blocks were cut into 4-μm slides and adhered on the slides glass. The sections were deparaffinized, rehydrated, and subjected to blockade of endogenous peroxidase activity with 3% H_2_O_2_, and high-temperature antigen retrieval. The tissues were incubated with 3% BSA at room temperature for 30 min, and then incubated overnight at 4 °C with the primary detection antibodies for different organs (**Additional file 11: Table S59**). The slides were then incubated with the secondary antibody (HRP polymer, anti-rabbit IgG) at room temperature for 50 min. Subsequently, fluorophore (tyramide signal amplification, TSA plus working solution; Servicebio, Cat. no. G1222/3/4) was applied to the tissues. The slides were microwave heat-treated after each TSA treatment and the primary antibodies were applied sequentially for different organs, followed by incubation with the secondary antibody and TSA treatment. Nuclei were stained with 4’-6’-diamidino-2-phenylindole (DAPI; Invitrogen, Cat. no. D1306) after all the antigens had been labeled. Negative controls were performed using similar procedure above, expect for replacing the primary antibody with 1x DPBS. To obtain multispectral images, slides were scanned using the Pannoramic MIDI II system (3DHISTECH, Hungary).

## AUTHOR CONTRIBUTIONS

JXB, XSH and ZYG conceived and supervised this study. SH, LHW, YL, YQL, JHX, HTC, WP, GWL, PPW, BL, XXJ, and DW performed the sample preparation, library construction, and data generation. SH, LHW, YL, and JXB performed the data analyses and annotated the results. SH, JXB, and ZYG wrote the manuscript. All authors read and approved the final manuscript.

## ACKNOWLEDGEMENTS

We acknowledge Chunlin Luo, Xixi Chen, Xiangyu Xiong, Shuqiang Liu, Xiaochen Xu, Yaqing Zhou, Yanchun Feng, Jingjing He, and Liping Xu from SYSUCC for performing the tissue dissociation, and Cliff Y Yang, Junchao Dong, and Jun Chen from Sun Yat-sen University Zhongshan School of Medicine for scientific discussions. We thank all staffs at the High-Throughput Analysis Platform (HTAP) of SYSUCC for data generation and processing.

## Ethics approval and consent to participate

Informed consent for the sample collection for single-cell RNA sequencing was obtained from the donor’s family delegate. The management of organs was compiled according to the organ procurement guidelines. For samples collected for immunohistochemistry assays, informed consent was obtained from each donor’s family delegate. This study was approved by the Ethics Committee for Clinical Research and Animal Trials of the First Affiliated Hospital of Sun Yat-sen University (permit No. [2018]255) and was conducted in accordance with the Declaration of Helsinki principles.

## Availability of data and materials

The AHCA dataset has been deposited in Gene Expression Omnibus (GEO) repository with the primary accession code GSE159929[91]. For B and epithelial cells in the HCL dataset[20], we obtained the gene expression matrices excluding batch genes from the website (https://figshare.com/articles/HCL_DGE_Data/7235471). Two skin datasets were available at the Gene Expression Omnibus (GEO; https://www.ncbi.nlm.nih.gov/geo/), with accession numbers GSE130973[86] and GSE147424[87]. A heart dataset was retrieved via “https://singlecell.broadinstitute.org/single_cell/study/SCP498/transcriptional-and-cellular-diversity-of-the-human-heart”[88]. Two FibSmo cell datasets were obtained from the websites[20, 89] (https://figshare.com/articles/Single-cell_transcriptomic_map_of_the_human_and_mouse_bladders/8942663/1 and https://db.cngb.org/HCL). All related codes and data analysis scripts are available at https://github.com/bei-lab/scRNA-AHCA[92] and Zenodo (doi: 10.5281/zenodo.4136735)[93].

## Supplementary information

**Addition file 1. Figure S1-S31.**

**Figure S1.** Quality information of single-cell RNA sequencing, related to Figure 1. **Figure S2.** Distribution of cell cluster in each organ tissue, related to Figure 1. **Figure S2.** Distribution of cell cluster in each organ tissue (continued). **Figure S3.** The expression of marker genes and distribution of sweat gland epithelial cells and a novel fibroblast subtype, related to Figure 1. **Figure S4.** Expression of *DCD, SCGB2A2, PIP, KRT19*, and *MUCL1* genes in skin cells derived from public datasets, related to Figure 1. **Figure S5.** Expression of *COCH* and *MMP2* in cells from multiple human samples from public datasets, related to Figure 1. **Figure S6.** Gene signature of *COCH^+^* fibroblasts in skin, related to Figure 1. **Figure S7.** Expression of marker genes in cells from bladder samples derived from public datasets, related to Figure 1. **Figure S8.** Representative images of immunofluorescence staining of sweat gland epithelial cells in skin samples. **Figure S9.** Representative images of immunofluorescence staining of COCH^+^ fibroblasts in the skin and heart tissues. **Figure S10.** Representative images of immunofluorescence staining of FibSmo cells in the bladder, rectum, and heart tissues. **Figure S11.** Distribution of cell clusters in different organs, related to Figure 1. **Figure S12.** Comparisons of cell-type determinations by whole single-cell transcriptome dataset and manual annotation of each organ, related to Figure 1. **Figure S13.** The heterogeneity and developmental trajectory analysis of T cells, related to Figure 2. **Figure S14.** The heterogeneity and clonalities of T cell clones in human body, related to Figure 2. **Figure S15.** The heterogeneity and clonalities of T cell clones in the human body, related to Figure 2. **Figure S16.** The heterogeneity of B and plasma cells, related to Figure 3. **Figure S17.** Heterogeneity and trajectory analysis of macrophages, related to Figure 4. **Figure S18.** The gene expression heterogeneity of epithelial cells among inter- and intra-organ tissues, related to Figure 5. **Figure S19.** Expression profiles and altered pathways in digestive and nondigestive epithelial cell clusters. **Figure S20.** The antigen-presenting heterogeneity and regulons of epithelial cells among inter- and intra-organ tissues, related to Figure 5. **Figure S21.** The heterogeneous expression pattern of endothelial cells. **Figure S22.** The similarity and heterogeneity of fibroblasts and smooth muscle cells. **Figure S23.** Gene expression and biological functions of fibroblasts, smooth muscle and FibSmo cells. **Figure S24.** Overview of the ligand-receptor interactions between different cell types, related to Figure 6. **Figure S25.** Overview of the ligand-receptor interactions between epithelial and myeloid cells, related to Figure 6. **Figure S26.** Overview of the ligand-receptor interactions between CD8^+^ T and myeloid cells, related to Figure 6. **Figure S27.** Overview of the ligand-receptor interactions between CD8^+^ T and stromal cells (fibroblast, smooth muscle cell, and FibSmo cell), related to figure 6. **Figure S28.** Pseudotime trajectory analysis and gene expression profiles of TNF_T_RM_ and STMN1_T_CM_. **Figure S29.** Overview of the ligand-receptor interactions within epithelial cells. **Figure S30.** Histogram plots of the density of dissociation scores for each organ. **Figure S31.** Representative images of immunofluorescence staining of FOS and CD8A human tissue samples.

**Additional file 2. Table S1-S20.**

**Table S1.** Basic information of sequencing, related to Figure 1. **Table S2.** Cell number, the median of UMI and genes detected in each organ, respectively, related to Figure 1. **Table S3.** Marker genes and related references. **Table S4-S18.** List of markers information (top 50) for each cell type in the 15 organs, related to Figure 1. **Table S19.** List of markers information (top 50) for each cell type in the merged dataset, related to Figure 1. **Table S20.** Cell counts in each organ for each cluster indicated in Figure S11.

**Additional file 3. Supplementary Notes**

This supplementary material includes a detailed description for the validation of sweat gland epithelial cells, *COCH^+^* fibroblasts, and FibSmo cells in existing datasets, and the evidence showing that the expression of *HSPA1A, FOS*, and *JUN* in the CD8^*+*^ T cells are unlikely stress-induced artefacts.

**Additional file 4. Table S21-S27.**

**Table S21.** Distribution of major cell types in each organ, related to Figure 2. **Table S22.** List of marker information (top 50) for each subpopulation of CD4^+^ T cells, related to Figure 2. **Table S23.** List of marker information (top 50) for each subpopulation of CD8^+^ T cells, related to Figure 2. **Table S24.** List of TFs information for each subpopulation of CD4^+^ T cell, related to Figure 2. **Table S25.** List of TFs information for each subpopulation of CD8^+^ T cells, related to Figure 2. **Table S26.** Detailed information of CD4^+^ TCR repertoire, related to Figure 2. **Table S27.** Detailed information of CD8^+^ TCR repertoire, related to Figure 2.

**Additional file 5. Table S28-S32.**

**Table S28.** List of marker information for each subpopulation of B and plasma cells in AHCA dataset, related to Figure 3. **Table S29.** List of TFs information for each B and plasma cells subpopulation in AHCA dataset, related to Figure 3. **Table S30.** List of marker information for each subpopulation of B and plasma cells in HCL dataset, related to Figure 3. **Table S31.** List of TFs information for each subpopulation of B and plasma cells in HCL dataset, related to Figure 3. **Table S32.** Detailed information of BCR repertoire, related to Figure 3.

**Additional file 6. Table S33-S34.**

**Table S33.** List of marker information (top 50) for each subpopulation of myeloid cells, related to Figure 4. **Table S34.** List of TFs information for each myeloid cell subpopulation, related to Figure 4.

**Additional file 7. Table S35-S42.**

**Table S35.** List of marker information for epithelial cells of each organ in AHCA dataset, related to figure 5. **Table S36.** Cell counts in each organ for each cluster indicated in Figure 5C in AHCA dataset. **Table S37.** List of marker information (top 50) of each subpopulation of epithelial cells in AHCA dataset, related to figure 5. **Table S38.** Marker genes and related references for HCL epithelial cells. **Table S39.** Cell counts in each organ for each cluster indicated in Figure S18E in HCL dataset. **Table S40.** List of marker information (top 50) of each subpopulation of epithelial cells in HCL dataset, related to figure 5. **Table S41.** List of TFs information for each subpopulation of epithelial cells in AHCA dataset, related to Figure 5. **Table S42.** List of TFs information for each subpopulation of epithelial cells in HCL dataset, related to Figure 5.

**Additional file 8. Table S43-S45.**

**Table S43.** List of marker information (top 50) for each endothelial cell cluster. **Table S44.** List of marker information (top 50) for each fibroblast, smooth muscle and FibSmo cell cluster. **Table S45.** List of marker information for fibroblast, smooth muscle and FibSmo cell.

**Additional file 9. Table S46-S47.**

**Table S46.** Frequency of potential interacting pairs, related to Figure 6. **Table S47.** Detailed information of interacting pairs in each tissue related to Figure 6.

**Additional file 10. Table S48.**

**Table S48.** Detailed information of interacting pairs across tissues, related to Figure 6.

**Additional file 11. Table S49-S59.**

Table S49. **The digestion protocols for each organ.** Table S50. **The PCs and resolution used for clustering of each organ or major cell type.** Table S51. **Optimal pK values for each organ.** Table S52. **Basic information of the top 2% genes with high UMI in each tissue.** Table S53. **List of marker information (top 50) for each subpopulation of NK cells.** Table S54. **Suspiciously contaminated genes removed in each tissue for fibroblast, smooth muscle and FibSmo cell clustering.** Table S55. **Suspiciously contaminated genes removed in each tissue for T and NK cell clustering.** Table S56. **Suspiciously contaminated genes removed in each tissue for B and plasma cell clustering.** Table S57. **Suspiciously contaminated genes removed in each tissue for endothelial cell clustering.** Table S58. **Suspiciously contaminated genes removed in each tissue for myeloid cell clustering.** Table S59. **Antibodies used for immunostaining.**

