## Supplementary figure 1-31 for "Single-cell transcriptome profiling of an adult human cell atlas of 15 major organs"

Appendix Figure S1

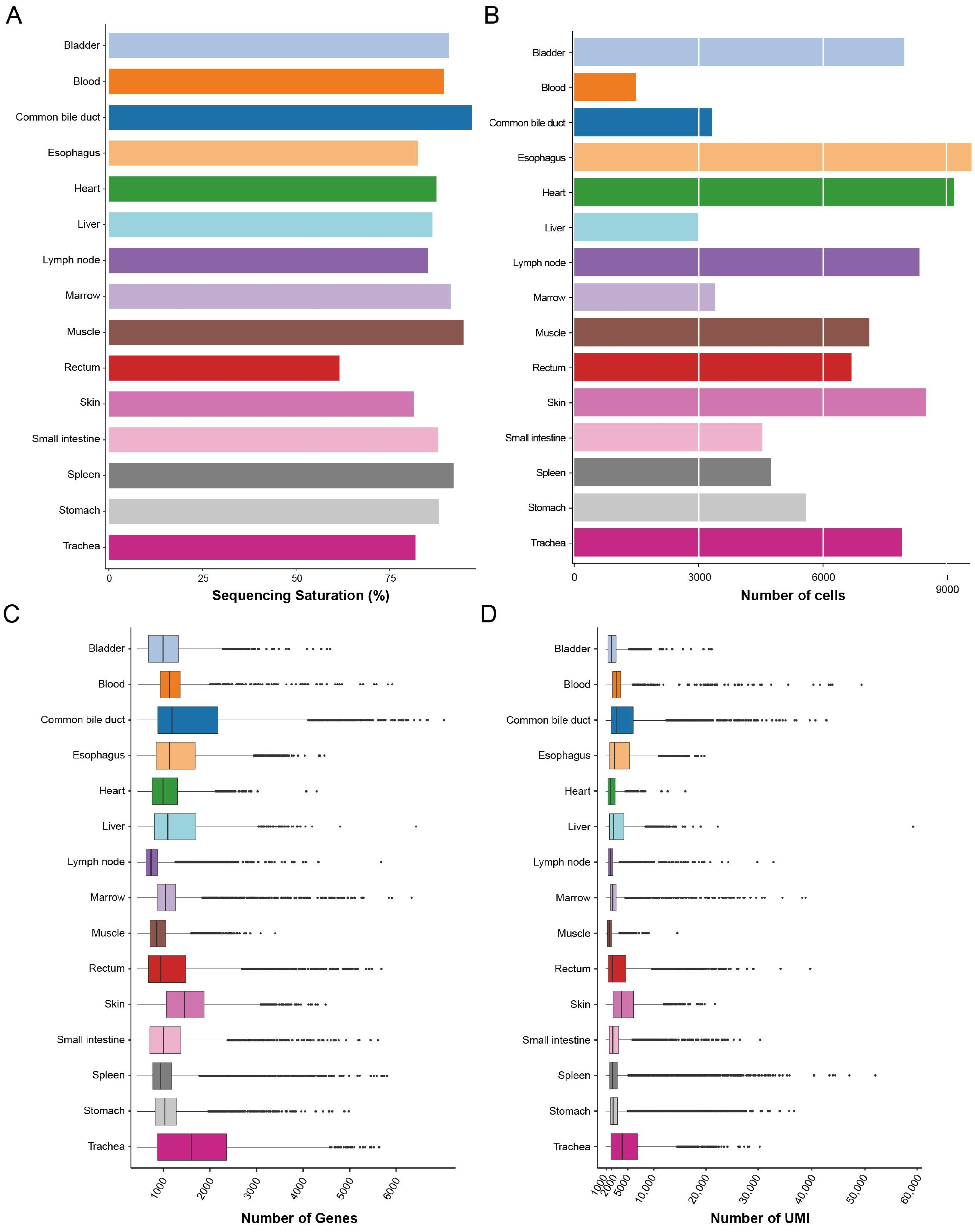


**Quality information of single-cell RNA sequencing, related to Figure 1.**
**A.** Bar plot showing the sequencing saturation of each organ indicated on the left.
**B.** Bar plot showing the numbers of high-quality cells (UMI ≥ 1000, Gene number ≥ 500 and the proportion of mitochondrial gene UMI ≤ 0.25) in each organ indicated on the left (colored).
**C, D.** Box plots showing the numbers of genes (**C**) and UMI (**D**) detected in each cell of each organ indicated on the left.

Appendix Figure 2

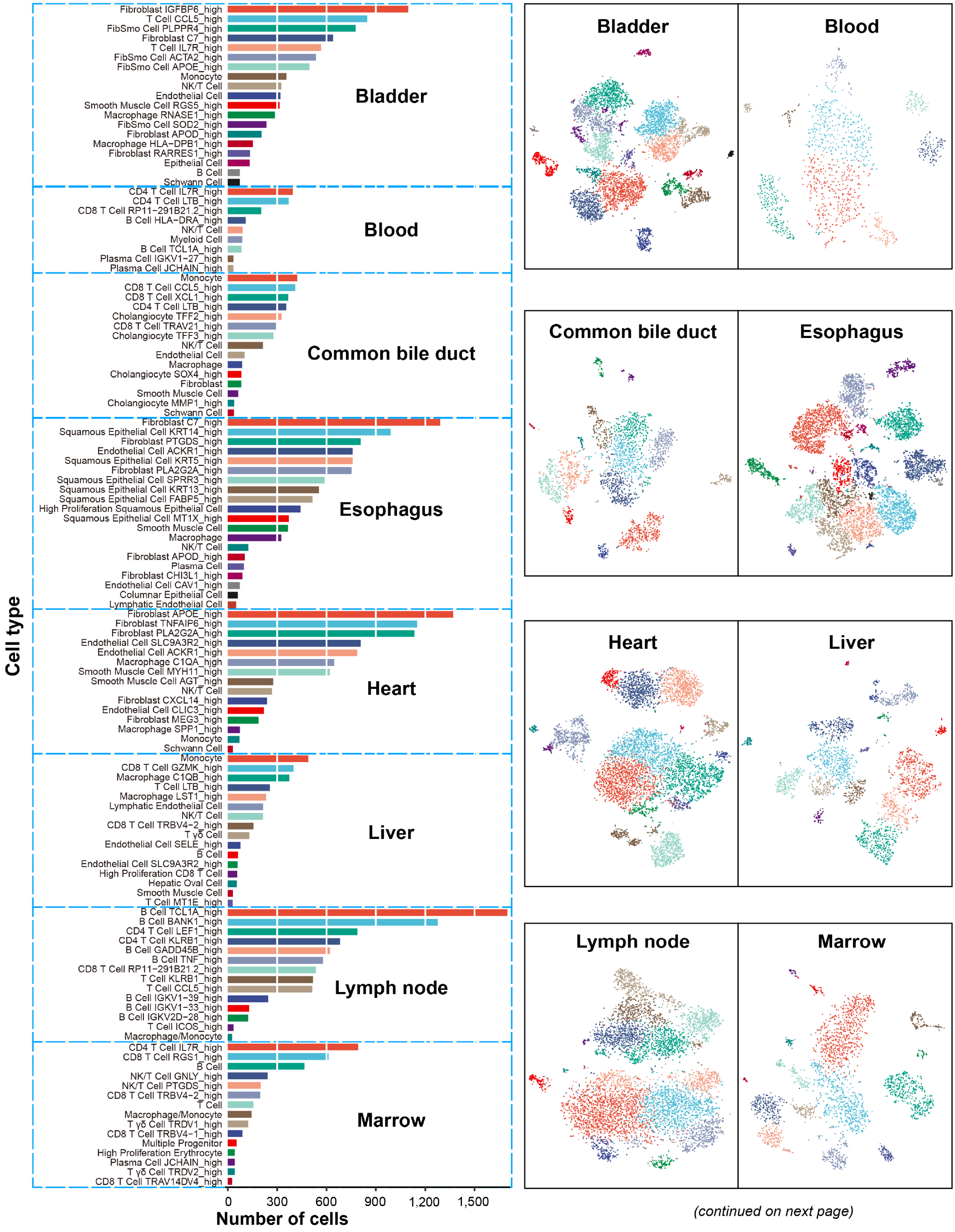


(continued on next page)

**Distribution of cell cluster in each organ tissue, related to Figure 1.**
Bar plots on the left panels show the numbers of cells of different cell types, and the t-SNE plots on the right panels, show their relevant clustering. Cell types indicated on the left were determined by differential gene expression of known markers between clusters. The colors for each cell type are consistent between the bar plot and t-SNE plot for the same organ. Each dot in the t-SNE plot represents one cell.

Appendix Figure S2

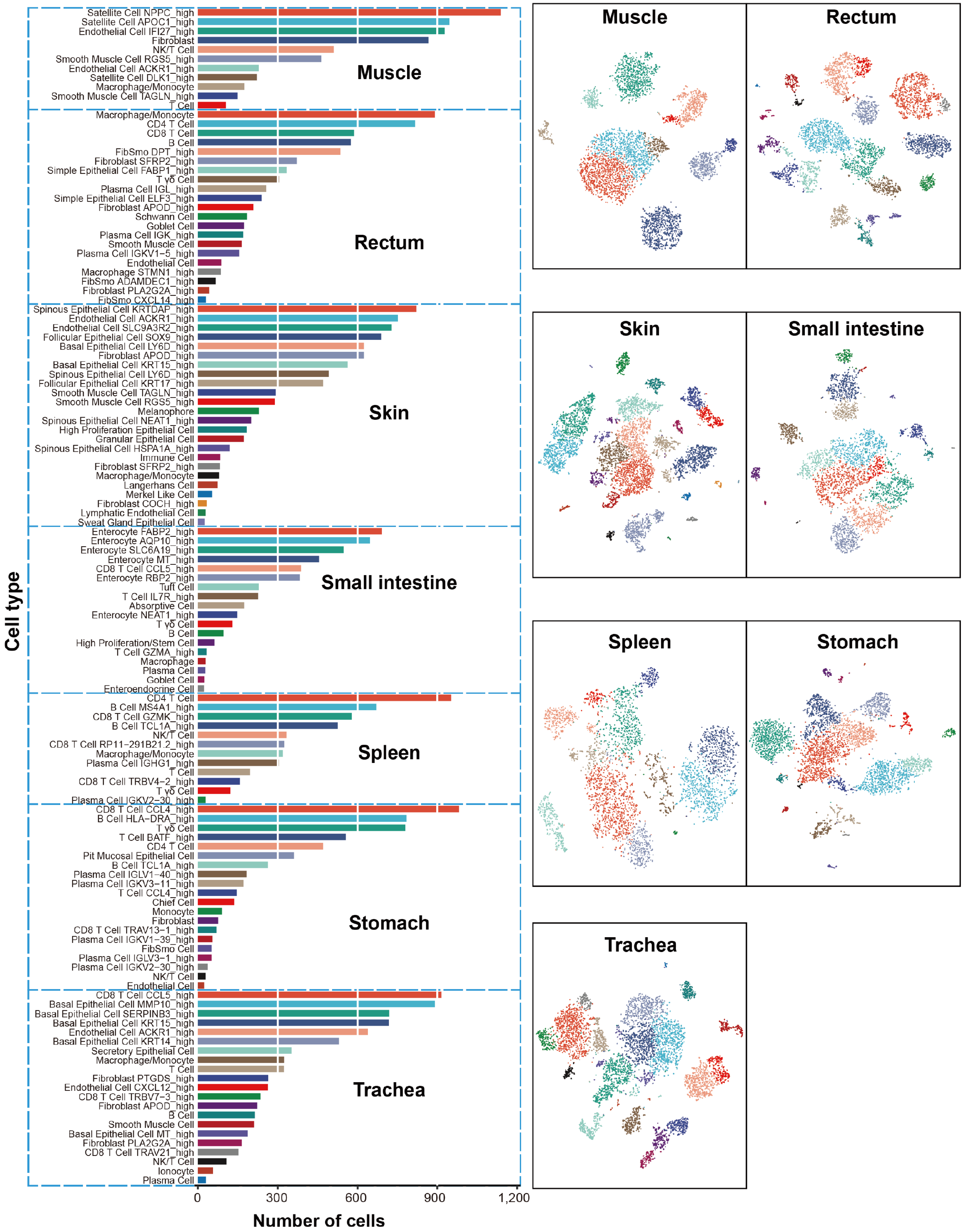


**Distribution of cell cluster in each organ tissue (continued).**

Appendix Figure S3

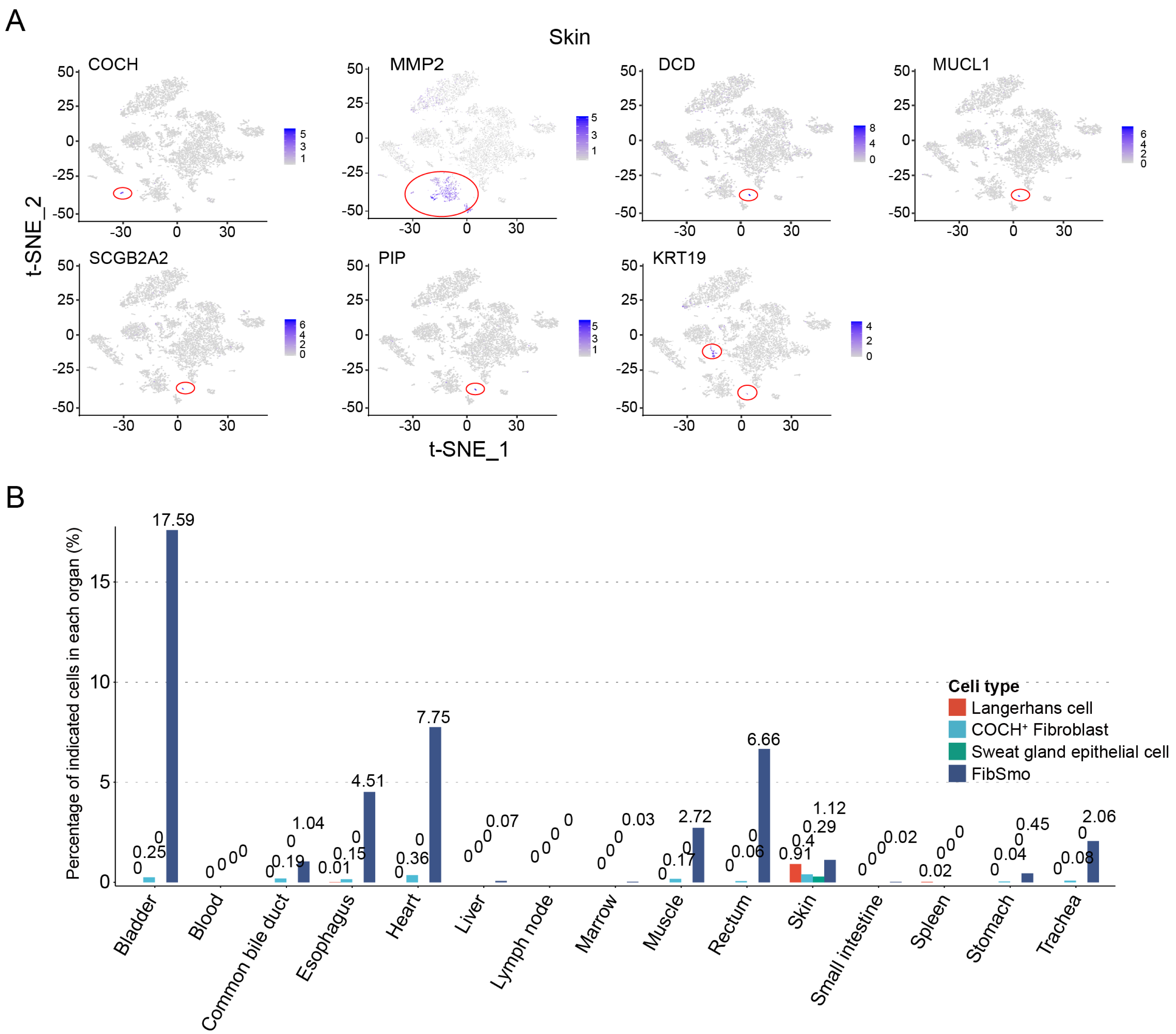


**The expression of marker genes and distribution of sweat gland epithelial cells and a novel fibroblast subtype, related to Figure 1.**

**A.** t-SNE plots show skin cells colored by the normalized expression of *COCH*, *MMP2*, *SCGB2A2*, *MUCL1*, *PIP*, *MUCL1*, and *DCD* genes. Each dot represents one cell. Color temperature represents the expression level.

**B.** Bar plot showing the distribution of sweat gland epithelial cells (with the co-expression of *SCGB2A2*, *KRT19*, *PIP*, *MUCL1*, and *DCD*), *COCH*⁺ fibroblasts (with the co-expression of *COCH* and *MMP2*), and FibSmo cells (with the co-expression of *ACTA2* and *MMP2*) in different organs indicated in the x-axis.

Appendix Figure S4

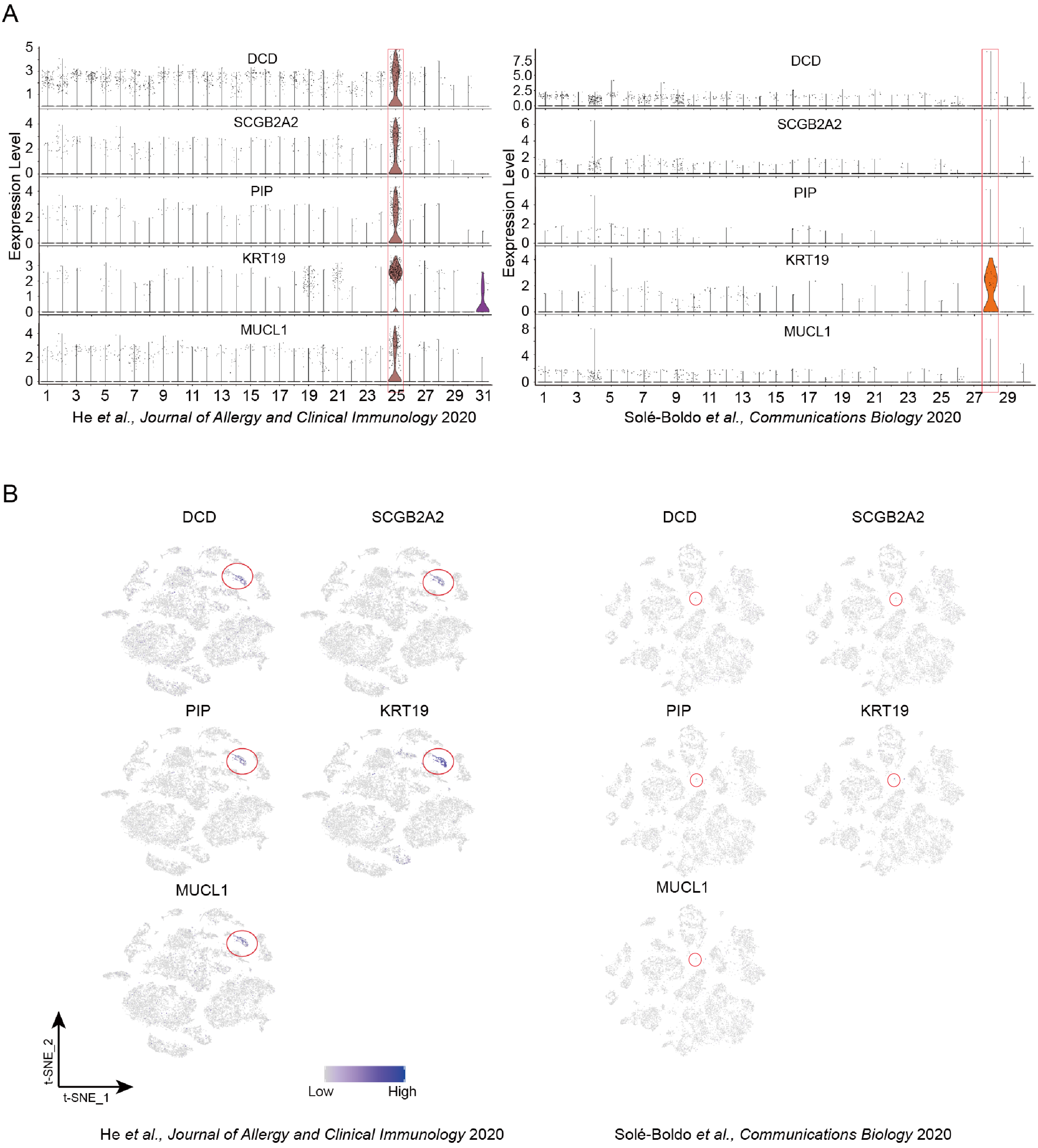


**Expression of *DCD*, *SCGB2A2*, *PIP*, *KRT19*, and *MUCL1* genes in skin cells derived from public datasets, related to Figure 1.**
**A.** Violin plots showing the normalized expression of *DCD*, *SCGB2A2*, *PIP*, *KRT19*, and *MUCL1* in cells of different clusters from the skin tissues retrieved from public datasets as referenced at the bottom. Clusters are numbered on the x-axis, and the expression level is shown on the y-axis. Each dot represents one cell. The clusters for *SCGB2A2*$^{+}$ sweat gland cells are boxed.
**B.** Feature plots showing the normalized expression of *DCD*, *SCGB2A2*, *PIP*, *KRT19*, and *MUCL1* in cells from skin tissues retrieved from public datasets as referenced at the bottom. The clusters for sweat gland cells are circled.

Appendix Figure S5

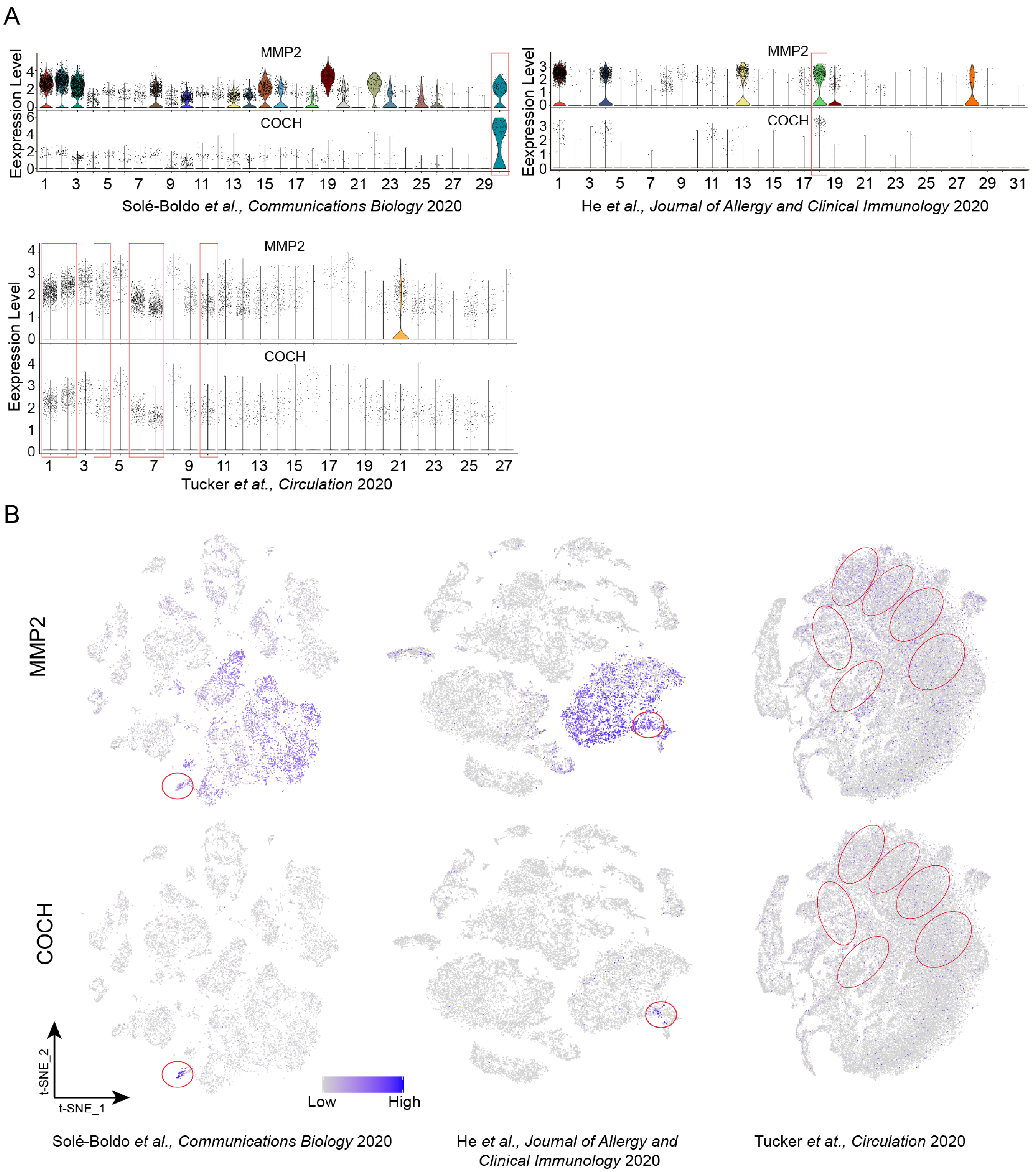


**Expression of *COCH* and *MMP2* in cells from multiple human samples from public datasets, related to Figure 1.**
**A.** Violin plots showing the normalized expression of *COCH* and *MMP2* in cells of different clusters from the skin (top) and heart (left bottom) tissues retrieved from public datasets as referenced at the bottom of each plot. Clusters are numbered on the x-axis, and the expression level is shown on the y-axis. Each dot represents one cell. The clusters for *COCH*$^{+}$*MMP2*$^{+}$ fibroblasts are boxed.
**B.** Feature plots show the normalized expression of *COCH* and *MMP2* in cells from the skin (left and middle) and heart (right) tissues retrieved from public datasets as referenced at the bottom. Each dot represents one cell. The clusters for *COCH*$^{+}$*MMP2*$^{+}$ fibroblasts are circled.

Appendix Figure S6

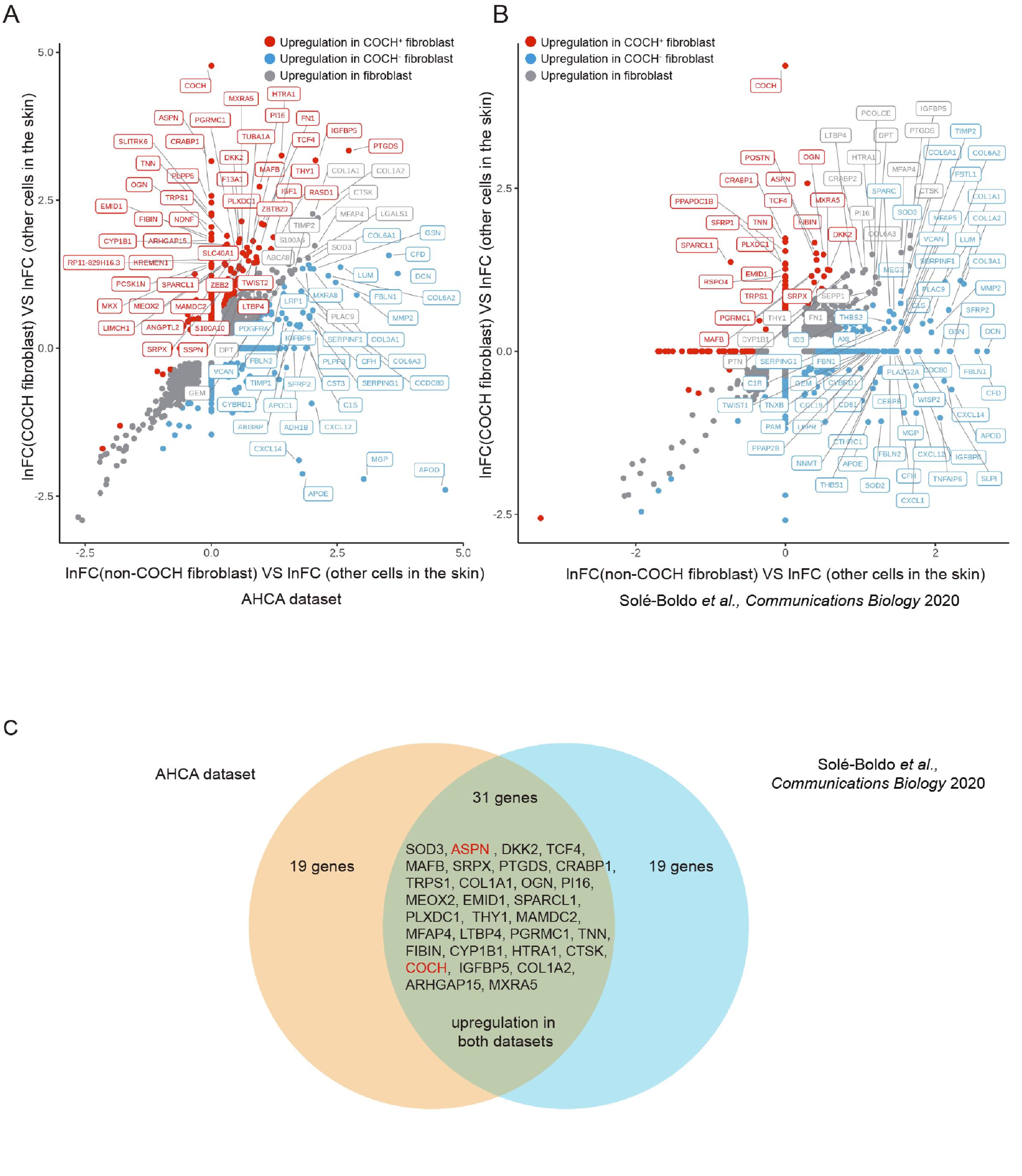


**Gene signature of *COCH*⁺ fibroblasts in skin, related to Figure 1.**
**A, B.** Scatter plots show the differentially expressed genes compared to other cells in skin samples from our AHCA or Solé-Boldo *et al.*'s datasets. Each dot represents one gene, which is labelled if it had ln(fold-change) greater than 4 and 3 for upregulation (in red), or smaller than 0.25 and 0.33 for downregulation (blue) in the AHCA and Solé-Boldo *et al.*'s datasets, respectively.
**C.** Venn diagram shows the sharing of upregulated genes in *COCH*⁺ fibroblasts in comparison to all other cells between two datasets.

Appendix Figure S7

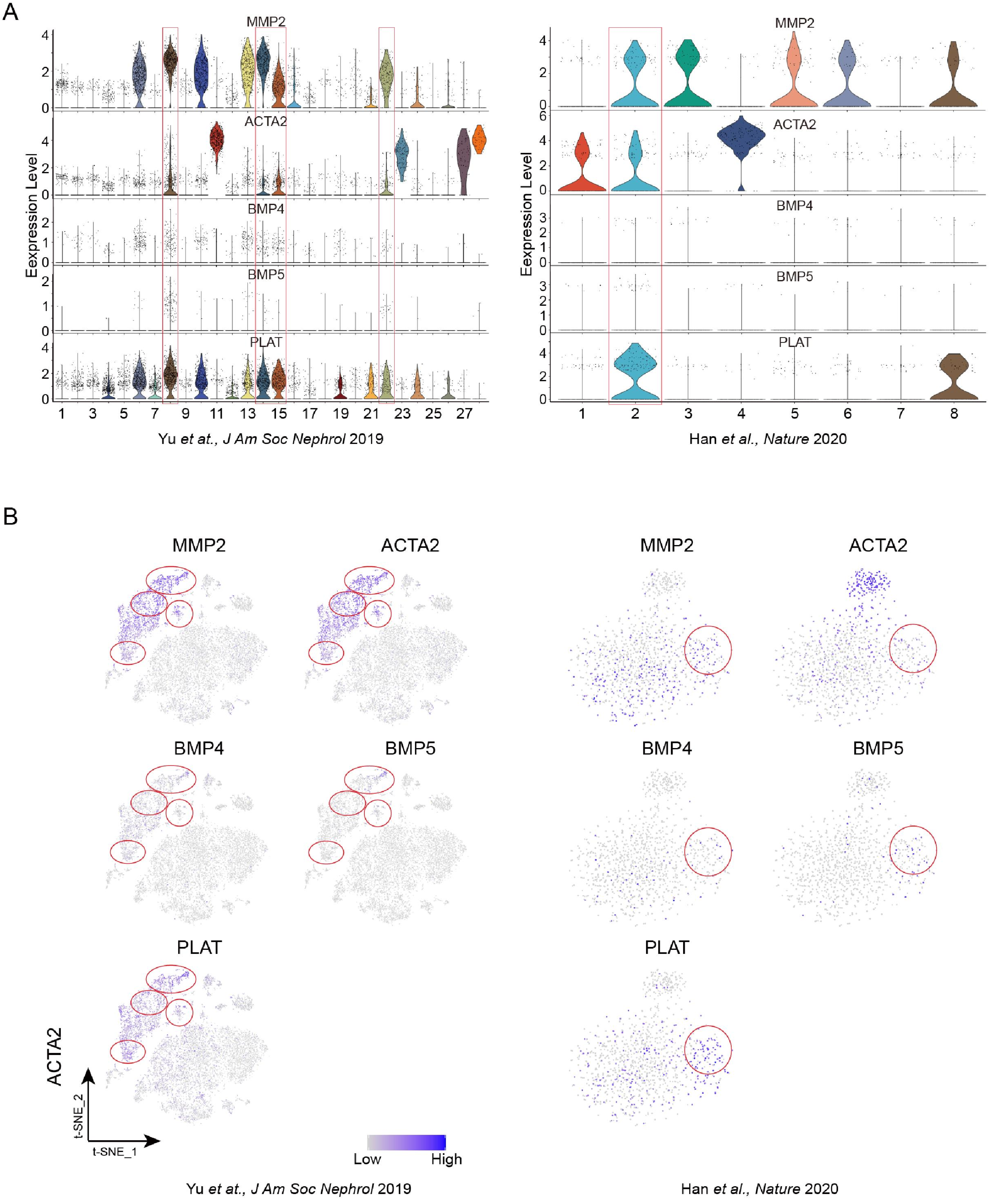


**Expression of marker genes in cells from bladder samples derived from public datasets, related to Figure 1.**

**A.** Violin plots show the normalized expression of *MMP2*, *ACTA2*, *BMP4*, *BMP5*, and *PLAT* in cells of different clusters from the bladder tissues retrieved from public datasets as referenced at the bottom. Clusters are numbered on the x-axis, and the expression level is shown on the y-axis. Each dot represents one cell. The clusters for FibSmo cells are boxed.

**B.** Feature plots showing the normalized expression of *MMP2*, *ACTA2*, *BMP4*, *BMP5*, and *PLAT* in cells from skin tissues retrieved from public datasets as referenced at the bottom. The clusters for FibSmo cells are circled.

Appendix Figure S8

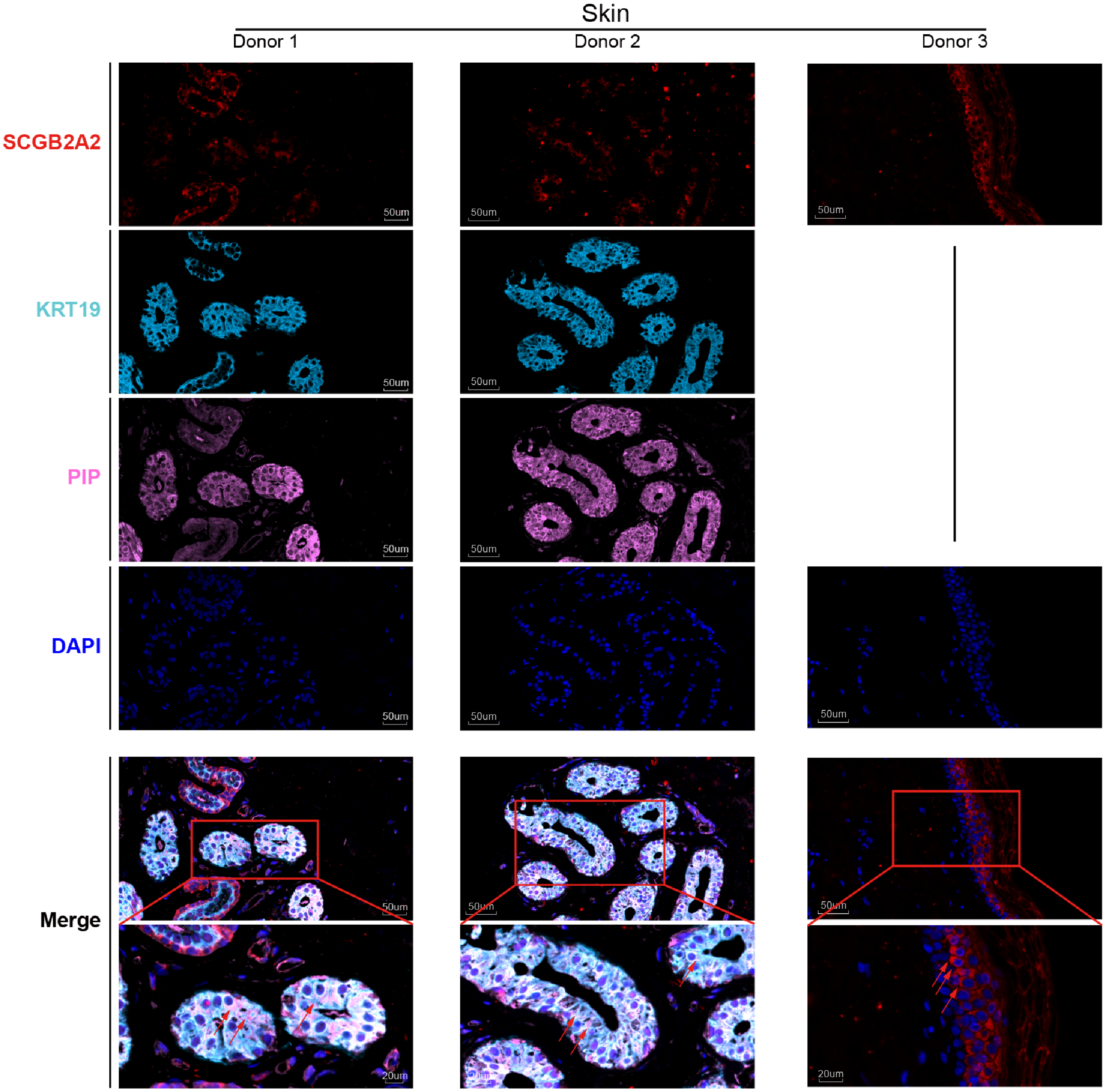


**Representative images of immunofluorescence staining of sweat gland epithelial cells in skin samples.**
Proteins used in the assays are indicated on the left, and the samples from different donors (1-3) are listed as columns. The red arrows indicate the co-expression of SCGB2A2, PIP, and KRT19 proteins in cells of interest in donors 1, and 2. Donor 3 served as a positive control of SCGB2A2 staining. The red arrows indicate the specific expression of SCGB2A2 in certain cell types of the skin from donor 3. Scale bar is as indicated in the image.

Appendix Figure S9

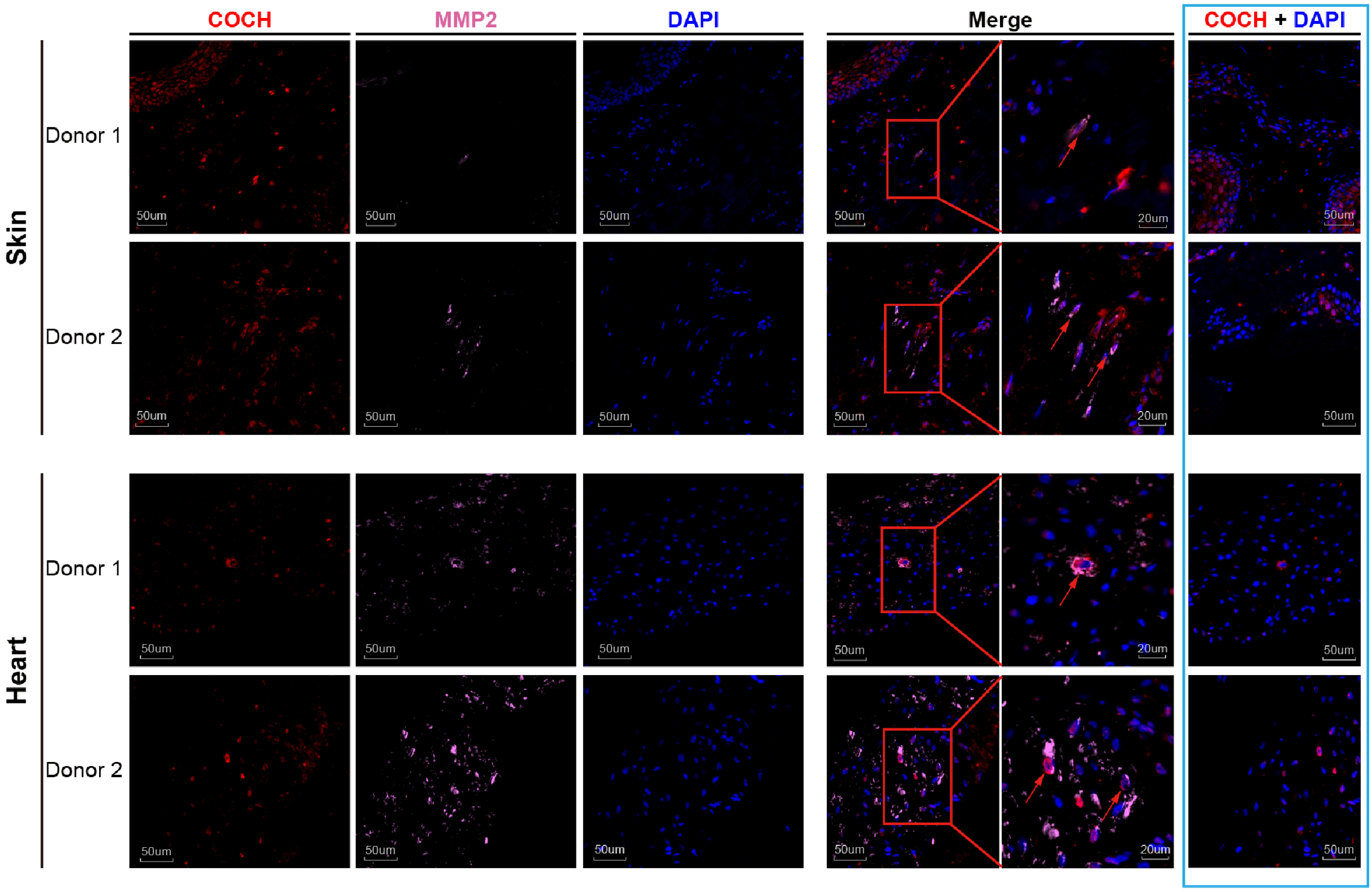


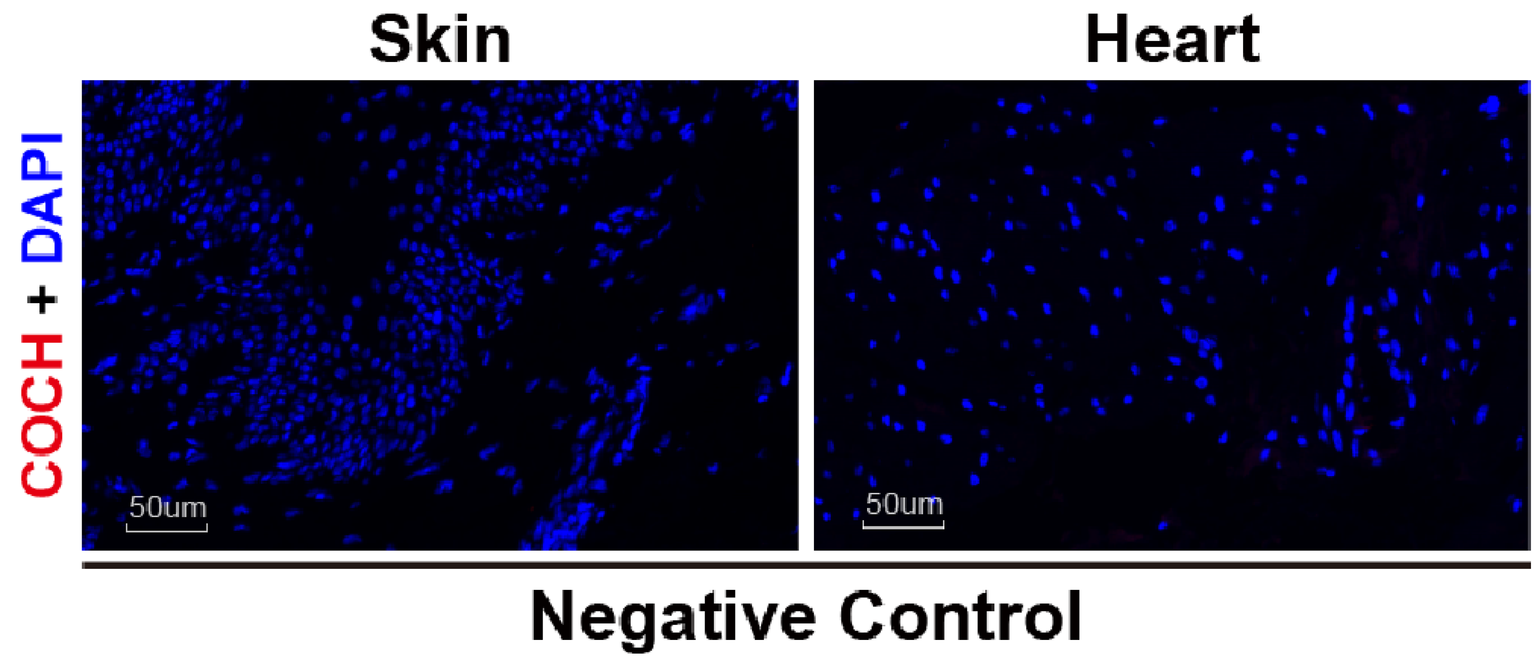


**Representative images of immunofluorescence staining of COCH⁺ fibroblasts in the skin and heart tissues.**
Proteins used in the assays are indicated on top, and the skin and heart samples from different donors (1-2) are listed as rows. The red arrows indicate the co-expression of COCH and MMP2 proteins in the cells of interest in tissue samples as indicated on the left. For each sample, the right panel highlighted in a box was imaged field different from the left panel on the same tissue slide, showing the co-existence of COCH negative and positive cells. Images on the bottom panel show the negative controls of COCH.

Appendix Figure S10

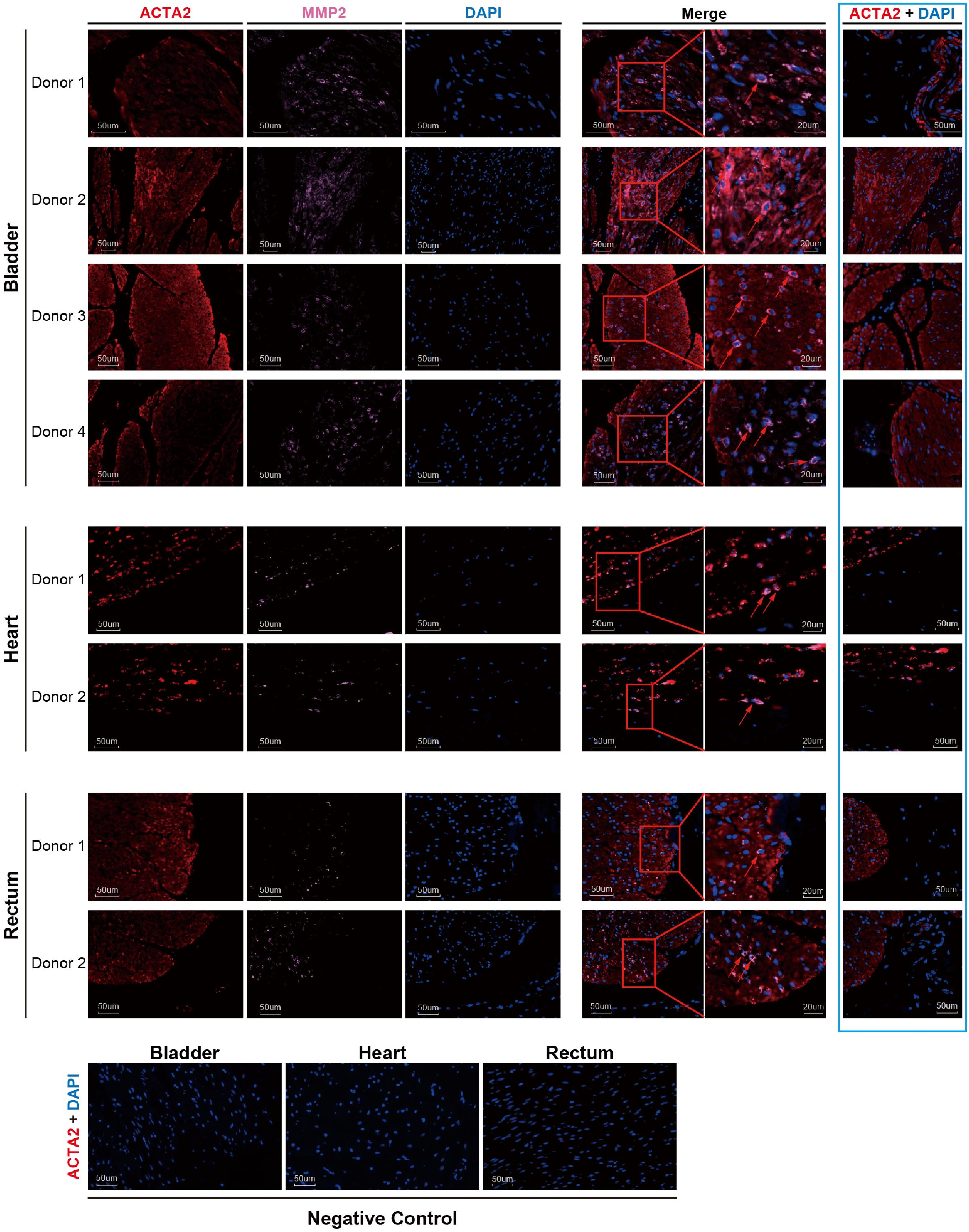


**Representative images of immunofluorescence staining of FibSmo cells in the bladder, rectum, and heart tissues.**
Proteins used in the assays are indicated on top, and the bladder, heart, and rectum samples from different donors (1-4) are listed as rows. The red arrows indicate the co-expression of ACTA2 and MMP2 proteins in the cell of interest in tissue samples indicated on the left. For each sample, the right panel highlighted in a box was imaged field different from the left panel on the same tissue slide, showing the co-existence of ACTA2 negative and positive cells. Images on the bottom panel show the negative controls of ACTA2.

Appendix Figure S11

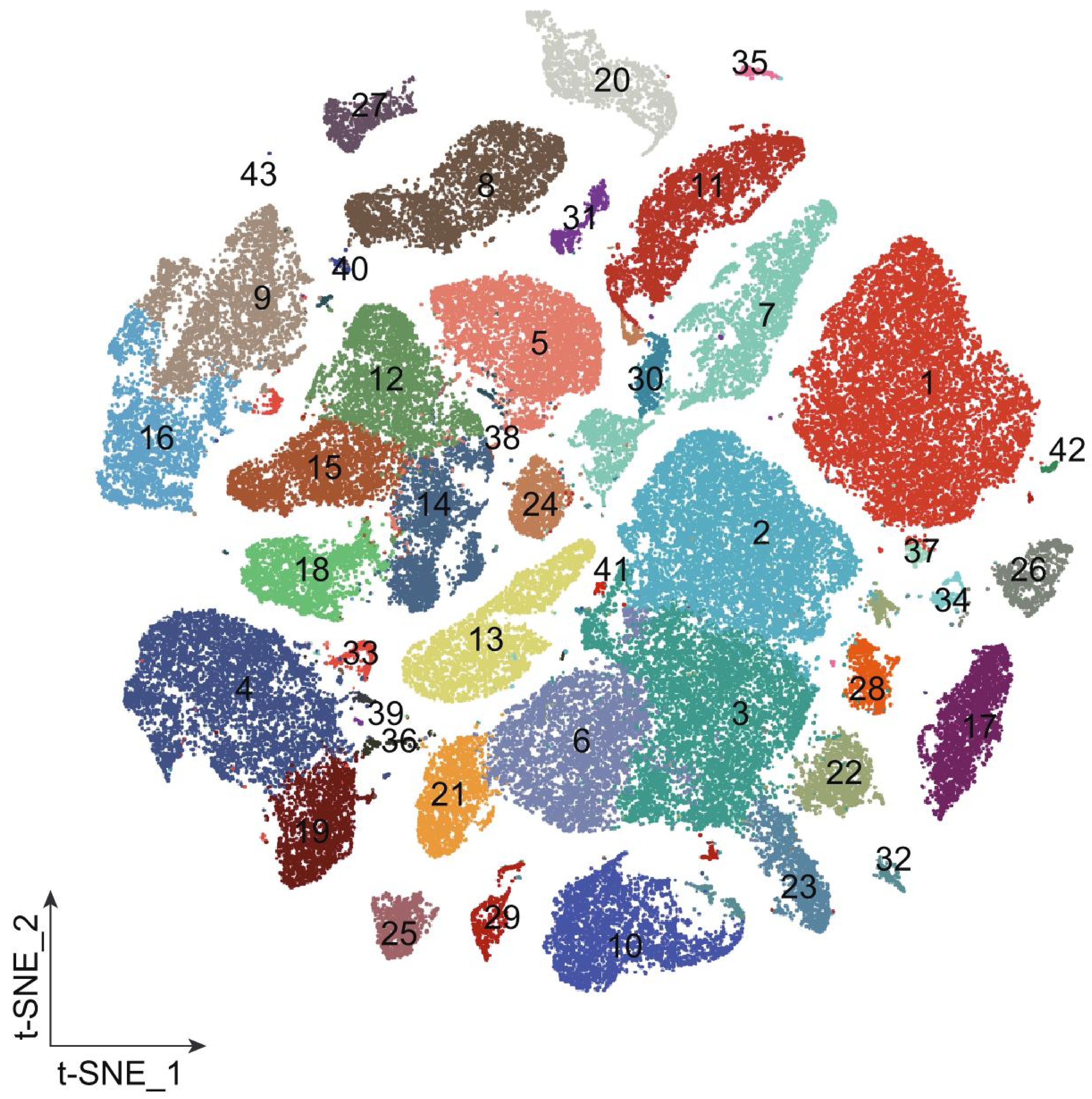


- 1 B Cell CD79A
- 2 T Cell IL7R
- 3 T Cell XCL1
- 4 Endothelial Cell ACKR1
- 5 Fibroblast PTN
- 6 T Cell GZMK
- 7 Keratinocyte KRT1
- 8 Smooth Muscle Cell
- 9 Macrophage C1QB
- 10 Basal Cell KRT17
- 11 Squamous Epithelial Cell KRT13
- 12 Fibroblast PLA2G2A
- 13 Enterocyte APOA1
- 14 Fibroblast PTGDS
- 15 Fibroblast C7
- 16 Monocyte
- 17 Satellite Cell
- 18 FibSmo Cell
- 19 Endothelial Cell FABP4
- 20 Plasma Cell JCHAIN
- 21 NK/T Cell GNLY
- 22 T Cell CCL5
- 23 T Cell GZMA
- 24 Squamous Epithelial Cell KRT15
- 25 Fibroblast APOD
- 26 Epithelial Cell TFF3
- 27 Cholangiocyte
- 28 T Cell RGS1
- 29 Mucosal Epithelial Cell
- 30 Keratinocyte KRT14
- 31 Melanocyte
- 32 Secretory Cell
- 33 Macrophage FCN3
- 34 Absorptive Cell
- 35 Tuft Cell
- 36 T Cell PRF1
- 37 B Cell MS4A1
- 38 Fibroblast GSN
- 39 Lymphatic Endothelial Cell
- 40 Macrophage RNASE1
- 41 High Proliferation Erythrocyte
- 42 B Cell VPREB3
- 43 Plasma Cell IGKV1-12

**Distribution of cell clusters in different organs, related to Figure 1.**
t-SNE plot showing cell clusters in 15 organs (84,363 cells). Each dot represents one cell, which all together form clusters and color-coded according to the cluster names listed at the bottom. The rare or novel cells identified above are grouped into different clusters based on their genetic profiles. Langerhans cells, *COCH*[+] fibroblasts, and sweat gland epithelial cells were grouped in the cluster 9 "Macrophage C1QB", the cluster 14 "Fibroblast PTGDS_high", and the cluster 32 "Secretory Cell", respectively. FibSmo cells were grouped in cluster 18 "FibSmo cell".

Appendix Figure S12

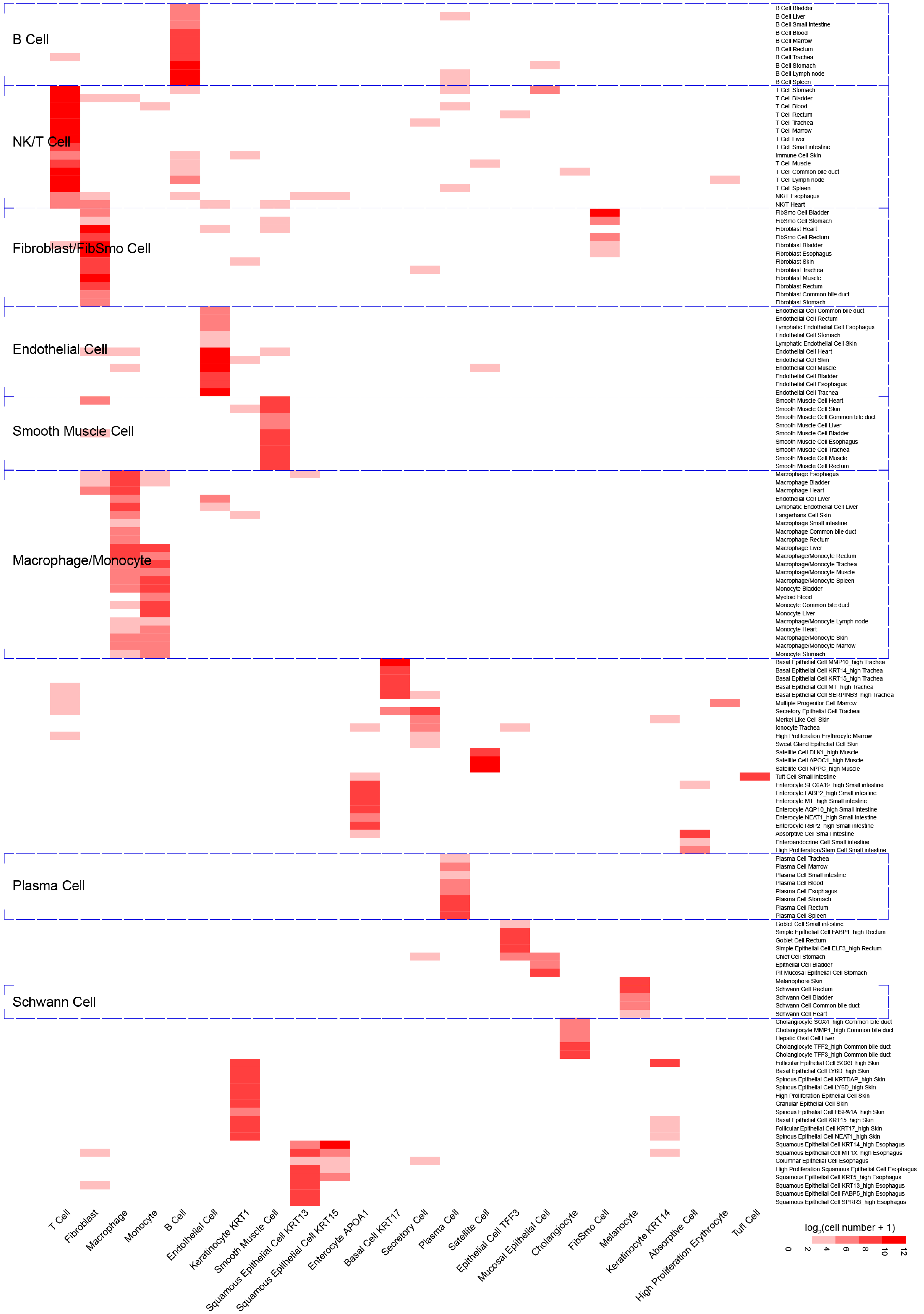


**Comparisons of cell-type determinations by whole single-cell transcriptome dataset and manual annotation of each organ, related to Figure 1.** The x-axis indicates clusters from the main **Figure 1B**, and the y-axis indicates clusters annotated manually, which were determined using single-cell transcriptomes from all 15 tested organs and the expression profile of canonical marker genes in the cells from individual organ, respectively. The results indicate that similar cell types from distinct organs were clustered into a major cell type. In particular T cells, NK cells, B cells, plasma cells, endothelial cells, fibroblasts, smooth muscle cells, macrophages, monocytes, and Schwan cells from different organs were grouped into limited clusters (regions outlined in dashed boxes). Heatmap is colored according to the $\log_2$(cell number +1) of sharing cells.

### Appendix Figure S13

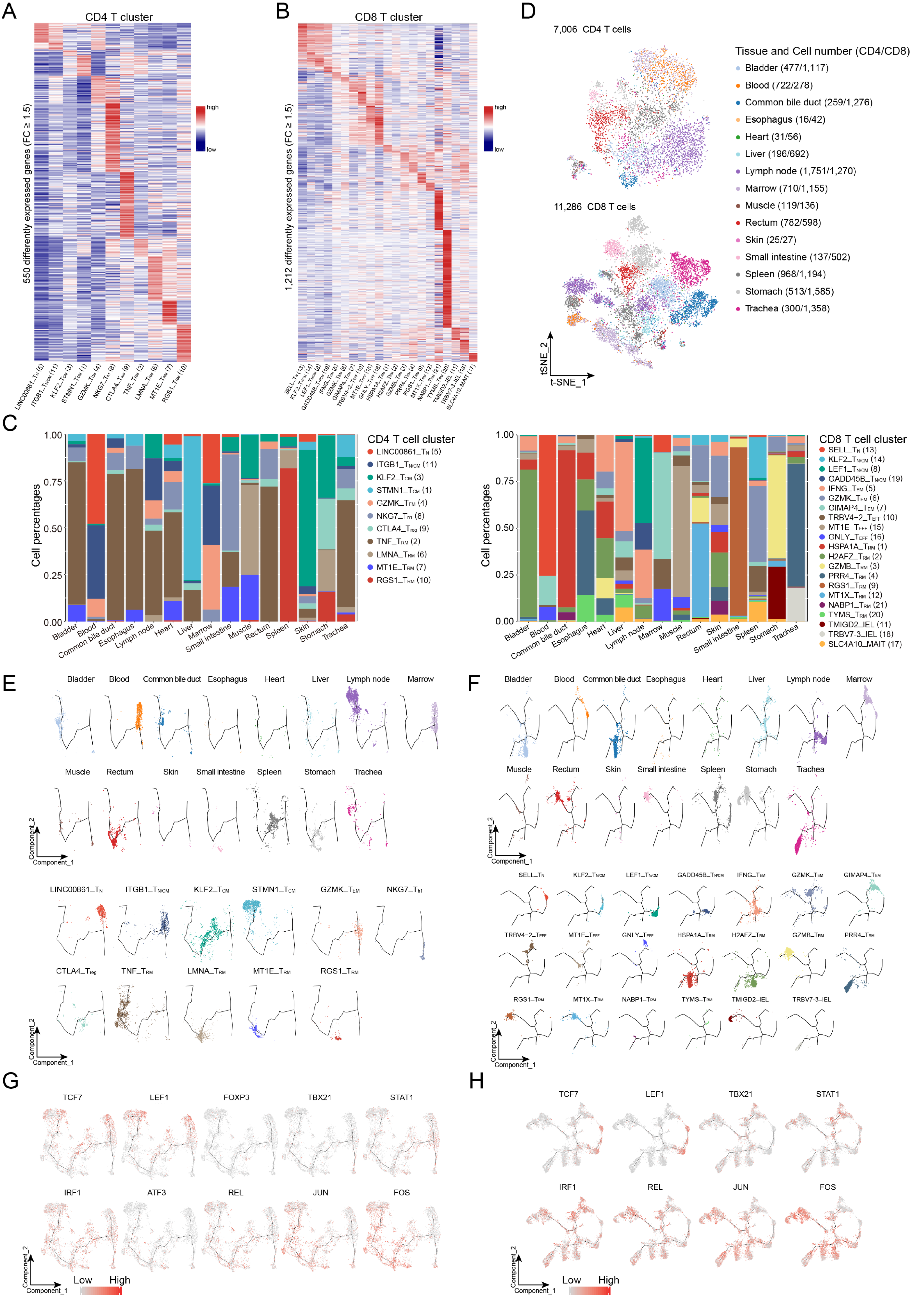

**The heterogeneity and developmental trajectory analysis of T cells, related to Figure 2.**

**A, B.** A heat map showing the expression profiles of each $CD4^+$ (**A**) and $CD8^+$ (**B**) T cell clusters as indicated at the bottom. The expression of 550 and 1,212 genes with FC ≥ 1.5, pct.1 ≥ 0.2 and *p.adjust* < 0.05 are shown as lines, colored from blue to red according to the expression from low to high, for $CD4^+$ (**A**) and $CD8^+$ (**B**) T cell clusters, respectively.

**C.** Bar charts show the proportions of cell types in each tissue for $CD4^+$ (left panel) and $CD8^+$ T cells (right panel). For each tissue at the x-axis, proportions of each cell cluster indicated on the right are shown in the y-axis.

**D.** t-SNE plot of 7,006 $CD4^+$ (top panel) and 11,256 $CD8^+$ T cells (bottom panel) from the 15 organ tissues. Each point represents one cell, color-coded according to their origins of tissues as indicated on the right.

**E, F.** Projection of each group of T cells on the pseudo-time trajectory tree of all $CD4^+$ (**E**) and $CD8^+$ T cells (**F**) according to cell type or origins of tissues. Each dot represents one cell and are color-coded according to their origin of tissue (top panels) or cluster (bottom panels).

**G, H.** The normalized expression of representative TFs in cells along the pseudo-time trajectory trees in **Figure 2F** and **G** for $CD4^+$ (**G**) and $CD8^+$ (**H**) T cell, respectively. Each dot represents one cell with expression from grey to red, indicating low and high expression, respectively. *TCF7* and *LEF1* for $CD4^+$, $CD8^+$ $T_N$, $T_{N/CM}$, and $T_{CM}$ cells; *FOXP3* for $CD4^+$ $T_{reg}$ cells; *TBX21*, *STAT1*, and *IRF1* for both $CD4^+$ ($T_{h1}$) and $CD8^+$ $T_{EFF}$ cells; enhanced expression of *ATF3* for $CD4^+$ $T_{RM}$ cells; enhanced expression of *FOS*, *JUN*, and *REL* for both $CD4^+$ and $CD8^+$ $T_{RM}$ cells.

Appendix Figure S14

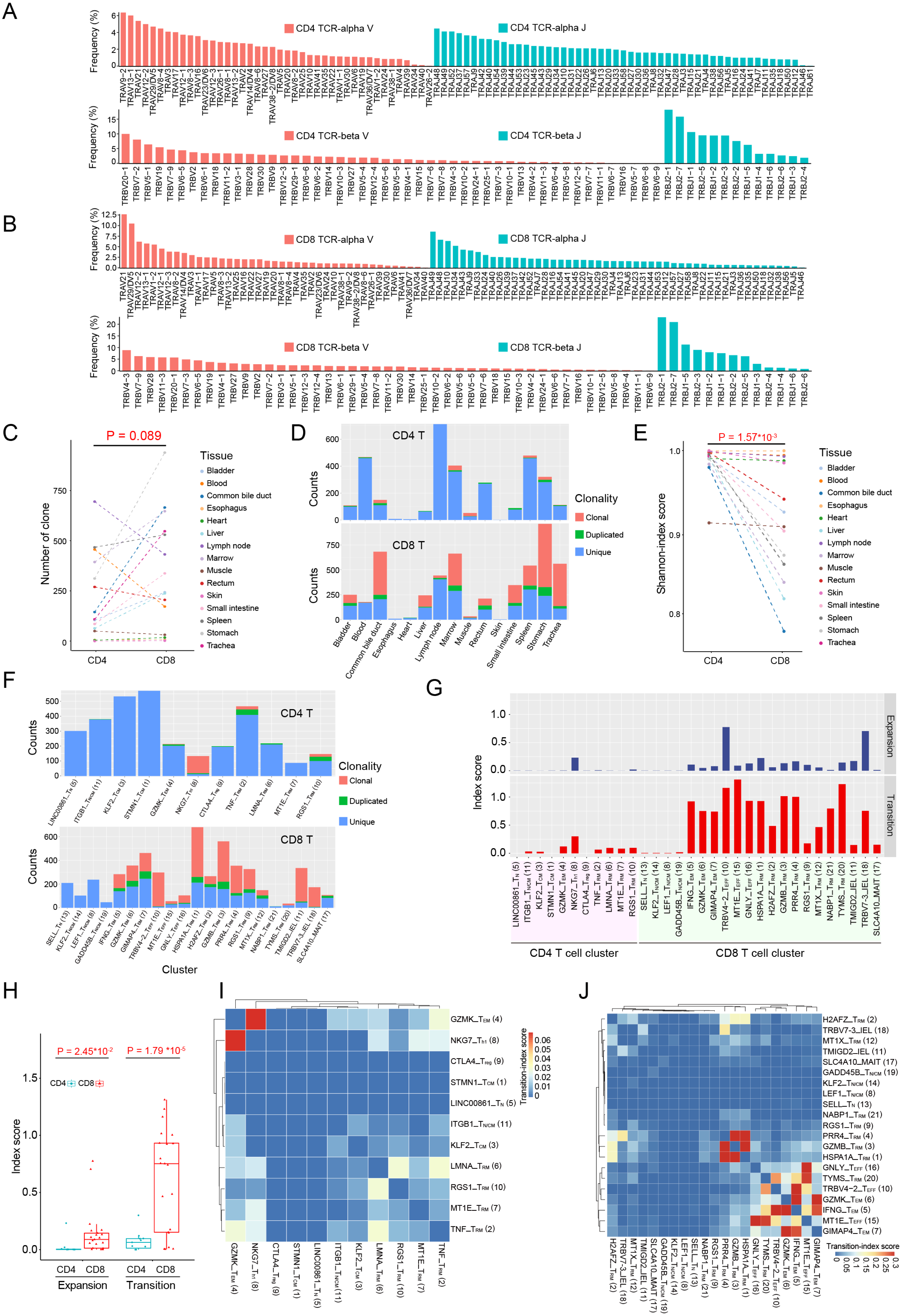

**The heterogeneity and clonalities of T cell clones in human body, related to Figure 2.**

**A, B.** Frequencies of the V and J segments of the TCR α (top panels) and β chains (bottom panels) for $CD4^+$ (**A**) and $CD8^+$ T cells (**B**). The x- and y- axes indicate the gene segment and frequency of usage, respectively.

**C.** Comparison of TCR clone sizes between $CD4^+$ and $CD8^+$ T cells in each tissue as indicated by different colors. Two-sided paired Student's t-test was conducted.

**D.** Bar plots showing the number of clonal cells in each organ for $CD4^+$ (top panel) and $CD8^+$ (bottom panel) T cells. The clonotypes are categorized as unique ($n = 1$), duplicated clonal ($n = 2$) and multiple clonal ($n \geq 3$) based on their numbers of cells sharing a same TCR.

**E.** Comparison of diversity between $CD4^+$ and $CD8^+$ T cells in each organ using Shannon entropy. A two-sided paired Student's t-test was conducted.

**F.** Bar plots showing the numbers of clonal cells in each cell cluster for $CD4^+$ (top panel) and $CD8^+$ (bottom panel) T cells. The definitions for clonotypes are as same as in **D**.

**G.** Expansion- (top panel) and transition-index (bottom panel) scores of each $CD4^+$ and $CD8^+$ T cell cluster calculated using STARTRAC. Cell clusters are indicated at the bottom.

**H.** Box plot showing the expansion- and transition-index scores of $CD4^+$ or $CD8^+$ T cells. The comparison was tested using a Wilcoxon rank-sum test.

**I, J.** Heat maps showing the pairwise transition-index scores for the $CD4^+$ (**I**) and $CD8^+$ T cell (**J**) clusters. Clusters are shown on the right and the bottom. Squares are filled with colors from blue to red representing the transition-index scores from low to high. Values were truncated to 0.3 for $CD8^+$ T cells.

Appendix Figure S15

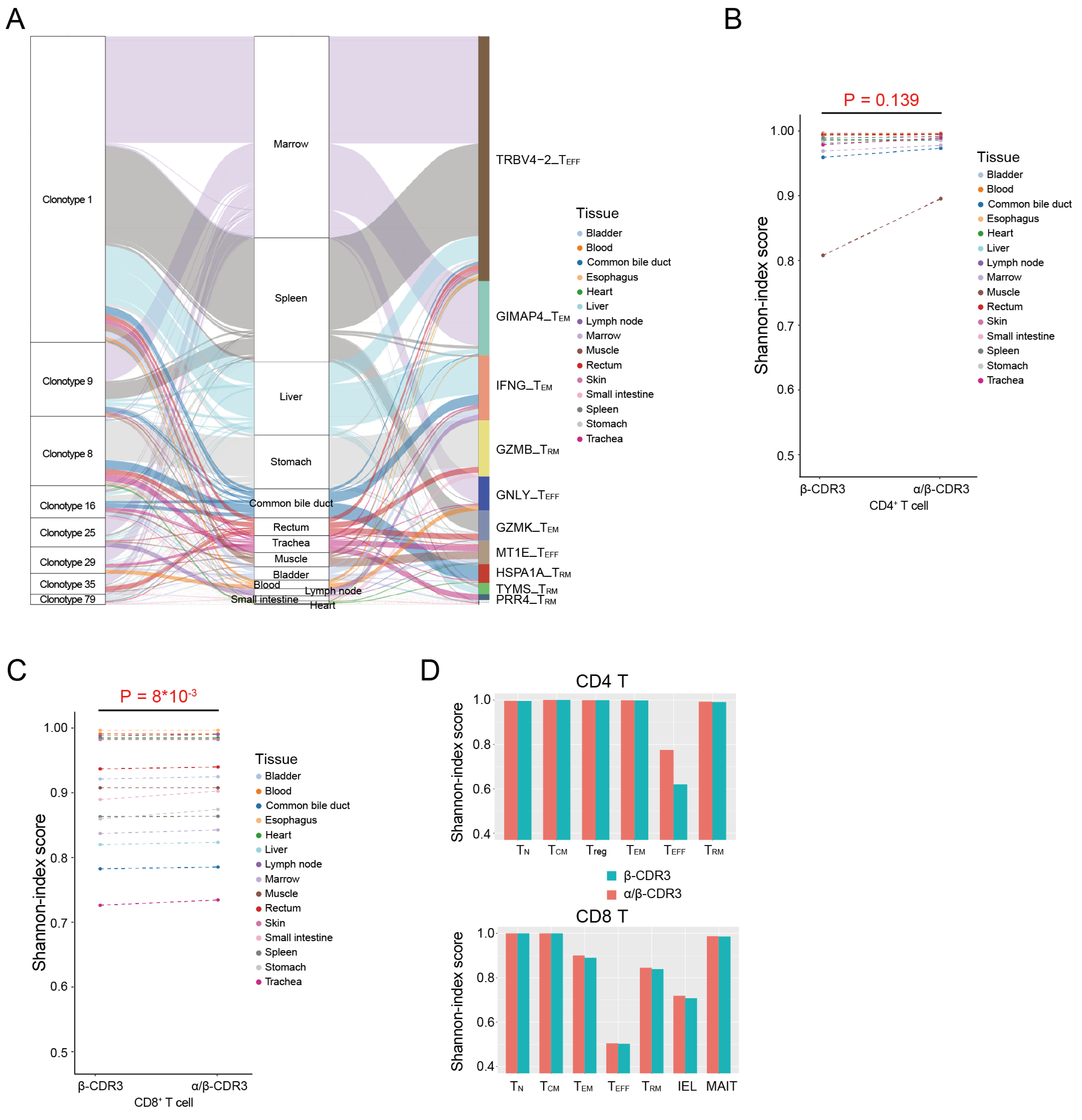


**The heterogeneity and clonalities of T cell clones in the human body, related to Figure 2.**

**A.** An alluvial diagram showing CD8⁺ T cells with the same TCRs detected in at least seven tissues and different cell types. Each line represents the sharing of a cell with specific clonotype labeled at the left box, colored by tissue as indicated on the right.

**B, C.** Diversities of CD4⁺ (**B**) and CD8⁺ (**C**) T cells in each organ. Shannon-index score was measured using TCR β chain or paired TCR α-β chains. The organs are listed on the right. Comparison was tested with a two-sided paired Student's t-test.

**D.** Diversities of subtypes of CD4⁺ (top panel) and CD8⁺ (bottom panel) T cell clusters (x-axis). Shannon-index score was measured using TCR β chain or paired TCR α-β chains.

### Appendix Figure S16

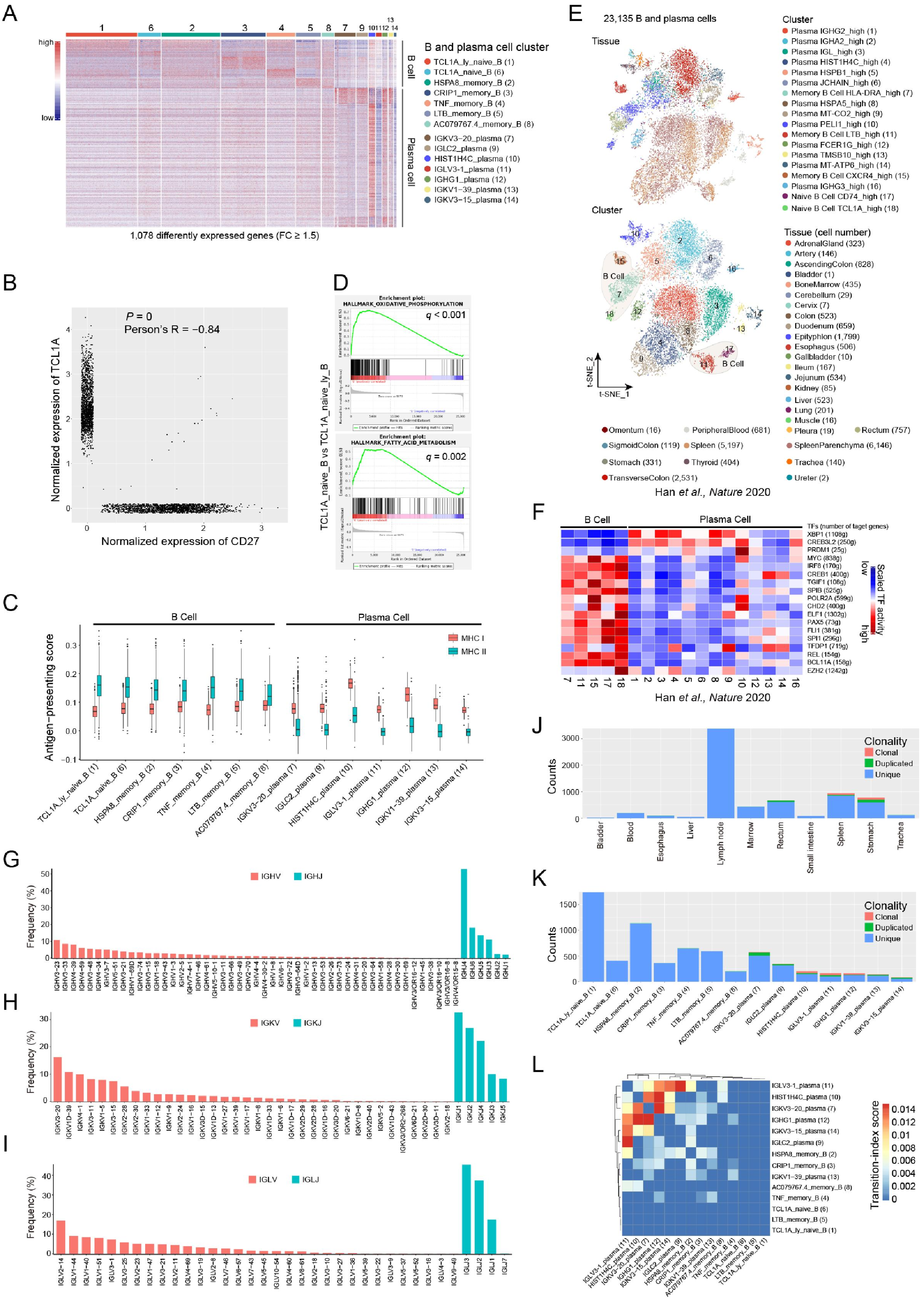

**The heterogeneity of B and plasma cells, related to Figure 3.**

**A.** Expression profiles of each B and plasma cell clusters as indicated by the numbers on top and illustrated at the right inlet box. The expression of 1,078 genes with FC ≥ 1.5, pct.1 ≥ 0.2 and *p.adjust* < 0.05 are shown as colored lines according to their expression levels and IGK/L/H genes were removed.

**B.** A scatter plot showing the reverse correlation between the expressions of *CD27* and *TCL1A* in all B and plasma cells. Cells with no expression of both the genes were excluded. Person's R = -0.84, *P* = 0.

**C.** Box plot showing the antigen-presenting scores for intra- (MHC I) and extracellular (MHC II) antigens of each B and plasma cell cluster as indicated on the x axis.

**D.** GSEA analysis results showing the significant enriched pathways of '*oxidative phosphorylation'* and '*fatty acid metabolism'* in the naïve B in non-lymph node anatomical position.

**E.** t-SNE plots showing 18 clusters (23,135 cells) of B and plasma cells colored by tissue (top panel) and subtype (bottom panel) in the HCL dataset. Each dot represents one cell.

**F.** Heatmap showing the activation scores of TFs of each B and plasma cell subtypes in the HCL dataset. Cell clusters are indicated at the bottom, and the scores were estimated using SCENIC analysis. The top 10 TFs selected with the highest difference in activation scores estimates between each cluster and all other cells are shown.

**G-I.** Frequencies of V and J gene segments of IGH (**G**), IGK (**H**), and IGL (**I**) chains. The x- and y- axes indicate the gene segment and frequency of usage, respectively.

**J, K.** Bar plots showing the numbers of clonal cells in each organ (**J**) and cell cluster (**K**). The clonotypes are categorized as unique (n = 1), duplicated clonal (n = 2) and multiple clonal (n ≥ 3) based on their numbers of cells sharing a same BCR.

**L.** Heat maps showing the pairwise transition-index scores for the B and plasma cell clusters. Clusters are shown on the right and the bottom. Squares are filled with colors from blue to red representing the transition-index scores from low to high.

Appendix Figure S17

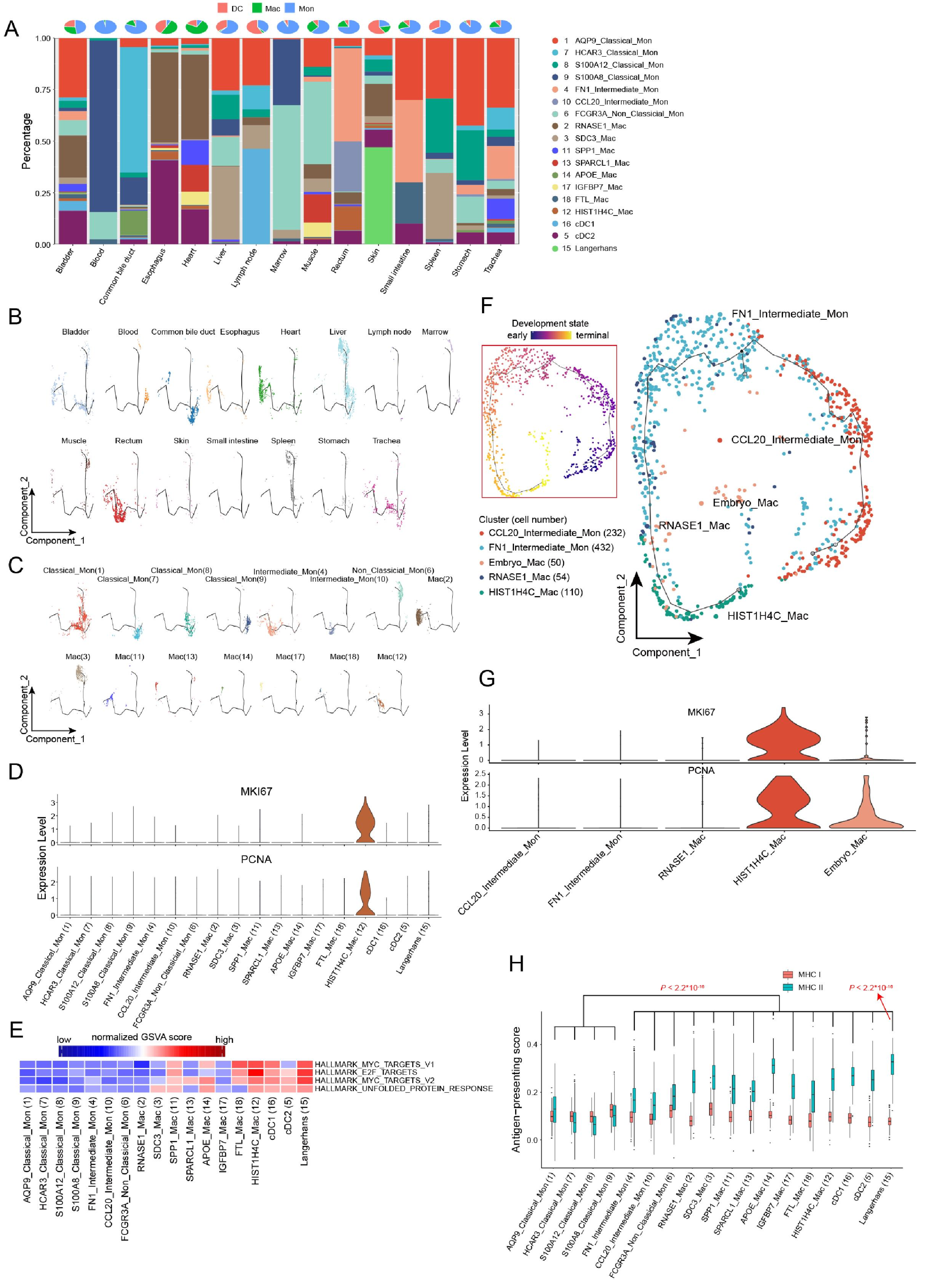

**Heterogeneity and trajectory analysis of macrophages, related to Figure 4.**

**A.** The distribution of myeloid cell subpopulations in each tissue. Pie charts on top illustrate the composition of macrophages, monocytes, and DC cells in each tissue. The stacked bars represent the percentage of each cluster (on the right) in the organ indicated at the bottom.

**B, C.** Pseudo-time trajectory analysis of myeloid cells in each tissue (**B**) or cluster (**C**). Each dot represents one cell.

**D.** Violin plots showing the normalized expression of proliferated marker genes (*MKI67* and *PCNA*) in each myeloid cell cluster as indicated on the x axis.

**E.** GSVA analysis showing the pathways with significant difference of activation between monocytes and macrophages clusters indicated at the bottom. Activations indicated as boxes in the rows were determined using GSVA and comparison was tested with an R package limma.

**F.** Pseudo-time trajectory analysis of myeloid cells in the intestinal macrophages derived from our and Gao *et al*.'s datasets. Each dot represents one cell and was colored. The inlet t-SNE plot shows each cell with a pseudo-time score from dark blue to yellow, indicating early and terminal states, respectively.

**G.** Violin plots showing the normalized expression of proliferated marker genes (*MKI67* and *PCNA*) in each myeloid cell cluster as indicated on the x axis.

**H.** A box plot showing the antigen-presenting scores for intra- (MHC I) and extracellular (MHC II) antigens of each myeloid cell cluster. Comparisons were done using a Wilcoxon rank-sum test.

Appendix Figure S18

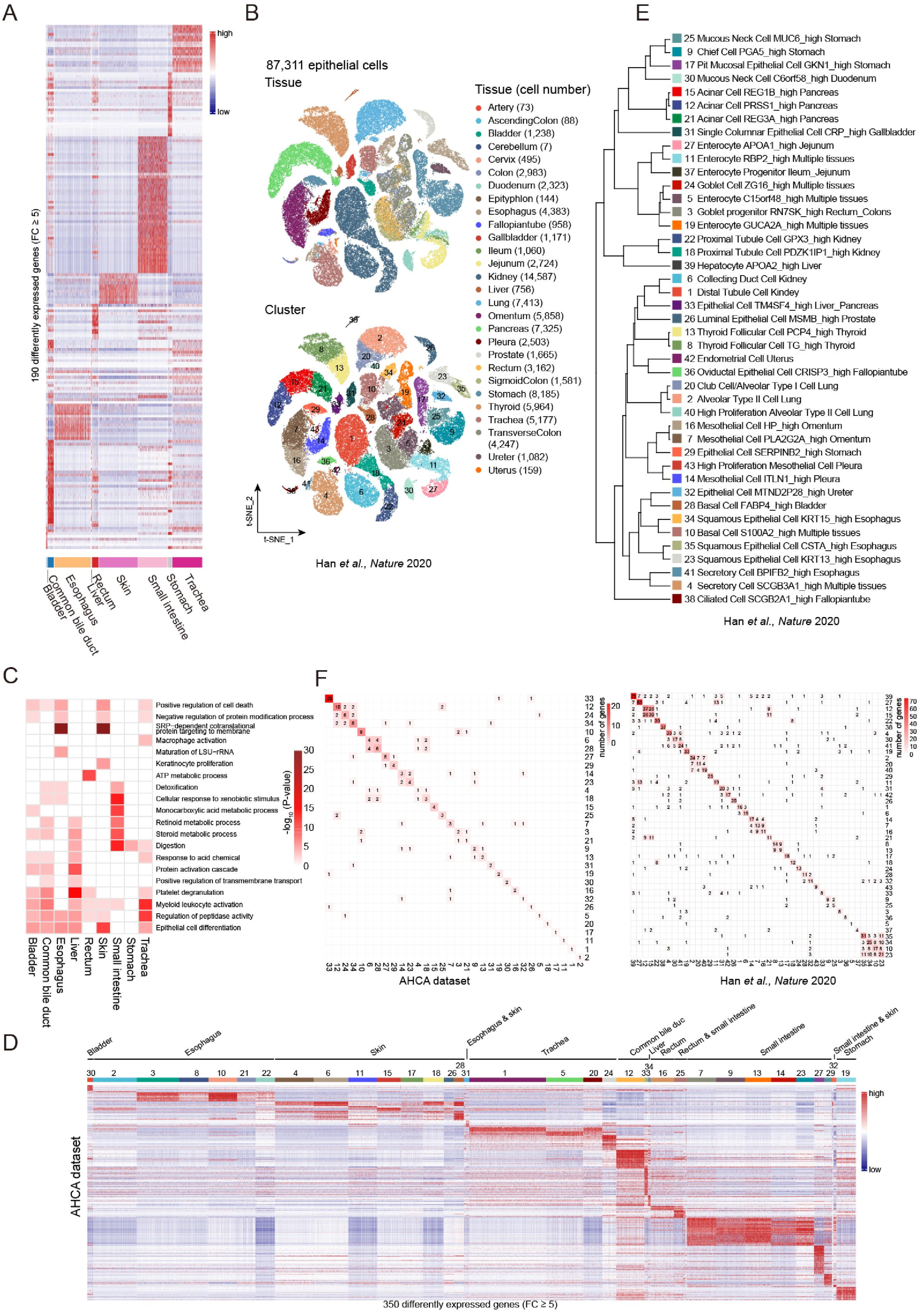

**The gene expression heterogeneity of epithelial cells among inter- and intra-organ tissues, related to Figure 5.**

**A.** A scaled heatmap showing the expression of 190 tissue-specific genes in epithelial cells from different tissues as indicated at the bottom. The tissue-specific genes passing FC ≥ 5, pct.1 ≥ 0.2 and *p.adjust* < 0.05 are shown in lines, colored with blue to red representing the expression levels from low to high.

**B.** t-SNE plots showing 87,311 epithelial cells, colored according to their origins of tissues (top panel) or cell clusters (bottom panel) in the HCL dataset. Each dot represents one cell.

**C.** Heatmap showing the top 20 pathways enriched significantly according to the GO analysis ($p < 0.05$) in epithelial cell from each organ as indicated at the bottom. The depth of color represents $-\log_{10}P$ calculated by the online tool "metascape".

**D.** A scaled heat map showing the expression of 350 signature genes in epithelial cell clusters as indicated by numbers on top, which refer to those in **Figure 5C**. The signature genes were determined by FC ≥ 5, pct.1 ≥ 0.2, pct.2 ≤ 0.2 and *p.adjust* < 0.05 and shown in lines, colored with blue to red representing the expression levels from low to high.

**E.** A dendrogram of 43 clusters based on their normalized mean expression values (correlation distance metric, complete linkage) in the HCL dataset. Only genes with ln(fold-change) above 0.25, *p.adjust* < 0.05 and pct.1 ≥ 0.2 in each cluster were included in the analysis.

**F.** Heatmaps showing the number of cluster-specific genes shared between clusters in our AHCA dataset (left panel) and in the HCL dataset (right panel). Clusters in numbers are indicated at the right and the bottom, which refer to those in **Figure 5C** and **Figure S18E**. The number in the boxes represent the amount of shared cluster-specific genes between two clusters and the color from white to red represents the number of shared genes from small to large.

Appendix Figure S19

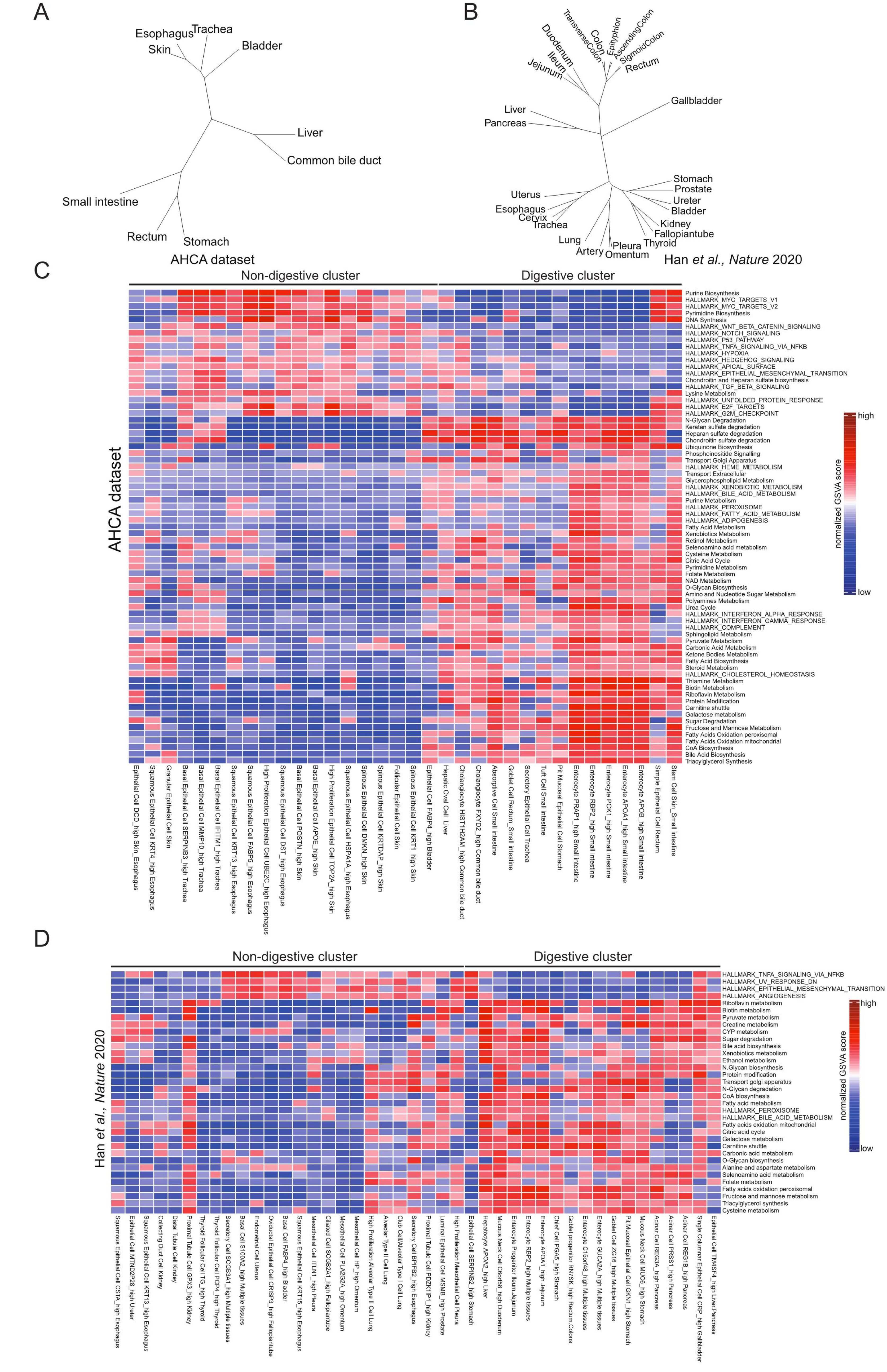

**Gene expression profiles and altered pathways in digestive and non-digestive epithelial cell clusters.**

**A, B.** Phylogenetic tree of tissues in AHCA (**A**) and HCL (**B**) datasets, based on their normalized mean expression values of epithelial cells (correlation distance metric, complete linkage). Only genes with ln(fold-change) above 0.25, *p.adjust* < 0.05 and pct.1 ≥ 0.2 in each organ were included in the analysis. Organs with less than 50 cells were excluded.

**C, D.** GSVA analysis showing the pathways (hallmarker and curated metabolic pathways) with significantly different activation between digestive and non-digestive cell clusters in the AHCA (**C**) and HCL (**D**) datasets. The x- and y- axis indicate a cell cluster in a tissue and enriched pathway, respectively. Activation scores indicated as boxes in the rows were determined using GSVA and comparison was tested with an R package limma.

Appendix Figure S20

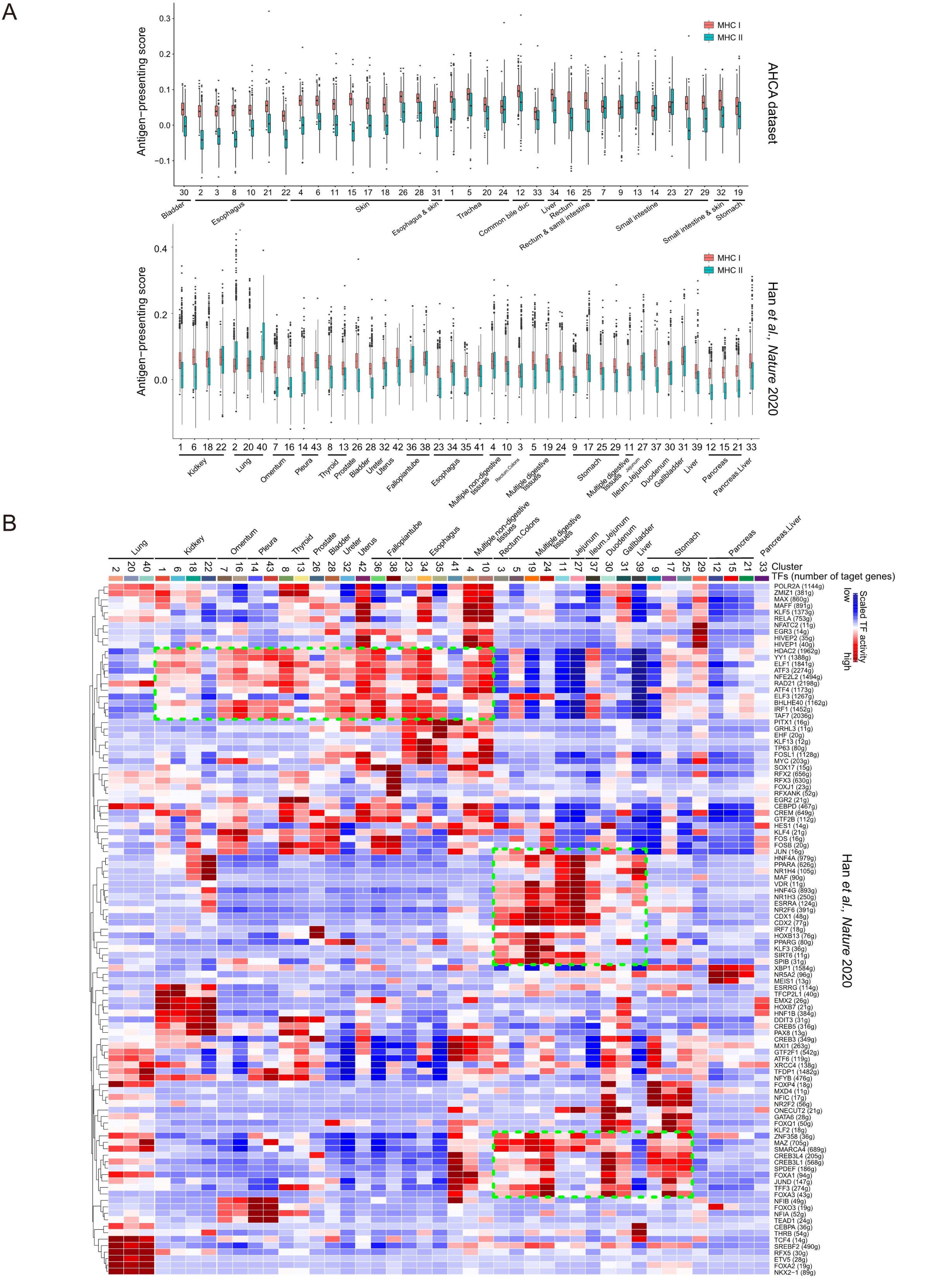

**The antigen-presenting heterogeneity and regulons of epithelial cells among inter- and intra-organ tissues, related to Figure 5.**

**A.** Box plots showing the antigen-presenting scores for intra- (MHC I) and extracellular (MHC II) antigens of each epithelial cell cluster as indicated on the x axis in AHCA (top panel) and HCL dataset (bottom panel). The clusters types refer to those in **Figure 5C** and **Figure S18E**.

**B.** Heatmap showing the activation scores of epithelial cell subtypes in the HCL dataset. Cell clusters are indicated on top and refer to those in **Figure S18E**. The scores were estimated using SCENIC analysis. Shown are the top 10 TFs having the highest difference in expression regulation estimates between each cluster and all other cells, tested with a Wilcoxon rank-sum test. TFs with similar activity across multiple digestive or non-digestive organ-derived clusters are outlined in the green dashed boxes.

Appendix Figure S21

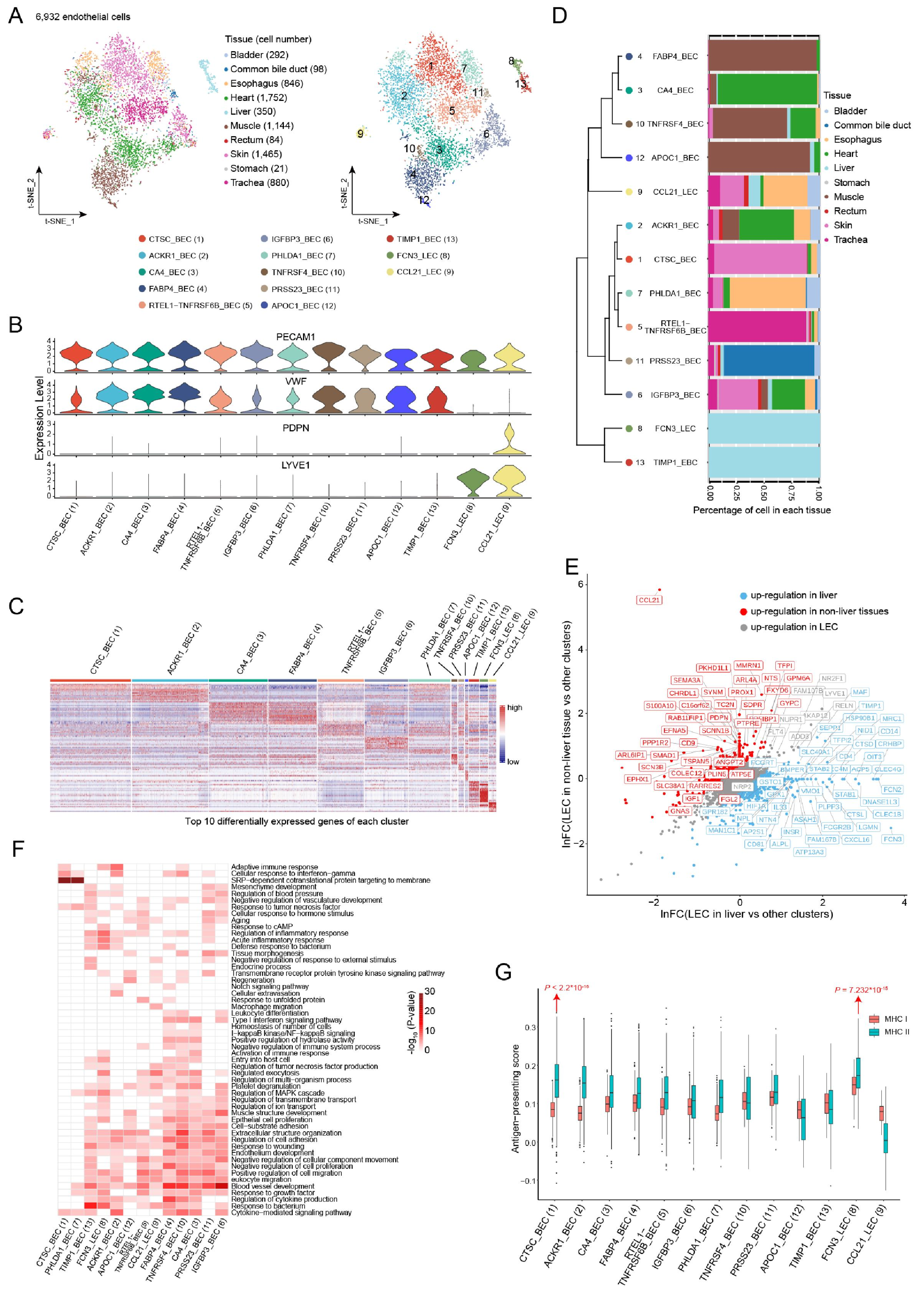

**The heterogeneous expression pattern of endothelial cells.**

**A.** t-SNE plot of 6,932 endothelial cells, colored according to tissue (left panel) or cluster (13 clusters, right panel). Each dot represents one cell.

**B.** Violin plots show the normalized expression of marker genes for pan-endothelial cells (*PECAM1*), blood endothelial cells (*VWF*), and lymphatic endothelial cells (*LYVE1* and *PDPN*) in each endothelial cell cluster as indicated at the bottom.

**C.** A scaled heat map showing the expression of the top 10 cluster-specific genes in endothelial cell clusters as indicated on top. The cluster-specific genes were defined as pct.1 ≥ 0.2 and *p.adjust* < 0.05. The expression levels of genes in lines for high and low are indicated in red and blue, respectively.

**D.** A dendrogram of cell clusters based on their normalized mean expression values of 13 clusters (correlation distance metric and complete linkage). Only genes with ln(fold-change) above 0.25, *p.adjust* < 0.05 and pct.1 ≥ 0.2 in each cluster were used to calculate the distance. Bar plots at the right illustrate the proportions of cells in the corresponding organs, colored according to tissue as in **A** (left panel).

**E.** A scatter plot showing the differential gene expression of LECs in the liver (blue) and other organs (red) compared with all other endothelial cells. Each dot represents one gene. Signature genes common for all LECs are indicated in the grey box.

**F.** Heatmap showing the GO enrichment pathways in each endothelial cell cluster. Each box represents a pathway in a cell cluster indicated on the right and at the bottom, respectively, with colors from white to red representing the significance level ($-\log_{10}P$) from low to high calculated using the online tool “metascape”.

**G.** A box plot showing the antigen-presenting scores for intra- (MHC I) and extracellular (MHC II) antigens of each endothelial cell cluster. A comparison was tested with a Wilcoxon rank-sum test. A *p-value* was calculated as in the comparison of one cluster indicated at the bottom to all other cells.

Appendix Figure S22

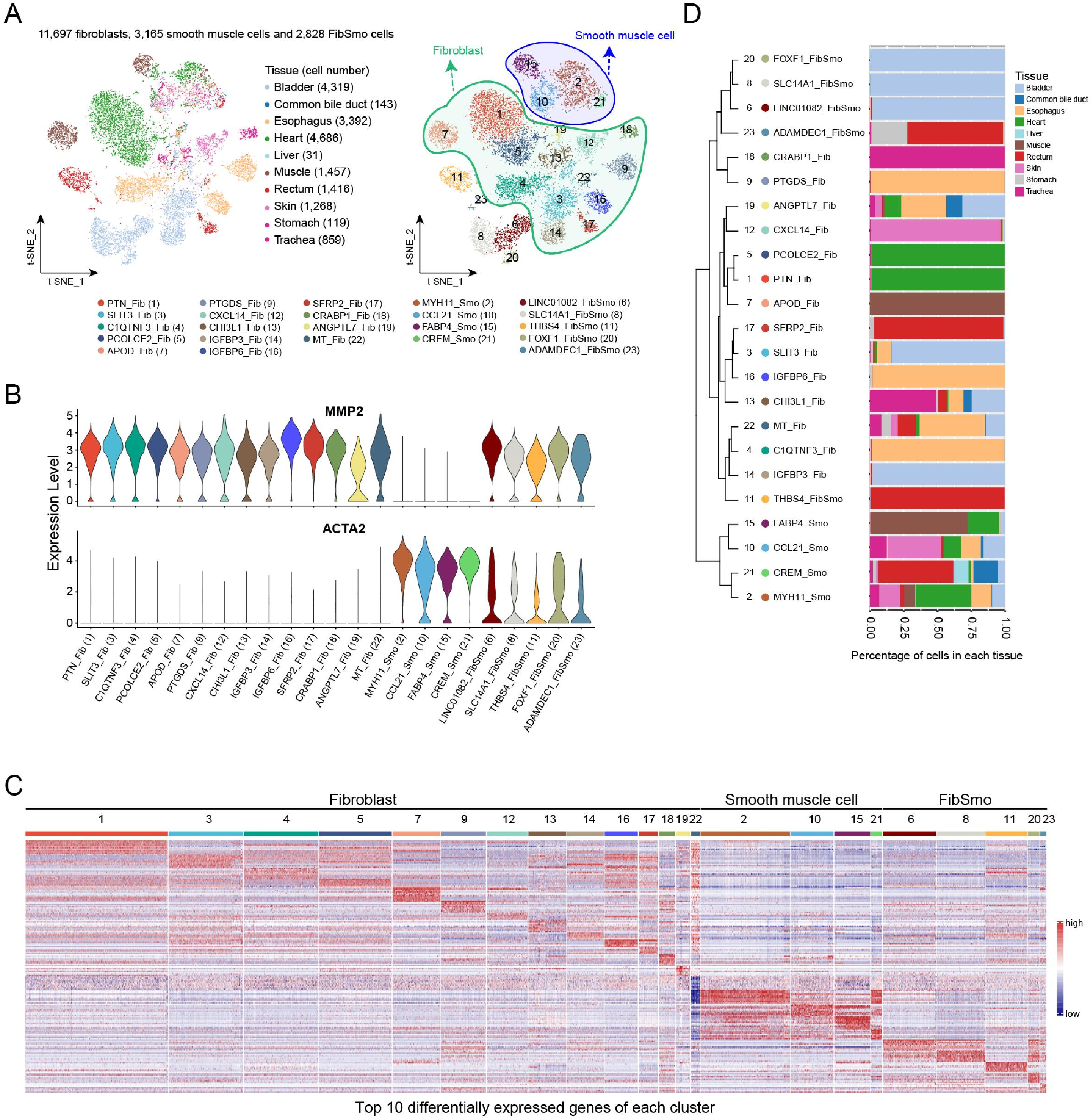


**The similarity and heterogeneity of fibroblasts and smooth muscle cells.**

**A.** t-SNE showing a total of 17,690 stromal cells, colored according to their origins of tissues (left panel) or cell clusters (23 clusters, right panel). Each dot represents one cell.

**B.** Violin plots showing the normalized expression of marker genes for fibroblasts (*MMP2*), and smooth muscle cells (*ACTA2*) in each cell cluster indicated at the bottom.

**C.** A scaled heat map showing the expression of the top 10 cluster-specific genes in each stromal cell cluster. The cell clusters are coded as numbers indicated on top, which are grouped into fibroblasts, smooth muscle cells, and FibSmo cells. The cluster numbers refer to **B.** The cluster-specific genes were defined as pct.1 ≥ 0.2 and *p.adjust* < 0.05. Each gene is shown as a line, with color heatmap from blue to red representing the expression levels for low and high, respectively.

**D.** A dendrogram of clusters based on the normalized mean expression values of 23 clusters (correlation distance metric, average linkage). Only genes with ln(fold-change) above 0.25, *p.adjust* < 0.05 and pct.1 ≥ 0.2 in each cluster are used to calculate the distance. Bar plots at the right illustrate the proportions of cells in corresponding organs, colored according to tissues as in **A** (left panel).

Appendix Figure S23

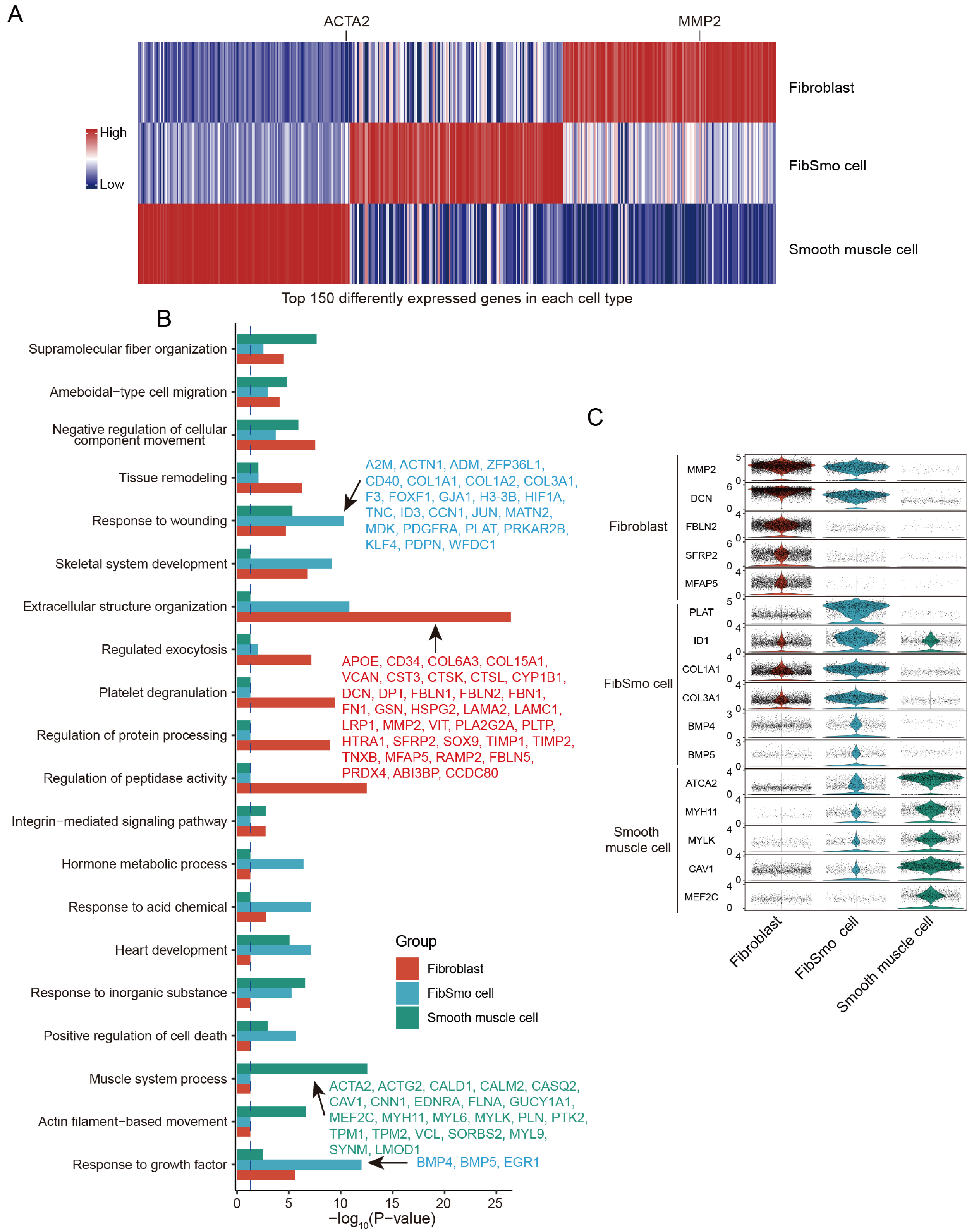


**Gene expression and biological functions of fibroblasts, smooth muscle and FibSmo cells.**

**A.** Heatmap showing the top 150 differentially expressed genes (447 genes ranked descending according to fold change) in fibroblasts, smooth muscle, and FibSmo cells. Each vertical line represents a gene, with low to high expression level color-coded in blue to red, respectively.

**B.** Bar plot showing the selected GO terms of fibroblasts, smooth muscle, and FibSmo cells. GO terms are indicated on the left and the highly enriched genes are listed in the corresponding GO term of each cell type.

**C**. A violin plot showing the normalized expression of the highly enriched gene (on the left) in fibroblasts, smooth muscle and FibSmo cells (at the bottom).

Appendix Figure S24

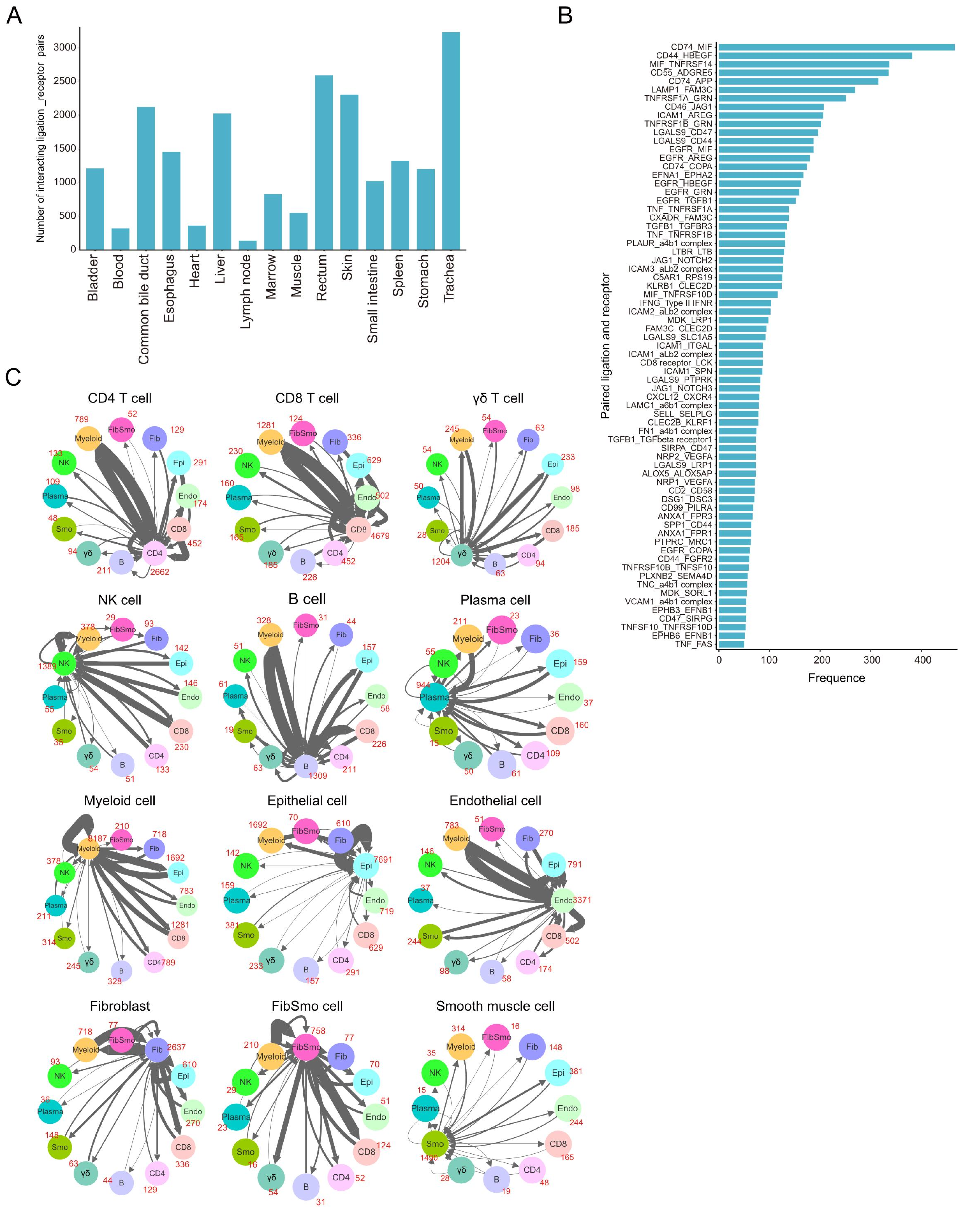


**Overview of the ligand-receptor interactions between different cell types, related to Figure 6.**

**A.** Bar plot showing the counts (on the y-axis) of significant ligand-receptor interactions in each organ indicated on the x-axis.

**B.** Bar plot showing the frequency of each ligand-receptor interaction among all cell types. Interactions with a frequency larger than 50 are shown on the y-axis, with frequencies on the x-axis.

**C.** Connection plots showing the numbers of interactions within and between a major cell type as subtitled and other major cell types as indicated in the colored circles. Numbers in red indicate the counts of ligand-receptor pairs for each intercellular link. A count of interaction between any two major cell types was a uniquely significant interacting pair between them (average expression > 0 and *P*-value < 0.05). CD4, $CD4^+$ T cell; CD8, $CD8^+$ T cell; γδ: γδ T cell; B, B cell; Plasma, plasma cell; Myeloid, myeloid cell; NK, NK cell; Epi, epithelial cell; Fib, fibroblast; Smo, smooth muscle cell; FibSmo, FibSmo cell; and Endo, endothelial cell.

Appendix Figure S25

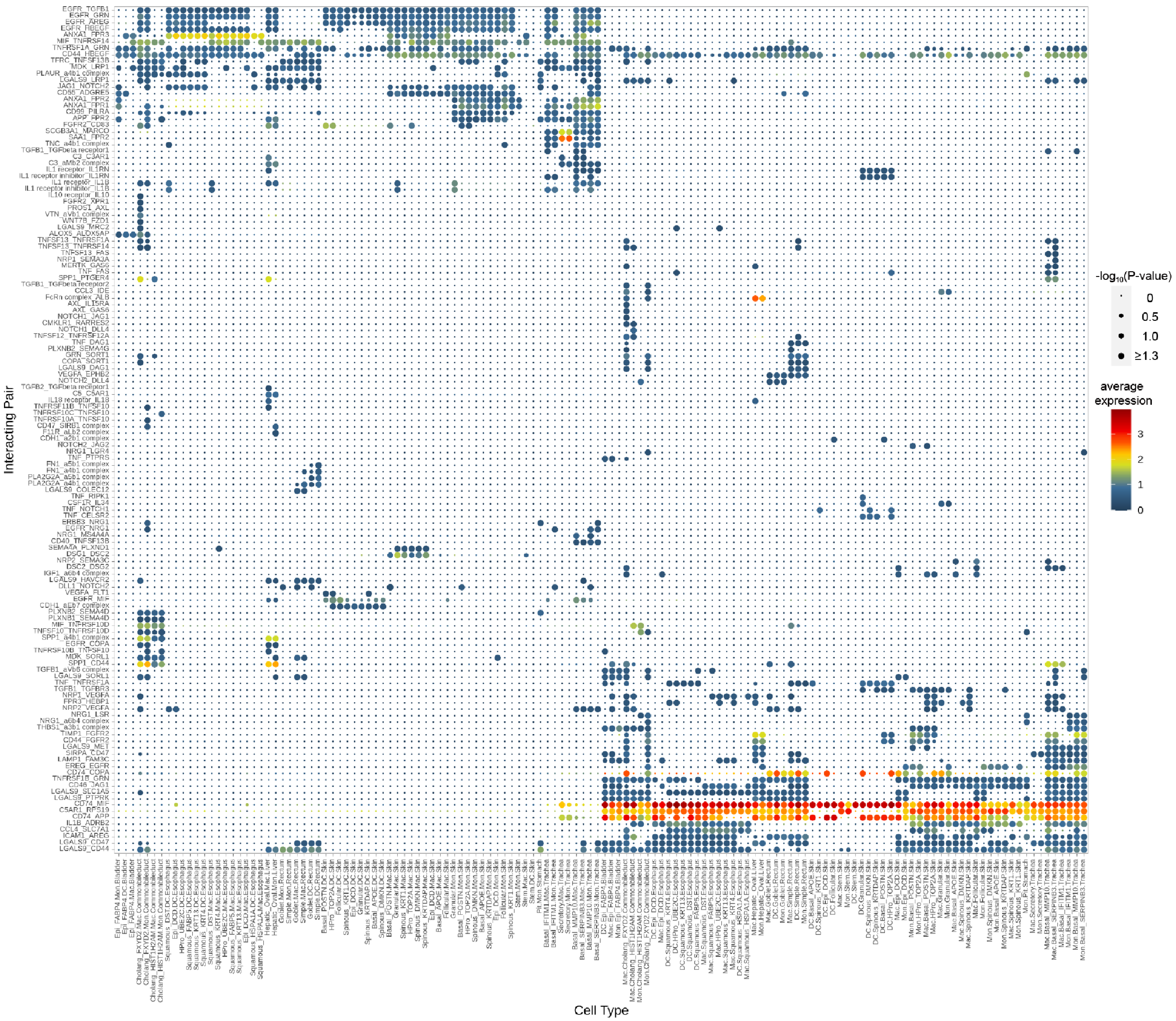


**Overview of the ligand-receptor interactions between epithelial and myeloid cells, related to Figure 6.**
Dot plot showing all the significant ligand–receptor interactions between epithelial and myeloid cell types within each organ. An interaction is indicated as color-filled circle at the cross of interacting cell types in a tissue (x-axis) and a ligand-receptor pair (y-axis), with circle size representing the significance of -$\log_{10}$(P-values) in a permutation test and colors representing the means of the average expression level of the interacting pair. The naming system is as follows, taking an example of "EGFR_TGFB1" in "cholang_FXYD2.Mac.Commonbileduct", the ligand-receptor pair is *EGFR* and *TGFB1*, and the circle is colored based on the expression levels of *EGFR* in cholang_FXYD2 cluster and *TFGB1* in Mac cluster in the tissue Commonbileduct. Mon: Monocyte, Mac: Macrophage, DC: Dendritic cell, Epi: epithelial cell. For epithelial cells, the full names of each cluster refer to **Table S47**.

Appendix Figure S26

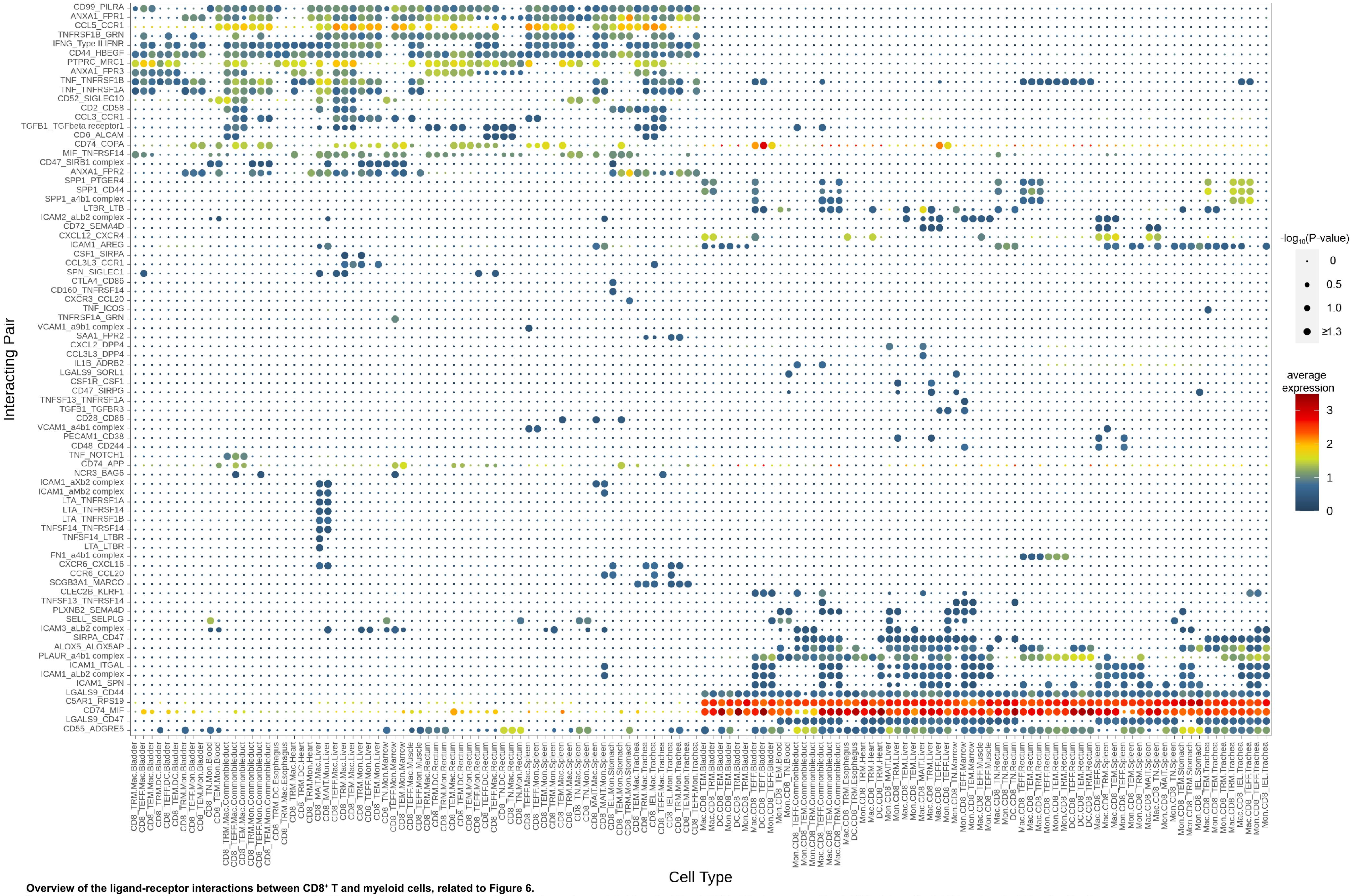


**Overview of the ligand-receptor interactions between CD8⁺ T and myeloid cells, related to Figure 6.**
Notes as in **Figure S25**. TRM: Tissue-resident memory T cell, TEFF: Effector T cell, TGD: γδ T cell, MAIT: Mucosal associated invariant T cell, TEM: Effector memory T cell, TIEL: Intraepithelial T lymphocyte, TN: naïve T cell.
esents a gene as indicated on the right; column bars represent the cells ordered by pseudotime (from left to right) and different colours represent the scaled (Z-scored) expression of each gene in each cell, with low to high levels in blue to red colours, respectively.

### Appendix Figure S27

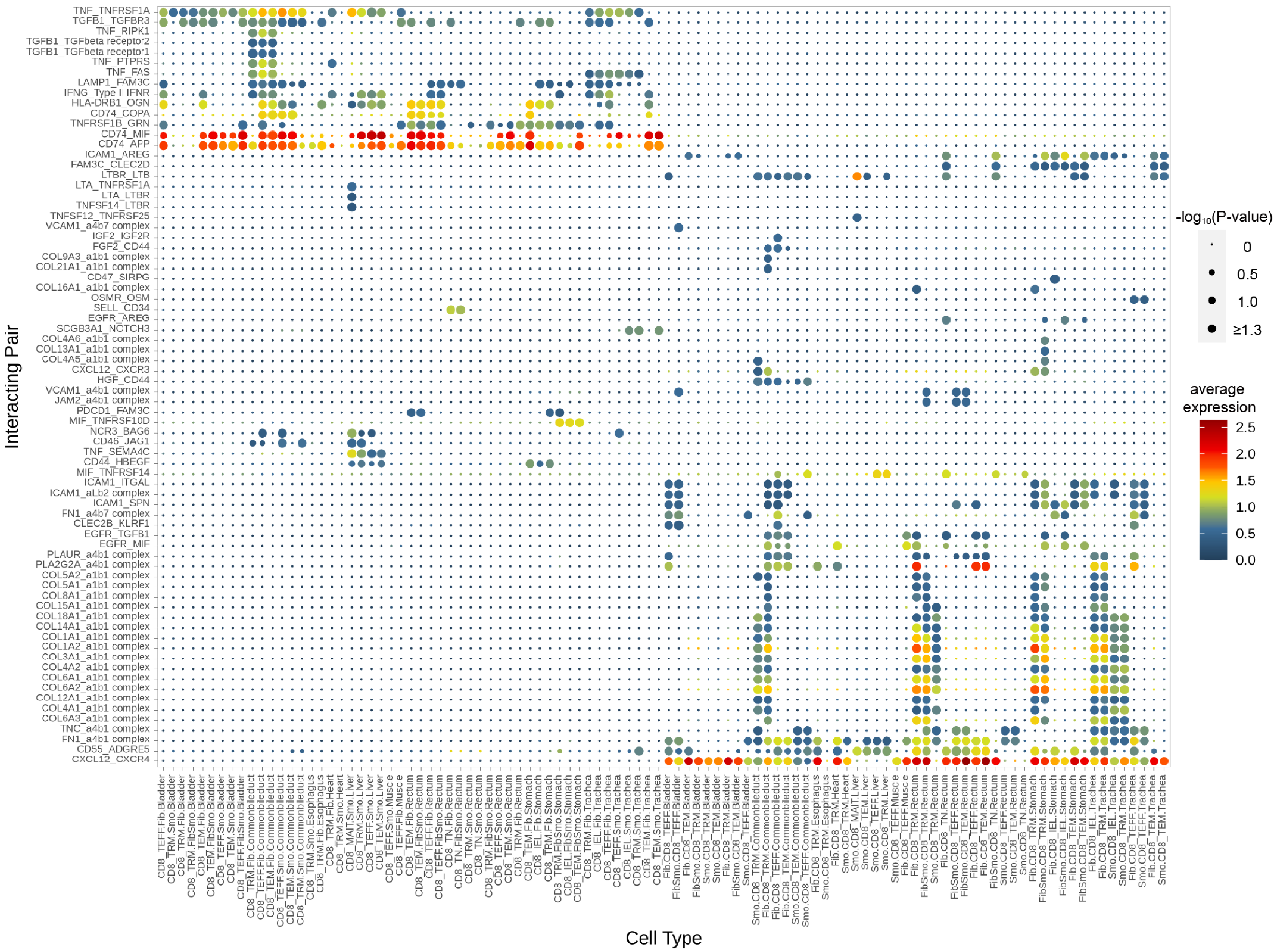


**Overview of the ligand-receptor interactions between CD8⁺ T and stromal cells (fibroblast, smooth muscle cell, and FibSmo cell), related to figure 6.** Notes as in **Figure S25** and **S26**.

Appendix Figure S28

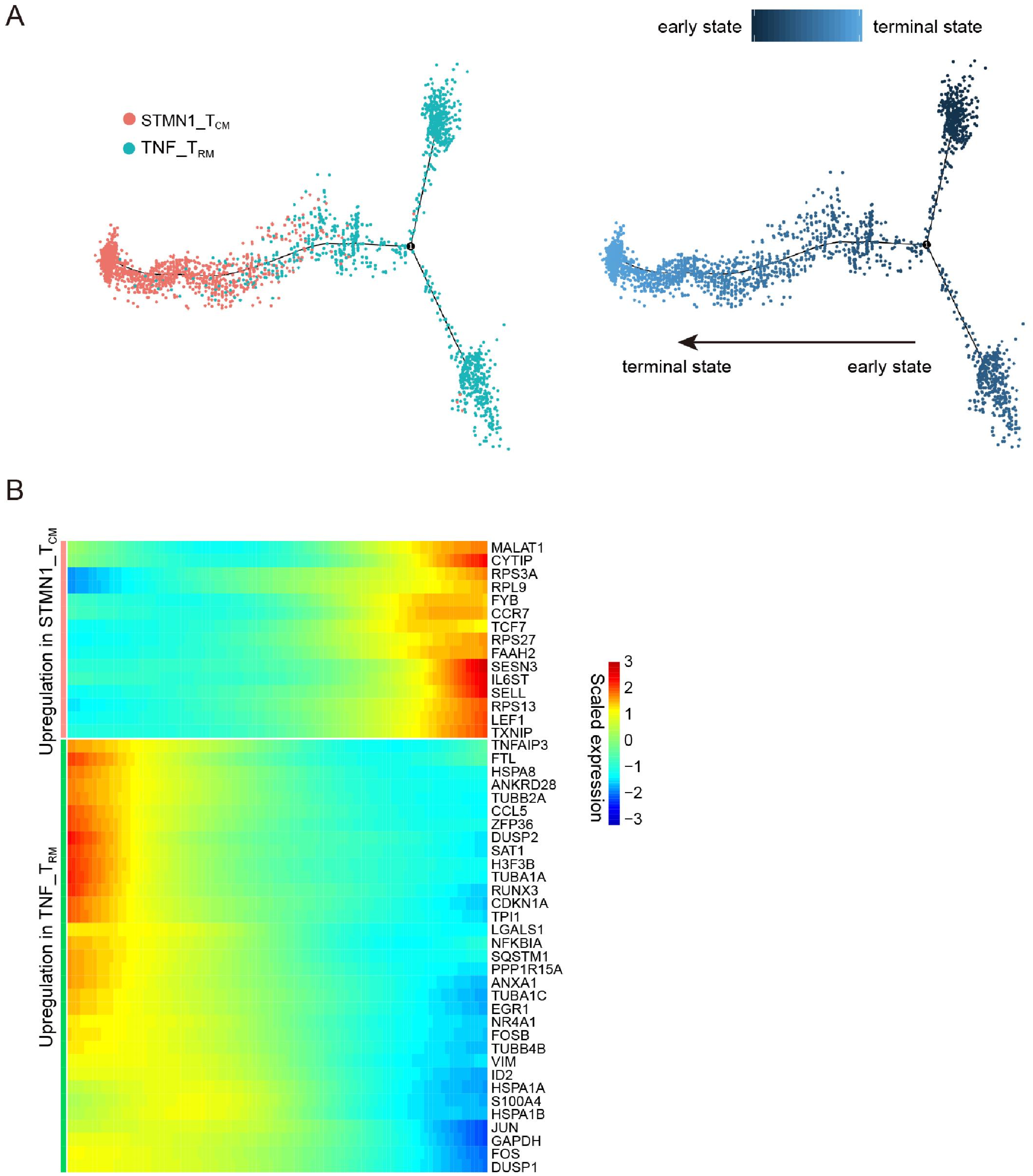


**Pseudotime trajectory analysis and gene expression profiles of TNF_$T_{RM}$ and STMN1_$T_{CM}$.**

**A.** Branched pseudotime trajectory with TNF_$T_{RM}$ and STMN1_$T_{CM}$. Each dot represents one cell, which is colored according to a cluster label (left panel) or pseudotime value (right panel) from light to dark blue representing early and terminal stages, respectively.

**B.** Expression heat map of selected variable genes along the pseudotime trajectory (from Monocle). Each row represents a gene as indicated on the right; column bars represent the cells ordered by pseudotime (from left to right) and different colours represent the scaled (Z-scored) expression of each gene in each cell, with low to high levels in blue to red colours, respectively.

Appendix Figure S29

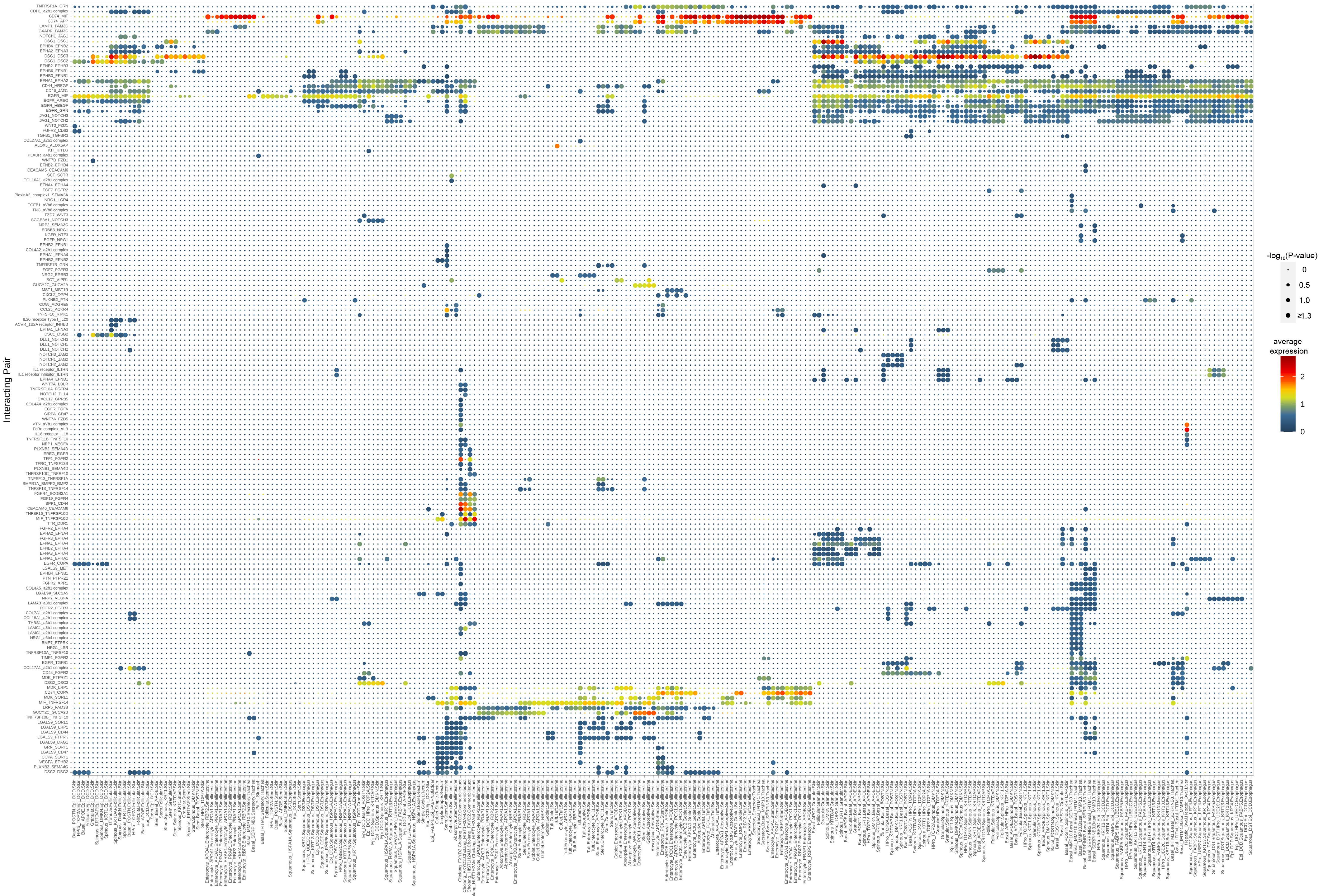


**Overview of the ligand-receptor interactions within epithelial cells.**
Notes as in **Figure S25**

Appendix Figure S30

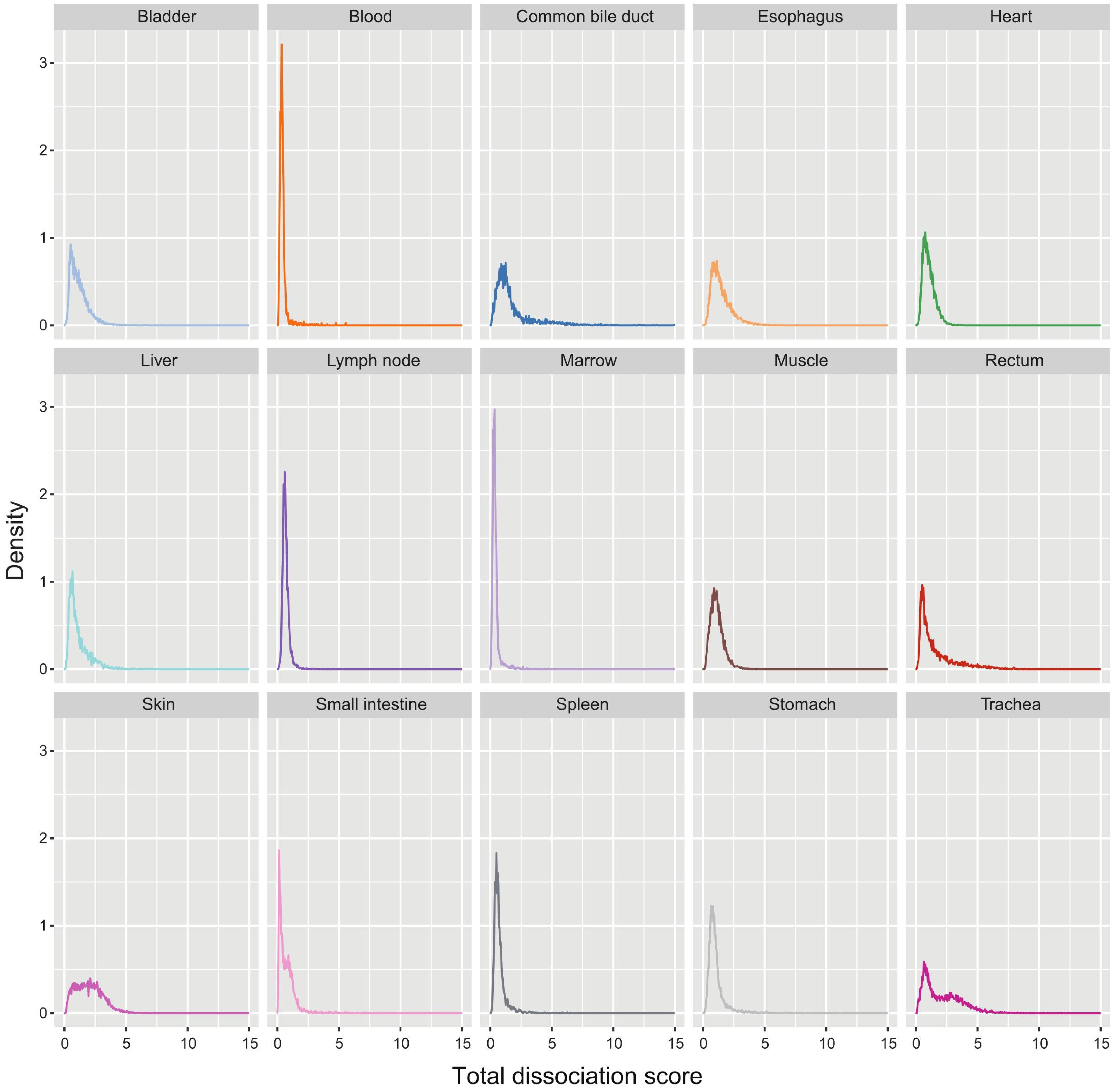


**Histogram plots of the density of dissociation scores for each organ.**
Scores at x-axis were calculated as the magnitude of the first principal component of principal component analysis based on the expression of 140 dissociation-associated genes. The y axis shows the probability density of the normalized histogram.

Appendix Figure S31

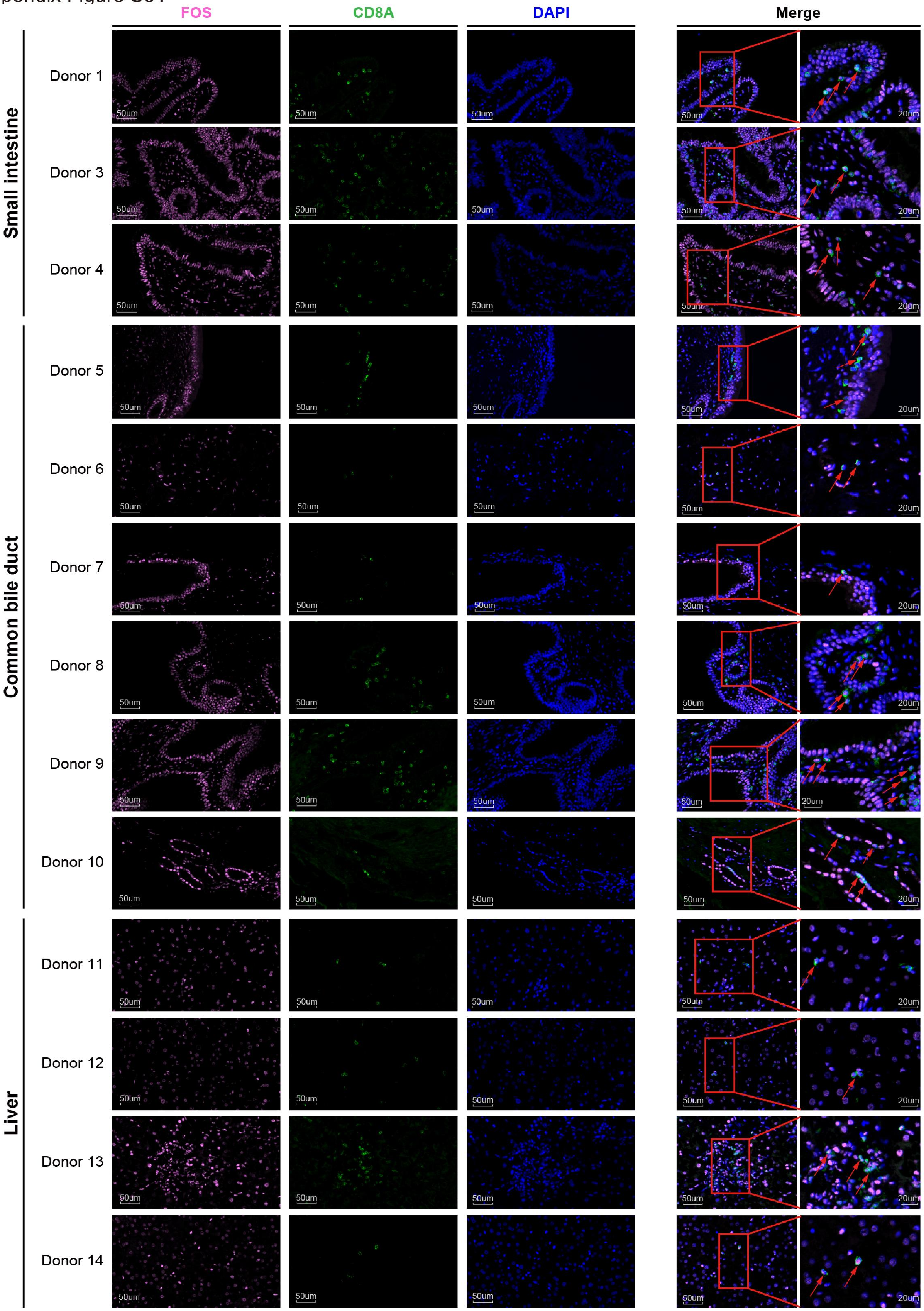


**Representative images of immunofluorescence staining of FOS and CD8A human tissue samples.**
Markers used are indicated on top. Samples of small intestine, common bile duct, and liver were derived from donors as indicated on the left. Red arrows indicate the co-expression of proteins FOS and CD8A. Scale bar is shown.
