## Supplementary Notes for "Single-cell transcriptome profiling of an adult human cell atlas of 15 major organs"

This supplementary material includes a detailed description for the validation of sweat gland epithelial cells, *COCH*^+^ fibroblasts, and FibSmo cells in existing datasets, and the evidence showing that the expression of *HSPA1A*, *FOS*, and *JUN* in the CD8^+^ T cells are unlikely stress-induced artefacts.

**1. A detailed description of validation for sweat gland epithelial cells, *COCH*^+^ fibroblasts, and FibSmo cells in existing datasets**.

**Sweat gland epithelial cells**

We validated the presence of *SCGB2A2*^+^ sweat gland cells in multiple skin samples from the two public datasets[1, 2] (**Additional file 1: Figure S4**). We observed a clear clustering of *SCGB2A2*^+^ sweat gland cells with co-expression of *DCD*, *SCGB2A2*, *PIP*, *KRT19*, and *MUCL1* genes in the skin dataset of He *et al.*’s (cluster 25; **Additional file 1: Figure S4:** left panels). A total of 346 *SCGB2A2*^+^ sweat gland cells (out of 25,326 cells in the skin) were identified in 14 skin samples in the dataset, ranging from 5 to 102 cells for each individual sample. By contrast, we only observed one *SCGB2A2*^+^ sweat gland cell with co-expression of *DCD*, *SCGB2A2*, *PIP*, *KRT19*, and *MUCL1* genes in Solé-Boldo *et al*.’s dataset (cluster 28; **Additional file 1: Figure S4A:** right panels), as limited epithelial cells (ranging from 99 to 969) were collected from each donor in the study.

**COCH^+^ fibroblasts**

We validated the presence of *COCH*^+^ fibroblasts in multiple skin and heart tissues (**Additional file 1: Figure S5**), using two skin datasets[1, 2] and one heart dataset[3] that are publicly available. We observed a clear clustering of *COCH*^+^ fibroblasts with the co-expression of *MMP2* and *COCH* genes in the skin dataset of Solé-Boldo *et al.*’s (cluster 30; **Additional file 1: Figure S5:** left top and **B:** left). A total of 144 *COCH*^+^ fibroblasts (out of 7,298 fibroblasts and 7,879 other cells in the skin) were identified in five skin samples in the dataset, with maximum 103 cells for one sample. We also observed a small group of *COCH*^+^ fibroblasts with a high expression of both *COCH* and *MMP2* in the skin dataset of He *et al.*’s (cluster 18; **Additional file 1: Figure S5A:** right and **B:** middle). A total of 125 *COCH*^+^ fibroblasts (out of 5,657 fibroblasts and 19,669 other cells in the skin) were identified in 14 skin samples in the dataset, with maximum 31 cells for one sample. By contrast, 454 *COCH*^+^ fibroblasts cells scattered among a total of 62,292 fibroblasts from the 26 heart samples in Tucker *et al*.’s dataset, with maximum 45 cells for one sample (clusters 1, 2, 4, 6, 7, 10; **Additional file 1: Figure S5A:** left bottom and **B:** right).

Next, we observed similar gene signatures of *COCH^+^* fibroblasts between our AHCA dataset and Solé-Boldo *et al.*’s dataset, compared to all other cells in the respective dataset (**Additional file 1: Figure S6A-C**). 31 out of 50 upregulated genes in *COCH^+^* fibroblasts were shared between the two datasets, among which *COCH* and *ASPN* are the two most significant ones (**Additional file 1: Figure S6A-C**). Both *COCH* and *ASPN* are secretory proteins in the extracellular matrix and have been involved in extracellular matrix remodelling [4, 5], suggesting the important role of *COCH^+^* fibroblasts in maintaining tissue homeostasis.

**FibSmo cells**

We validated the presence of *MMP2*^+^*ACTA2*^+^ cells (assigned as FibSmo cells) in multiple bladder samples from two independent public datasets[6, 7] (**Additional file 1: Figure S7A** and **B**). We observed a clear clustering of FibSmo cells with co-expression of *MMP2* and *ACTA2* genes in both datasets. A total of 1,447 FibSmo cells (out of 13,490 cells in the bladder) were identified in 3 bladder samples in the Yu *et al*.’s dataset, ranging from 42 to 709 cells for each individual sample (**Additional file 1: Figure S7A:** left panels). Similarly, a total of 178 FibSmo (out of 1,193 stromal cells in the bladder) were identified in 2 bladder samples in the Han *et al.*’s dataset, with 24 and 154 cells in each sample (**Additional file 1: Figure S7:** right panels).

**2. The expression of *HSPA1A*, *FOS*, and *JUN* in the CD8^+^ T cells are unlikely stress-induced artefacts.**

We had provided lines of evidence to show that dissociation processing in our study might have minimal impacts on the expression of *HSPA1A*, *FOS*, and *JUN* as well as the transcriptome of majority of cells in our dataset.

First, we observed dissociation-related genes were not significantly induced in the majority of cells (**Additional file 1: Figure S30**).

Second, the majority of *HSPA1A* CD8^+^ T cells (86.2%; 1,072 out of 1,244) were detected in the common bile duct, showing a tissue-specific distribution as reported previously in a mouse study[8]. The study demonstrated that *Hspa1a* was only enriched in the female reproductive tract (FRT) CD8^+^ T_RM_, but not in the spleen T_RM_ and T_CM_ cells. Moreover, although similar dissociation conditions were applied to multiple organs (**Additional file 11: Table S59**), high expression of *HSPA1A* was observed specifically in TRM_HSPA1A and a few other cell clusters (**Supplementary note Figure 1**). Supportively, exclusively high expression of *Hspa1a* was observed in only one of two T follicular helper (TFH) like cell clusters but no other T cells types in the lung of mouse[9]. These observations suggest *HSPA1A* to be one of the specific markers for T cell subtypes.

Third, we suspected that high expression of *FOS* and *JUN* in CD8^+^ T_RM_ populations (**Supplementary note Figure 1**) might reflect their role in maintaining the cells rather than in response to the dissociation-related stress. *FOS* and *JUN* are members of AP-1 transcription factor, of which dimerization has been shown to play numerous roles in a wide range of cellular process, cell growth, and proliferation, etc. We did observe the upregulation and the high regulon’s activity of many AP-1 dimerization partners (such as *FOS*, *JUN*, *JUNB*, *FOSL2*, and et) for CD4^+^ (**Fig. 2C**) and CD8^+^ T_RM_ cells (**Fig. 2D**) in our dataset. It has been reported recently that AP-1 dimerization partners such as *Fosl2* and *Junb* were highly expressed in T_RM_ cells, and the knockdown of *Junb* resulted in impaired T_RM_ cell differentiation, confirming the key role of AP-1 dimerization regulating T_RM_ differentiation[10]. Given that *Fosl2* is able to regulate *Smad3*[11], a key component of the TGF*β* signalling pathway, *Fosl2* may also promote T_RM_ cell differentiation through effects on TGF*β* signalling. We note that high expression of *FOS* and *JUN* was only observed in most of T_RM_ cells, but not non-T_RM_ cells (**Supplementary note Figure 1**) from the same organs that were undergone dissociation treatments with similar conditions. This observation, together with our previous observation that most organs had higher density of low total dissociation score (**Additional file 1: Figure S30**), suggests that dissociation-related genes were not significantly induced in the majority of cells.

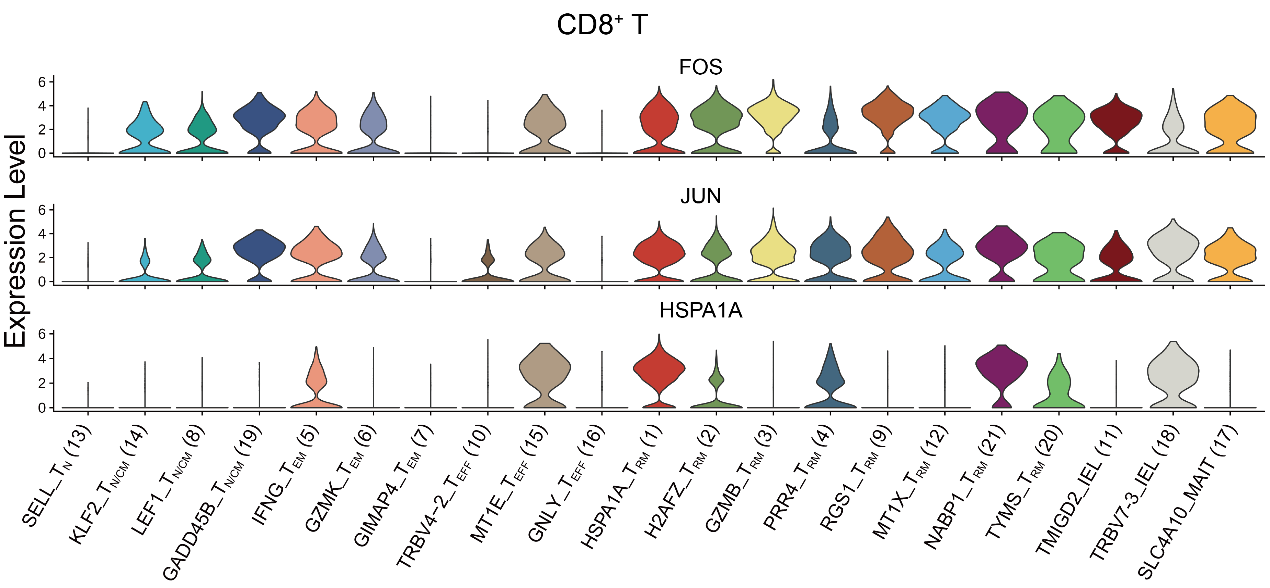

**Supplementary note Figure 1. The violin plots showed the normalized expression of *FOS*, *JUN*, and *HSPA1A* in CD8^+^ T cell clusters.**

Lastly and importantly, additional immunofluorescence assay with double staining of CD8A and FOS in common bile duct, liver, and small intestine from multiple donors confirmed the high expression of FOS in the CD8A cells before tissue dissociation (**Additional file 1: Figure S31**).

In summary, these observations suggest that the expression of *HSPA1A*, and *FOS*/*JUN* in the cell clusters in our present study are unlikely stress-induced artefacts.
